# Revealing centurial changes in butterfly wing shape via the automated analysis of museum specimens

**DOI:** 10.64898/2026.01.09.698621

**Authors:** Mahika K. Dixit, Richard J. Gill, William D. Pearse

## Abstract

1. Understanding how environmental change is shaping organismal morphology of key functional guilds is important, since morphological traits determine the delivery of ecosystem services we depend upon. Adult butterflies, as pollinators, are commercially and culturally important, but the longitudinal data across appropriate time frames to understand trait response trends are generally limited.
2. Here we leverage digitised museum butterfly specimens to help fill this data gap by employing automated methods (machine learning and Elliptical Fourier Analysis) to achieve high-throughput measurements of traits across large collections. We analyse how wing shape (the outline of the whole butterfly) varies in over 178,000 specimens of 65 butterfly species collected across the UK and Ireland over two centuries.
3. We identify two major biologically attributable axes of butterfly wing shape: aspect ratio and asymmetry. Analysing a subset of these specimens (108,511 from 34 species), we find directional trends in wing shape. Over a period of 105 years (1901-2006), wings generally became narrower and longer in over half (18) of the studied species. This response may be affecting butterfly behaviour by favouring gliding flight.
4. Ultimately, we highlight that historical processes including temperature change have historically shaped butterfly wing shape. We demonstrate that studying whole wing outline using automated methods allows us to extract the high-throughput, long-term data required to understand the responses of ecologically important taxa at scales that we were previously unable to achieve.

## Introduction

High rates of climate and land-use change over the past few decades have placed many pressures on insect species (Vanbergen et al., 2013; Wagner, 2020). Such environmentally driven selective pressures can reshape the functional trait space that populations represent, affecting the ways organisms respond to their environment and contribute to ecosystem functioning (Díaz et al., 2013). Ultimately, this can have complex effects on ecosystem service provision (Díaz et al., 2013). For insect pollinators, wing shape is one trait that can determine their functional roles and underpin fitness given its importance in governing flight and thermal basking (Bladon et al., 2020; Le Roy, Debat & Llaurens, 2019; Wootton, 1992). Investigating variation in wing shape is thus particularly interesting, especially in butterflies which have the largest wing area to body mass ratio of flying insects and two highly developed wing pairs (Le Roy, Debat & Llaurens, 2019). Previous work on butterfly wings has demonstrated that both historical processes and contemporary selection can shape the wing colour patterns and size of wings in UK butterflies (Dennis & Shreeve, 1989). Yet, we know far less about how environmental change over the last century has impacted wing shape or variance in wing shape, which can reveal signatures of developmental stress, environmental filtering, plastic trait change, and trait evolution (Arce et al., 2023; Sanderson et al., 2023).

Butterflies utilise both energy-intensive flapping flight and low-energy gliding flight. These behaviours are favoured by different morphological characteristics: short, broad wings with a low aspect ratio are more aerodynamically effective for flapping flight, while long, narrow wings with a high aspect ratio favour gliding flight. It has been suggested that butterfly species trade-off between these two strategies (reviewed in: Cespedes, Penz & DeVries, 2015; Le Roy, Debat & Llaurens, 2019). For instance, butterflies with high agility and manoeuvrability are generally associated with large, low aspect-ratio wings relative to body mass and favoured in more spatially cluttered environments, but these traits are traded off against low energy expenditure, which is favoured by high aspect-ratio wings (Cespedes, Penz & DeVries, 2015; DeVries, Penz & Hill, 2010; Le Roy, Debat & Llaurens, 2019). Temperature may also impact wing shape: for example, different ecotypes of *Coenonympha oedippus* (false ringlet butterfly), which inhabit different microclimates, significantly differ in their wing shape (Jugovic et al., 2018). Increased migration success in *Danaus plexippus* (monarch butterfly) is associated with longer, larger, and less rounded wings with a higher aspect ratio (Altizer & Davis, 2009; Dockx, 2007; Flockhart et al., 2017; Satterfield & Davis, 2014). Wamer environmental temperatures are linked to increased energy expenditure during migration, and so may also select for wings with a higher aspect ratio (Parlin et al., 2023). Together, this highlights how disentangling the environmental factors impacting wing shape in UK butterfly populations is key to understanding the pressures impacting butterflies and pollination services.

Wing symmetry is another key aspect of wing shape. Non-random deviation from symmetry biased to the left or right wing, known as *directional asymmetry*, is considered adaptive (Windig & Nylin, 1999). Conversely, random small deviation from perfect bilateral wing symmetry (Rossato et al., 2018), known as *fluctuating asymmetry*, is thought to be a proxy of developmental stress (Badyaev, Foresman & Fernandes, 2000; Gerard et al., 2018; Palmer & Strobeck, 1986). Understanding how wing symmetry changes over time or with environmental change may reveal contributors to developmental stress in butterflies. Previous work has linked fluctuating asymmetry to certain environmental conditions (Arce et al., 2023; Leary & Allendorf, 1989; Lens, Van Dongen & Matthysen, 2002; Parsons, 1992; Pellegroms et al., 2009) and genetic processes (e.g. loss of genetic variation, hybridisation; see Leary & Allendorf, 1989) in many taxa (Adamski & Witkowski, 2003; Leary & Allendorf, 1989; Lens et al., 2000; Parsons, 1992). Despite this, some studies find no link to environmental stress (Gerard et al., 2018; Hoffmann, Collins & Woods, 2002). Fluctuating asymmetry is thought to be linked to fitness, though this is hard to test (Lens, Van Dongen & Matthysen, 2002), and evidence is inconsistent (Graham et al., 2010). Understanding the relationships between increased temperatures and fluctuating asymmetry in butterflies may thus help identify stressed populations.

To understand morphological trends, studies are often limited by low taxonomic resolution, geographic location, and temporal representation (Ashe-Jepson et al., 2024), tending to focus on single functional traits (e.g., Hartfelder et al., 2020) and single species (e.g. Hoffmann, Collins & Woods, 2002). This is because manually extracting large volumes of complex functional trait information is time-consuming, expensive, and difficult given a lack of available data (Weeks et al., 2022). This limits our understanding of both the general ecology of butterflies, as well as the provision of ecosystem services. Digitised museum collections represent an under-utilised source of long-term functional trait data, helping to overcome any potential shifting baselines of traits (Bartomeus et al., 2019). Museums hold large collections of butterflies, some of which have been digitised (Bartomeus et al., 2019; Kharouba et al., 2019), making the long-term study of traits like wing shape possible (Wilson et al., 2022). These large datasets are ideal for processing using sophisticated computational techniques including machine learning, and the scale of these data mean we need to use automated methods such as Elliptical Fourier Analysis to quantify shape. Using automated methods to extract morphological information from these images allows datasets to be compiled in a fraction of the time without human error in shape estimation (Weeks et al., 2022). Since Elliptical Fourier Analysis considers whole outlines (Bonhomme et al., 2014), this method may give a more holistic perspective of the relationship between environmental change and butterfly wing shape. In this study we use automated methods to analyse over two centuries of UK butterfly museum specimens from the NHM London. We investigate how wing shape varies within and across species, quantifying trends in wing shape over time and with changing environmental temperature.

## Methods

Using images of butterfly specimens from the Natural History Museum London (NHM) Lepidoptera collection, we build on an existing ML pipeline, ‘mothra’ (Wilson et al., 2022), to output the wing outline of each butterfly specimen. Using Elliptical Fourier Analysis, we then quantified wing shape for each specimen. We performed a PCA to identify the major axes of shape variation and fitted linear and quantile regression models to determine how wing shape and asymmetry varied across species, time, and environmental temperature.

### Data

The high-resolution NHM butterfly specimen images (Paterson et al., 2016a, 2016b) include the specimen photographed on a grey background with any available labels. All associated label metadata for each specimen was also digitised and sourced from the NHM portal (Scott et al., 2019). Since *Inachis io* is also known as *Aglais io*, the metadata for these species were combined. Due to differences in nomenclature between the NHM database and the literature, we used the search term *Ochlodes sylvanus* for *Ochlodes venata* and *Fabriciana adippe* for *Argynnis adippe* to access the relevant metadata. Species were grouped into families; where possible, family information was assigned using a recent phylogeny (Kawahara et al., 2023), and otherwise the GBIF taxonomic classification was used (GBIF.org, 2025). Though *Leptidea sinapis* was recently found to encompass three cryptic species (Dincă et al., 2011), since morphological differentiation of these species is not reliable, we utilised the NHM species classification for consistency.

The final dataset used to understand variation in butterfly wing shape consisted of 178,359 specimens of 65 species (including migrant species) of UK butterfly. These data spanned the UK and Ireland but were mainly concentrated in the south of England (Fig. 1). There were 108,516 specimens of 34 species that had sufficient records for modelling (i.e. each species had >1000 observations), available metadata of suitable quality (i.e. included year and coordinate location of collection), and available temperature data (available post-1901). Five of these specimens were removed from the dataset for data analysis due to atypical outlines. Two specimens appeared to have incorrectly recorded location coordinates but did not meet the requirements to be included in data analysis regardless (Fig. 1).

**Fig. 1.**
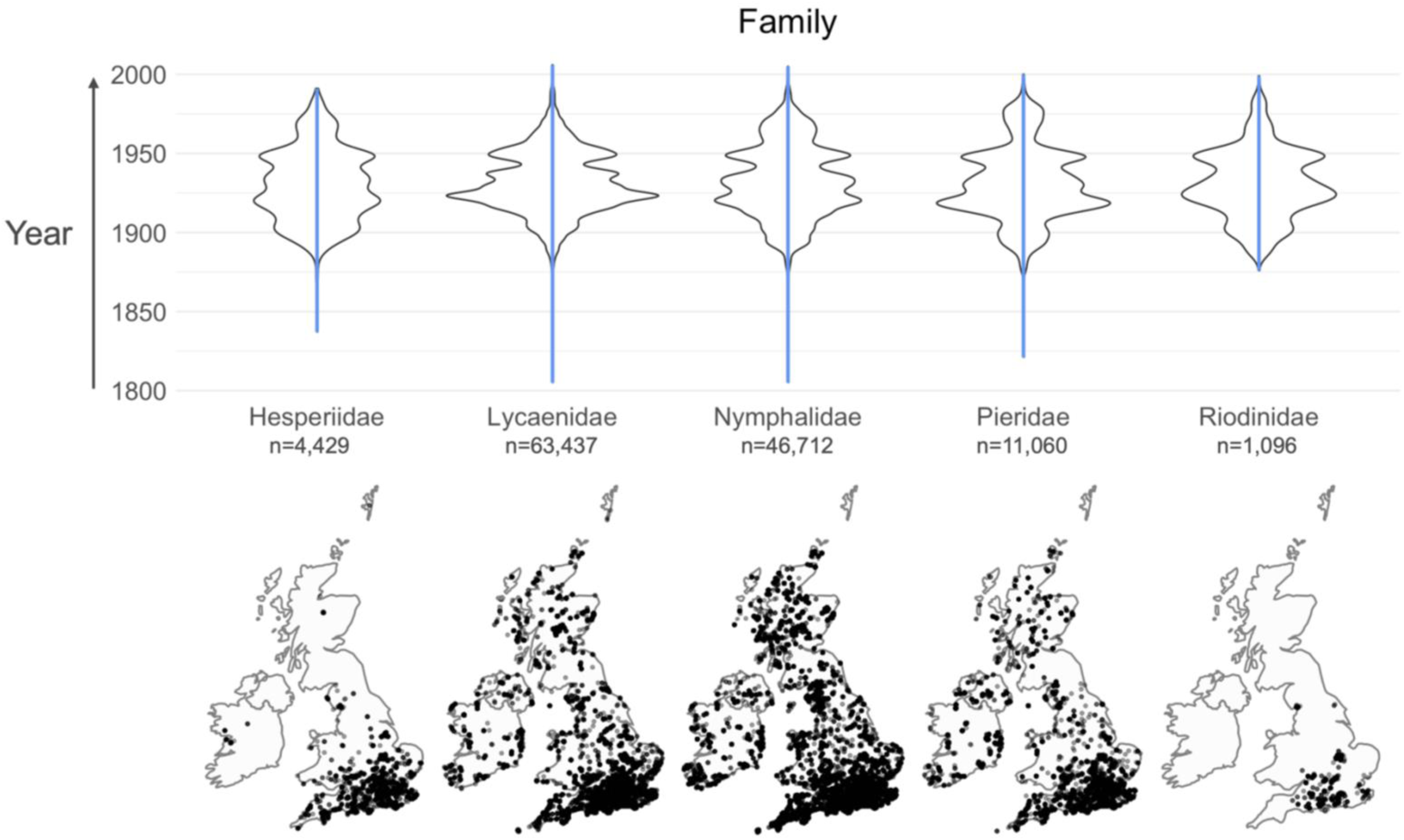
The digitised museum specimens used in this study had large temporal and spatial representation, covering the UK and Ireland across over two centuries (1805-2006) in three families (Lycaenidae, Nymphalidae, and Pieridae). Above, violin density plots grouped by family describe the variation in the year of specimen collection, with the range of years shown using a blue vertical line. Below, spatial distribution plots, grouped by family, show the coordinate locations where these specimens were collected. Here, each black point represents a single specimen. This figure represents the specimens (n=126,734) with sufficient associated metadata (including year and coordinate location of collection). The Riodinidae family, which was represented by a single species (Hamearis lucina), was collected in a relatively restricted area, but encompassed specimens collected across over a century. The family Papilionidae (which was represented by the single species Papilio machaon in the morphospace) is excluded from this plot, since there were fewer 1000 observations of this species in the dataset (see Methods).

We calculated the mean annual temperature in the year of lethal collection for each specimen with an available coordinate location and year of collection using data (CRU TS v. 4.04) from the UEA Climatic Research Unit (Harris et al., 2020) using *rdgal*, *raster*, and *ncdf4* (Bivand, Keitt & Rowlingson, 2021; Hijmans, 2022; Pierce, 2021).

### Machine learning pipeline

To segment (separate) the butterflies in the specimen images from their background, we extended the existing machine learning (ML) algorithm ‘mothra’ (Wilson et al., 2022), which was trained on the NHM Lepidoptera collections. Mothra corrects rotated images, detects the ruler in each image, scales the image, and crops out any labels. Identifying the unique regions in the image, it converts the original image into a binary image which can be outputted. The algorithm then identifies specific points on the butterfly and measures the distances between them. We modified mothra to output only cropped black and white binary images with each butterfly in the centre of the image (Fig. 2) for downstream Elliptical Fourier Analysis (see below).

**Fig. 2:**
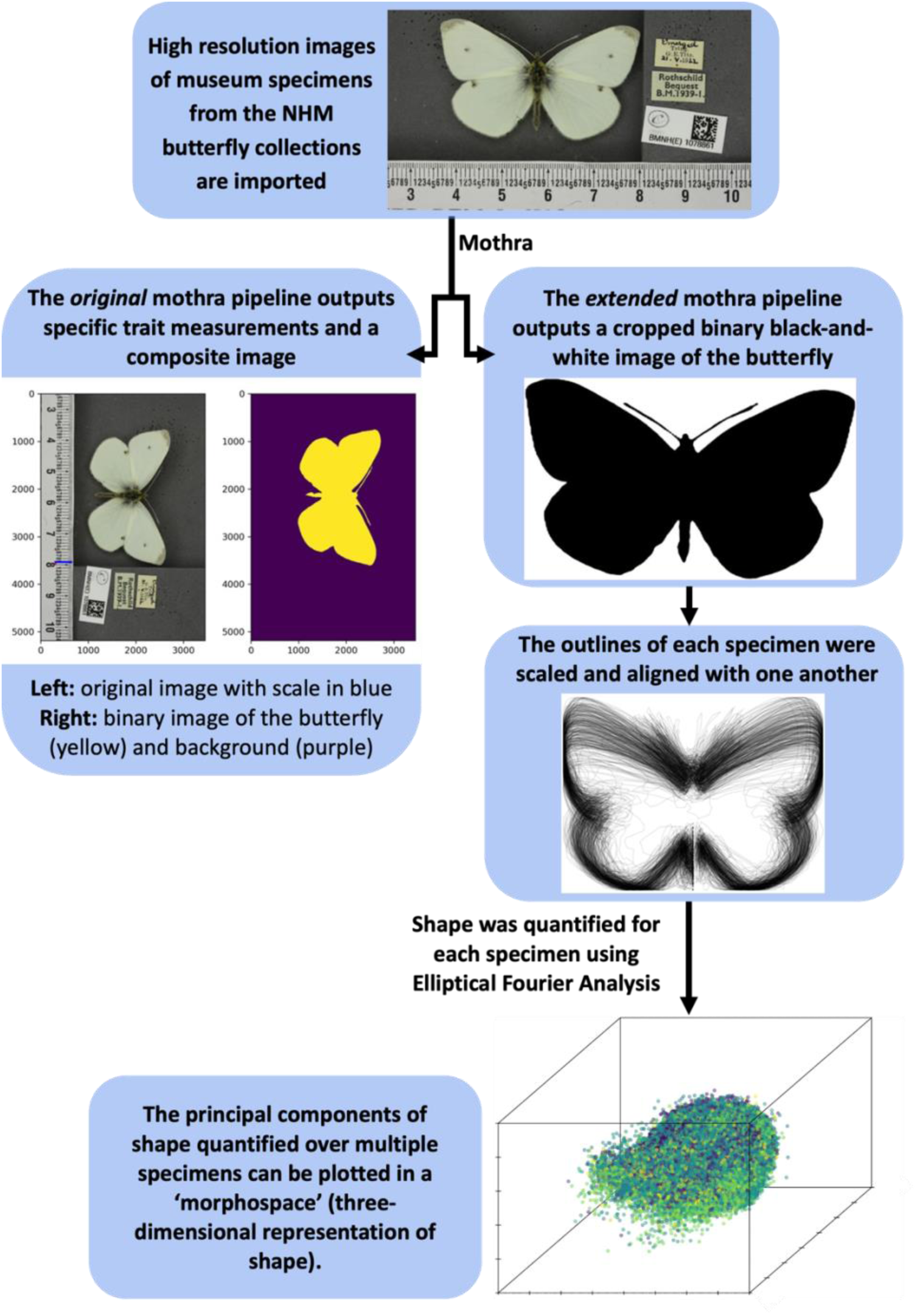
A schematic illustrating pipeline used in this study. We extended the machine learning pipeline ‘mothra’ to output binary images of butterfly specimens. Using Momocs in R, we imported the images outputted from our extended mothra pipeline, filtered out images with poor quality outlines, then scaled and aligned the specimens using a full generalised Procrustes alignment. After performing the Elliptical Fourier Analysis, we performed a Principal Component Analysis and used the output to plot a ‘morphospace’. An image of the species Pieris rapae is used to illustrate the input data and results of the original and modified Mothra pipelines. A small subset of specimens was used to produce the stacked output illustrating the Procrustes alignment.

### Extracting wing shape

We first filtered out any species with fewer than 50 specimens in the collections, to avoid including rare immigrants, reared butterflies, and very rare species. We then used the remaining images as input data for the ML pipeline. Importing the binary image output from the ML pipeline into R, we filtered out 12 images with fewer than 3,000 coordinates defining the specimen outline to ensure all retained images were of good quality. This left us with a dataset of 178,359 images from 65 species. We then scaled and aligned the specimens using a full generalised Procrustes alignment. For this, we split the images (n=178,359) into two batches (Batch 1: images 1 to 89,180; Batch 2: images 89,181–178,359), since aligning in one batch exceeded the available computing resources. We visually compared the stacked outlines of these two batches to ensure that this approach did not affect the alignment.

We then carried out Elliptical Fourier Analysis using *Momocs* (Bonhomme et al., 2014) in R. Elliptical Fourier Analysis approximates the outline of a shape using a finite series of trigonometric terms, termed harmonics (Bonhomme et al., 2014; Carlo, Barbeitos & Lasker, 2016). Here, lower frequency harmonics approximate coarser-scale, and higher frequency harmonics finer-scale, shape variation (Bonhomme et al., 2014). These methods must therefore balance demands of computational power (which increase with the number of harmonics computed) while effectively describing fine-scale shape variation. Uniquely, Elliptical Fourier methods allow almost any outline to be quantified and can produce harmonic coefficients (four of which are calculated per harmonic) independent of outline position and size (review: Bonhomme et al., 2014). Therefore, Elliptical Fourier Analysis is particularly suitable for quantifying shape in butterflies, since their shape is complex and species display significant size differences.

Following methods described in Rubin *et al*. (2018), we elected to use 12 harmonics, which captured 99.9% of the total harmonic power. Carrying out Elliptical Fourier Analysis and then a Principal Component Analysis in *Momocs*, we generated a three-dimensional morphospace of butterfly shape including all 178,359 images from the 65 species with sufficient data (Fig. 4). We extracted values for the first three Principal Components of the PCA since these axes explained the majority of variation in the data. To compare the amount of variation in wing aspect ratio (corresponding to PC1; see results) and wing asymmetry (corresponding to absolute PC2 value; see results) explained by species and family identity, we carried out a variance composition analysis using *lme4* (Bates et al., 2015).

We calculated symmetry in *Momocs* as the sum of the absolute values of harmonic coefficients A and D (which represent symmetrical variation for the central axis of each specimen) over the sum of the absolute value of all harmonic coefficients (Bonhomme et al., 2014; Yoshioka et al., 2004). We calculated correlation coefficients between these symmetry values and the three PC axes.

### Data analysis

Since the specific element of wing shape represented by PC3 was unclear, it was excluded from modelling in this study. We constructed linear models to determine the impact of the predictor variables *year:species* (an interaction between species and year), *temperature:species* (an interaction between species and temperature), and *species* on wing aspect ratio (corresponding to PC1; see results) and wing asymmetry (corresponding to absolute PC2 value; see results). The variables *year* and *temperature* (mean annual temperature in the year of collection) were scaled and centred (z-transformed) such that model coefficients represented standardised effect sizes, allowing direct comparison of effect sizes (Gelman, 2008). In our wing aspect ratio model, the response variable was ‘*scaled and centred PC1 value*’ – here a **positive** trend in ‘*scaled and centred PC1 value*’ represents a **decrease** in wing aspect ratio. In our wing asymmetry model, we used the response variable ‘*scaled and centred absolute PC2 value*’, since only the absolute PC2 value was strongly correlated with wing symmetry (Fig. 4). In both cases we removed the model reference intercept, overall temperature, and overall year terms to aid interpretation, since we were interested in determining whether species’ means significantly differed from zero, as opposed to a reference level. In accordance with Schielzeth (2010), we did not report the results from the F test, since this approach to model formation renders the test unsuitable. However, this approach allowed us to directly output the standard errors for each species’ slope (Schielzeth, 2010), and thus we used the model p-values to determine whether species’ slopes significantly differed from zero. For both models, 108,511 images of 34 species, lethally collected between 1901-2006, were used.

We also constructed equivalent quantile regression models using *quantreg* (Koenker, 2024). We chose τ values (representing the quantiles of interest) of 0.1, 0.25, 0.5, 0.75, and 0.9 to compare model results between subsets of the data representing specimens with extreme trait values and specimens with moderate or median trait values. This approach also allows trends in the variance of response variables to be understood if the direction or strength of trends differs between high and low quantiles (Fig. 3). For example, a positive trend at higher quantiles and negative trend at lower quantiles would be indicative of increasing variance. Similarly, a stronger positive trend at higher quantiles than at lower quantiles would be indicative of increasing variance. Both quantile regression models were fitted using a sparse implementation of the Frisch-Newton interior-point algorithm due to the size of the dataset.

**Fig. 3:**
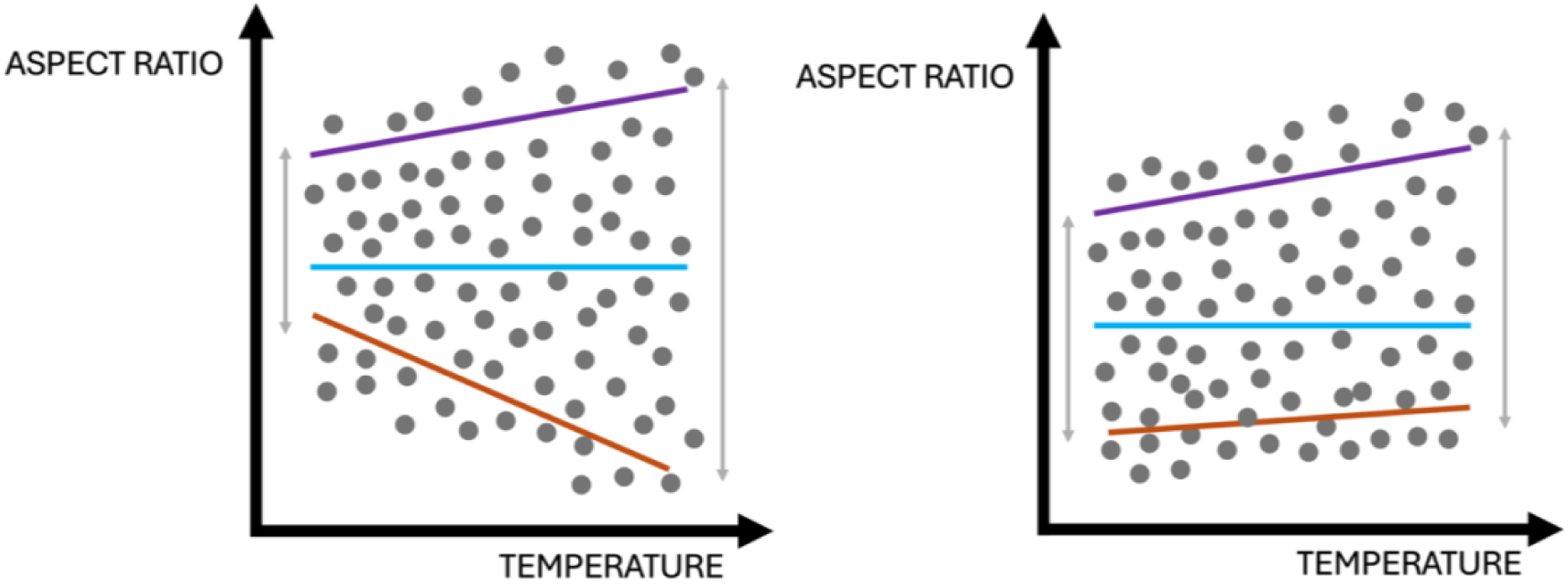
Quantile regression models can be used to understand changes in variance in a response variable (e.g. wing aspect ratio) with an explanatory variable (e.g. temperature). This example depicts two scenarios where at low temperature values there is less variation in aspect ratio than at higher temperatures. Here, trend lines in the linear model are shown in blue, trends for higher quantiles in purple, and for lower quantiles in orange. A positive trend at higher quantiles and negative trend at lower quantiles (left) and a stronger positive trend at higher quantiles than at lower quantiles (right) are indicative of increasing variance (where variance is depicted using grey arrows).

All plots were generated using *mapproj*, *mapdata*, *ggplot2*, *viridis*, *patchwork*, *Momocs*, and *scatterplot3d* (Becker, Wilks & Brownrigg, 2022; McIlroy, Brownrigg & Minka, 2023; Garnier et al., 2024; Pedersen, 2024; Bonhomme et al., 2014; Ligges & Mächler, 2003) in R (R Core Team, 2024).

## Results

### Morphospace

Our pipeline successfully extracted wing shape from 178,359 images of butterfly museum specimens, though Mothra was unable to separate the antenna from the wings during image segmentation in some cases (see Fig. 4). PC1, representing 46.2% of variation in the dataset, corresponded to wing aspect ratio (Fig. 4), forming an axis of wing shape from long, narrow wings (high aspect ratio; low PC1 values) to short, broad wings (low aspect ratio; high PC1 values). PC2 (11.8%) corresponded to wing asymmetry, where higher absolute PC2 values – i.e. higher deviation from the mean – explained higher wing asymmetry. Low PC2 values were indicative of deviations in the right wing, while high PC2 values were indicative of deviations in the left wing. PC3 (8.6%) was difficult to attribute to specific aspects of shape. Low PC3 values, which were rare, may be associated with poorly segmented specimens.

**Fig. 4:**
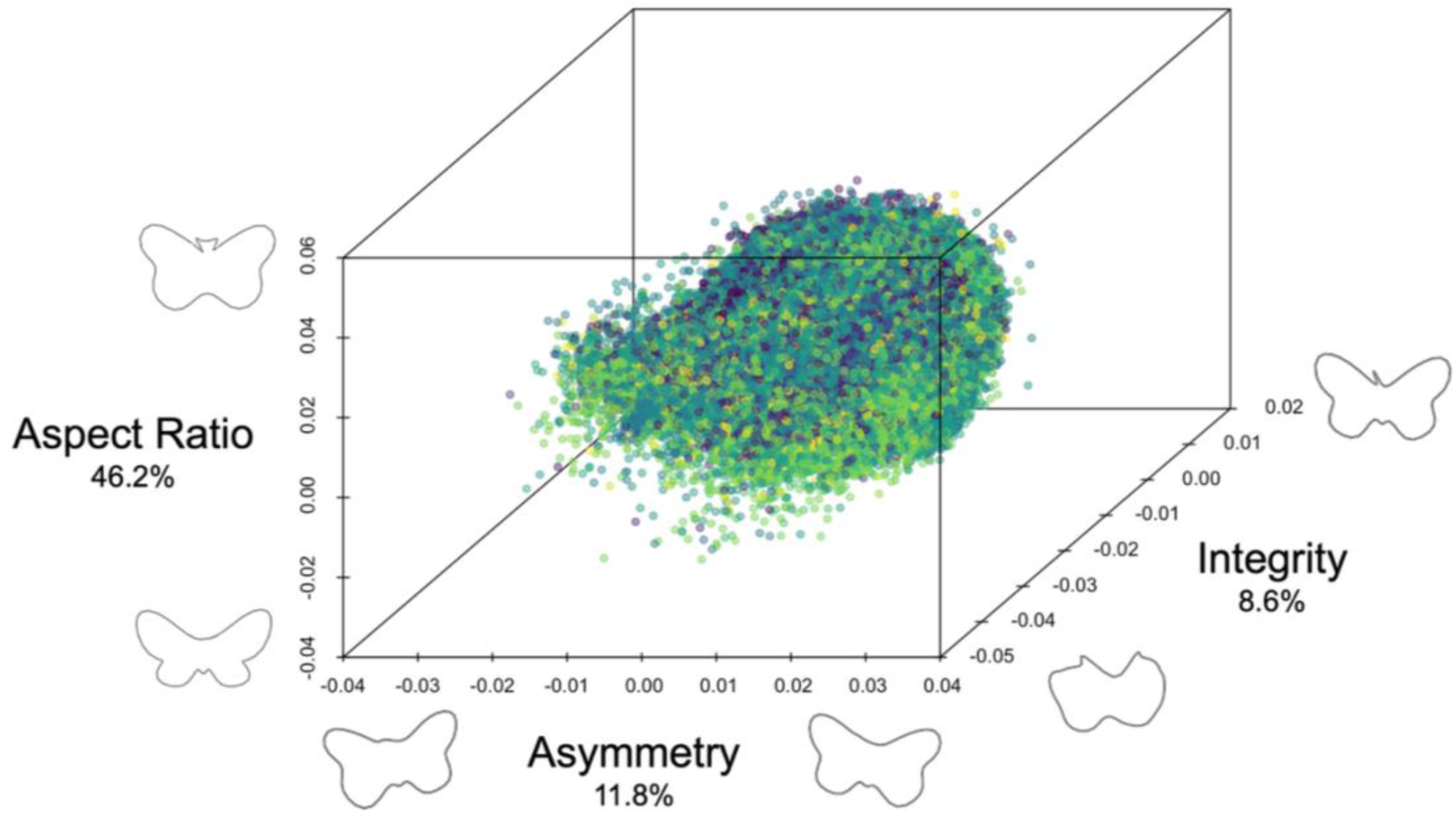
A three-dimensional morphospace, showing the variation in butterfly shape across the 178,359 specimens, indicating the major axes of wing shape variation are aspect ratio (corresponding to PC1 value), asymmetry (corresponding to absolute PC2 value), and integrity (corresponding to PC3 value). Each specimen is represented by a single point, and point colours represent different species identity. Overlaid are representations of the shape of specimens corresponding to low and high values of each PC axis, assuming the values of the other PCs are zero. PC1 explained an axis of variation of long, narrow wings to short, broad wings. Since specimens with low PC1 values represent specimens with a high aspect ratio (long, narrow wings), the values for this axis appear reversed. PC2 represents an axis of variation from right-biased to left-biased wing asymmetry. PC3 appeared to correspond to wing integrity, where low integrity wings corresponded to low PC3 values; however, very few specimens had low wing integrity.

Visual inspection of a subset of the associated images, extracted binary images from mothra, and outlines in Momocs suggest that low PC3 values were indicative of poorer outline quality in some ventrally photographed specimens (specifically relating to the body of the specimen). This was particularly apparent in the species *Hipparchia semele*, *Plebejus argus*, *Polygonia c-album*, and *Polyommatus icarus*, but importantly there did not appear to be systematic issues in outlines extracted from ventral specimens. Since the wing outlines of specimens with low PC3 values were generally of sufficient quality, these specimens were not systematically excluded from the morphospace.

We found high intraspecies and intrafamilial variation in wing aspect ratio and wing asymmetry (Fig. 5). The variance composition analysis revealed that 27.68% of variation in wing aspect ratio was explained by species and 0.58% by family (with the remaining 71.74% being residual variation). Contrastingly, 9.03% of variation in wing asymmetry was explained by species and 3.51% by family (with 87.46% residual variation).

**Fig. 5.**
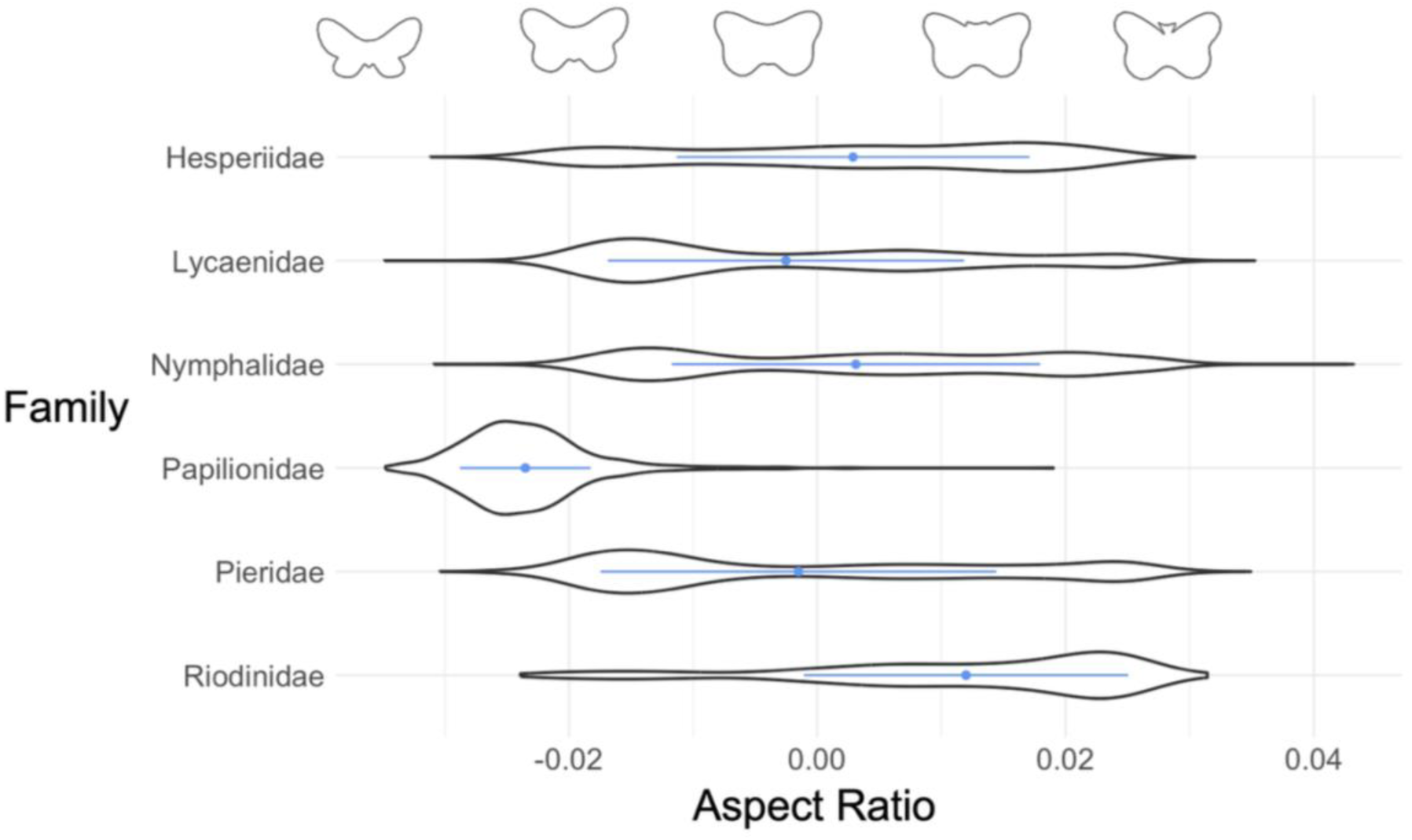
Five of the six families studied showed high variation in wing aspect ratio (corresponding to PC1 value), but the Papilionidae, represented in the dataset by a single species, showed limited variation in wing aspect ratio. The mean aspect ratio for each family is shown as a blue point, with lines representing the standard deviation. Outlines representative of specific aspect ratio values (assuming PC2 and PC3 values of zero) are shown above the plot. Since specimens with low PC1 values represent specimens with a high aspect ratio (long, narrow wings), the values for this axis appear flipped.

Symmetry as calculated using the ‘symmetry’ function in Momocs was weakly positively correlated to PC1 value (r=0.053, p<0.001), moderately positively correlated to PC2 and (r=0.365, p<0.001), and moderately negatively correlated to PC3 value (r=-0.274, p<0.001). However, wing symmetry was strongly positively correlated to the *absolute* PC2 value (r=0.688, p<0.001), which was also associated with wing asymmetry according to the morphospace.

### Trends in wing aspect ratio (PC1)

We found evidence that wing aspect ratio has changed over the 20^th^ century for 68% of the 34 species considered in the linear model (Fig. 6). Eighteen species showed increases in wing aspect ratio (became longer and narrower) and five decreases (became shorter and broader). In particular, *Aglais urticae* showed strong increases in wing aspect ratio over time (ES_linear model_ = -0.149, SE=0.0186). The increase in aspect ratio over the last century was particularly apparent when looking at only the upper quantiles (τ=0.9), where only one species (*Pieris napi*) decreased in wing aspect ratio over time.

**Fig. 6:**
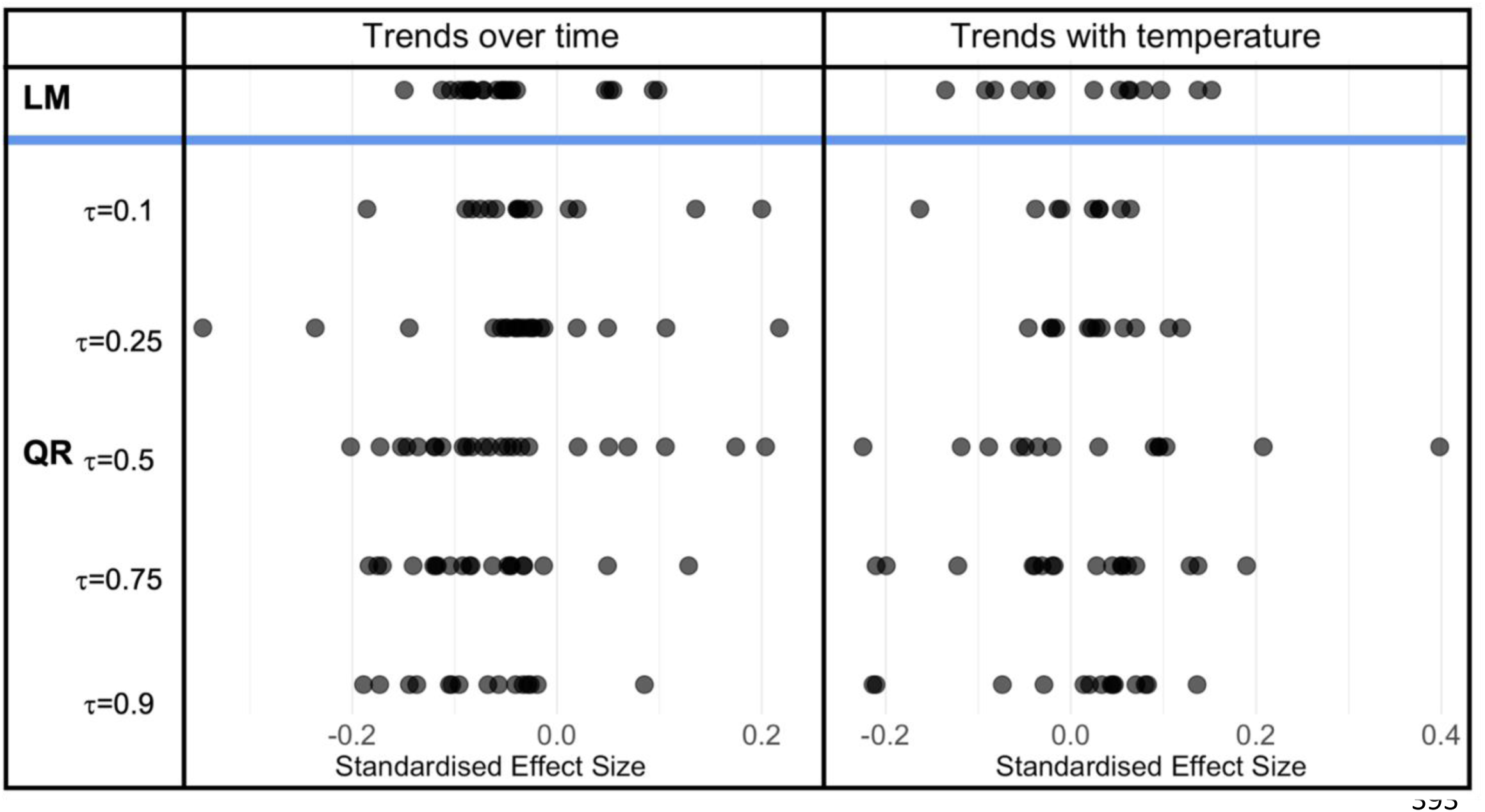
Butterfly wing aspect ratio (PC1 value) changed significantly over time and with temperature in many species, though the strength and direction of trends differed between species. Each filled circle represents the standard effect size for each species showing a significant trend in the linear model (LM) and quantile regression model (QR) for the model with wing aspect ratio as a response variable. For the quantile regression model, different quantiles of the data (indicated by τ) were considered to compare the responses between specimens of extreme versus moderate wing aspect ratios. Decreasing PC1 indicates that wings became longer and narrower, representing an increase in wing aspect ratio.

In eight species wings decreased in wing aspect ratio (becoming shorter and broader) with temperature, while in six species wings increased in wing aspect ratio (Fig. 6). *Polygonia c-album* showed a particularly strong trend of decreasing wing aspect ratio with temperature (ES_linear model_=0.152, SE=0.043 ; ES_τ=0.5_= 0.398, SE=0.141), while *Melitaea athalia* showed strong increases in aspect ratio with temperature (ES_linear model_= -0.135, SE=0.040). In some species, trend directions differed between quantiles, indicating that the variance of aspect ratio varied across temperatures. For example, in *Plebejus argus*, aspect ratio increased with temperature at low quantiles (ES_τ=0.1_= -0.038, SE=0.009) and decreased at high quantiles (ES_τ=0.9_=0.136, SE=0.025), suggesting that variance in aspect ratio increased with temperature (Fig. 3; Left), primarily by individuals exhibiting more extreme longer and broader phenotypes at hotter temperatures.

### Trends in wing asymmetry (absolute PC2 value)

Generally wing asymmetry did not change over time, since most species (22; 65%) did not show significant trends (Fig. 7). For species that did, however, wings tended to become more asymmetrical over time (11 species in the linear model), with only one species (*Melanargia galathea*) becoming less asymmetrical over time (ES_linear model_=-0.093, SE=0.018). Trends in *Melanargia galathea* were stronger at higher quantiles, indicating decreasing variance in asymmetry over time (ES_τ=0.1_=-0.013, SE=0.005 ; ES_τ=0.25_= -0.032, SE=0.011; ES_τ=0.5_=-0.087, SE=0.018 ; ES_τ=0.75_= -0.159, SE=0.029; ES_τ=0.9_=-0.168, SE=0.067). Species’ trends of asymmetry over time sometimes differed in direction between quantiles, indicating changes in the variance of asymmetry over time, for example in *Argynnis paphia* (ES_τ=0.1_= -0.015, SE=0.007; ES_τ=0.9_= 0.159, SE=0.077; corresponding to increasing variance over time).

**Fig. 7:**
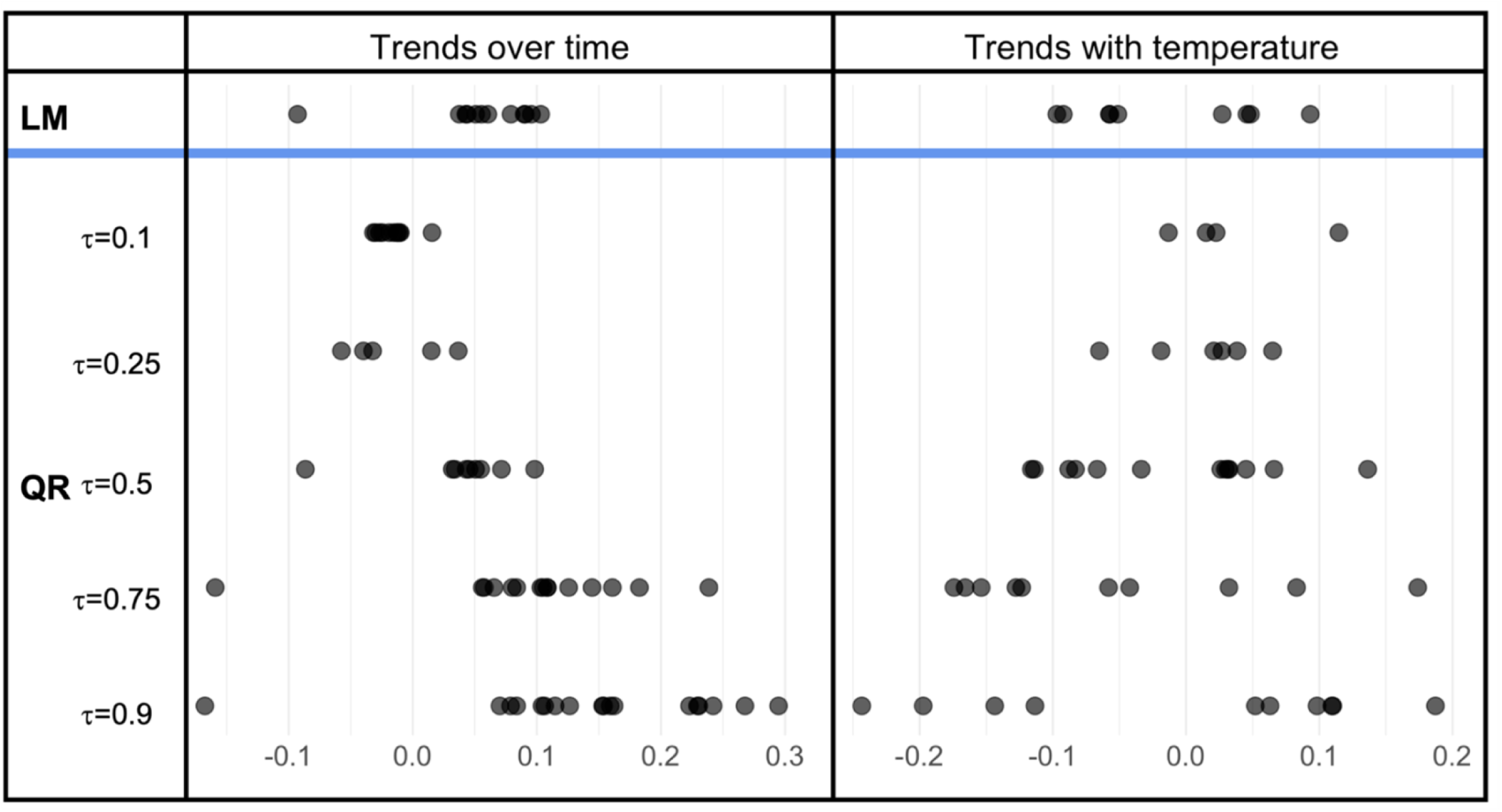
Butterfly wing asymmetry (absolute PC2 value) changed significantly over time and with temperature in some species, though the strength and direction of trends differed between species. Each filled circle represents the standard effect size for each species showing a significant trend in the linear model (LM) and quantile regression model (QR) for the model with wing asymmetry as a response variable. For the quantile regression model, different quantiles of the data (indicated by τ) were considered to compare the responses between specimens of extreme versus moderate wing asymmetry.

Generally wing asymmetry did not change with temperature, since most species (25; 74%) did not show significant trends (Fig. 7). Of species showing significant trends, four species showed increases in, and five species decreases in, asymmetry with temperature. These trends were consistently stronger at higher quantiles, indicating that variance in asymmetry is larger at higher temperatures (e.g. *Hipparchia semele*; ES_linear model_=0.093, SE=0.021 ; ES_τ=0.1_=n.s. ; ES_τ=0.25_= 0.021, SE=0.006; ES_τ=0.5_=0.045, SE=0.018; ES_τ=0.75_=0.174, SE=0.020; ES_τ=0.9_=0.187, SE=0.076).

## Discussion

In developing a high throughput method for extracting the dimensions of butterfly shape, we identified the major axes of variation in wing shape in UK butterflies as aspect ratio (PC1) and asymmetry (PC2). We find evidence of species-specific increases in wing aspect ratio over time, but trends of wing aspect ratio with temperature were mixed.

### Trends in wing aspect ratio

The major axis of shape variation corresponded to wing aspect ratio, which in butterflies is linked to flight strategy (review: Le Roy, Debat & Llaurens, 2019). Consistent with our results, a study on tropical butterflies found that the major axis of variation in shape in rainforest understorey butterflies was also aspect ratio (Cespedes, Penz & DeVries, 2015). We found high intraspecific variation in aspect ratio (27.68% of variation was explained by species), though the level of intraspecific variation differed between species. This could result from ecomorphological partitioning of butterflies with different flight strategies, since the wing traits favouring flapping and gliding flight are generally negatively correlated with one another (Cespedes, Penz & DeVries, 2015; Chai & Srygley, 1990). Alternatively, variation could result from phenotypic plasticity or mate selection preferences; wing shape is known to vary in *Heliconius* butterflies differing in host plant (Jorge et al., 2011). We note that we do not control for phylogeny, taxonomic bias, or sampling in these data; future work should explore whether interspecific variation in wing aspect ratio is related to differences in habitat, or due to phylogenetic relatedness. The Papilionidae tended to have the lowest PC1 values (long, narrow wings). This family was represented by a single species in the dataset (*Papilio machaon*, the swallowtail butterfly); the low aspect ratio values may be capturing the tail-like extensions of the hindwings (Koutrouditsou & Nudds, 2021) in this case.

#### Aspect ratio generally increased over time

We tended to see increases in aspect ratio over time (wings becoming longer and narrower) perhaps reflecting a greater requirement for gliding flight (Cespedes, Penz & DeVries, 2015; Le Roy, Debat & Llaurens, 2019). Since gliding flight minimises energy expenditure while maximising flight time or distance (Cespedes, Penz & DeVries, 2015; Le Roy, Debat & Llaurens, 2019), this may reflect selection to reduce the energetic costs of flight, which could be beneficial in modern landscapes where nutrient resources (flowers) have become reduced and fragmented. Alternatively, low aspect-ratio wings tend to allow higher agility and manoeuvrability, which tend to be favoured in more spatially cluttered habitats (Cespedes, Penz & DeVries, 2015; DeVries, Penz & Hill, 2010; Le Roy, Debat & Llaurens, 2019). Therefore, these results may be due to habitat change creating more open habitats, selecting for energy-efficient gliding flight as opposed to energy-intensive flapping flight. Other land-use factors may also underpin our results; evidence from the butterfly *Pararge aegeria* finds that individuals at the northern range boundary tend to have lower wing aspect ratios (Hill, Thomas & Blakeley, 1999).

#### Species-specific changes with temperature

Trends in aspect ratio with temperature were more mixed, suggesting that trends are species-specific, reflecting the different ecologies, life-histories, and histories of species and communities in their habitat. Some species (e.g. *Polygonia c-album*) showed particularly strong trends of decreasing wing aspect ratio with temperature, while in other species (e.g. *Melitaea athalia*) wing aspect ratio increased with increasing temperature. Variance in aspect ratio change with temperature in some species. In *Plebejus argus*, which has experience long-term increases in abundance (UKBMS, 2024), this was primarily due to individuals exhibiting more extreme longer and broader phenotypes at hotter temperatures. Evidence shows that butterflies with a higher wing aspect ratio show a greater thermal buffering ability than butterflies with lower wing aspect ratios (Laird-Hopkins et al., 2023). In species where wings decreased in aspect ratio, this may reflect selection for increased heat loss through basking, which is favoured by wings with a larger surface area (Bladon et al., 2020). Therefore, these mixed results may reflect different thermal buffering strategies in UK butterflies. Increased phenotypic differentiation in wing aspect ratio is associated with more northern latitudes in the butterfly *Pararge aegeria* (Vandewoestijne & Van Dyck, 2010). Thus, even where species’ mean aspect ratio did not change, changes to phenotypic variability may still impact populations.

#### Trends in wing asymmetry

The second axis of shape variation corresponded to wing asymmetry. Species showed high intraspecific variation in wing asymmetry (9.03% of variation was explained by species and 3.51% was explained by family) suggesting that this axis may be capturing fluctuating asymmetry as opposed to directional asymmetry. Since fluctuating asymmetry is thought to be a proxy of developmental stress (Badyaev, Foresman & Fernandes, 2000; Gerard et al., 2018; Palmer & Strobeck, 1986), our results indicate that developmental stress is not consistent across UK butterflies. Differences in wing angle or positioning due to the way specimens were pinned could have contributed to these results. However, since butterflies are large-winged insects that are typically pinned with the wings flat (Altizer & Davis, 2009; Ortega Ancel et al., 2017), and we had no *a priori* reason to assume that there should be systematic differences in pinning position between species, we concluded that these trends likely had a biological basis. We did not find evidence of directional asymmetry, contrasting evidence from the UK butterfly *Pararge aegeria*, where males show directional asymmetry of size and shape, which is thought to be adaptive (Windig & Nylin, 1999). Future work should explicitly consider whether there is a relationship between extinction risk and wing shape asymmetry.

#### Population-specific increases over time

Wing asymmetry did not change over time in the majority of species, but of the species that showed significant trends, wings tended to become more asymmetrical over time. This may be associated with factors like population declines (Adamski & Witkowski, 2003), food stress (Pellegroms et al., 2009), or even habitat fragmentation (though previously linked to size asymmetry; Pignataro et al., 2023). Interestingly, *Melanargia galathea* became less asymmetrical and the variance of asymmetry decreased over time. This may be associated with decreasing developmental stress, since *Melanargia galathea* has showed strong population increases since 1976 (UKBMS, 2024). Variance in asymmetry both decreased and increased over time across species. Where variance increased, this may indicate population-specific responses to stress, and where variance decreased this may indicate decreasing developmental stress or environmental filtering.

#### Population-specific changes with temperature

Wing asymmetry did not generally change with temperature, and species with significant trends varied in trend direction. Nonetheless, we find evidence of larger variance in asymmetry at higher temperatures, which may be indicative of increased developmental stress due to climate warming. Evidence from the literature is mixed, with some studies on insect pollinators finding evidence of increasing fluctuating shape asymmetry with increasing temperatures (Arce et al., 2023; Adamski & Witkowski, 2003), and others finding no relationship between temperature and shape (Gerard et al., 2018) or size asymmetry (Symanski & Redak, 2021). These results suggest that the impact of increasing environmental temperatures on wing shape asymmetry is complex, species-specific, and population-specific. Future work must consider these nuances to understand the impacts of environmental change on populations.

### Future work and Limitations

Our work reinforces the value of machine learning methods in studying complex functional traits like wing shape across taxonomic groups. The third axis of shape variation in our data corresponded partially to wing integrity, but was not fully attributable to a specific component of shape, and thus was excluded from data analysis. This suggests that our methodology is conservative since we only analysed axes with attributable biological signal. Nonetheless, though specimens with low wing integrity did not appear to have systematic issues with wing outline extraction, advances in image segmentation would mitigate any issues.

Mothra was unable to identify the sex of specimens in this study, and due to the size of the dataset manual classification of sex was unfeasible. However, male and female butterflies are known to have different ecologies and experience different selection pressures, which can result in sexual dimorphism of butterfly wing shape (Allen, Zwaan & Brakefield, 2011). Differences in flight behaviour between sexes (Cespedes, Penz & DeVries, 2015) should favour selection for different morphological characteristics in male and female butterflies (Allen, Zwaan & Brakefield, 2011; Cespedes, Penz & DeVries, 2015; DeVries, Penz & Hill, 2010), and indeed evidence supports this (Berwaerts, Van Dyck & Aerts, 2002; Cespedes, Penz & DeVries, 2015). Future work must therefore consider sex, which is likely to explain some of the intraspecific variation in wing aspect ratio in our data, if we are to gain a more nuanced understanding of the factors governing long-term changes in butterfly wing shape.

## Conclusions

We demonstrate the suitability of novel machine learning techniques and Elliptical Fourier Analysis for extracting morphological data from museum collections. These same techniques could be applied to other digitised museum collections to study long-term trends in shape across taxonomic groups. We also find that, for a subset of the species for which we have available data, wing aspect ratio and asymmetry may be increasing over time. These changes may affect flight strategies and behaviour, and potentially even fitness, with consequent effects for butterfly populations, other species, and the delivery of pollination services. Future studies should make use of these under-utilised data sources and novel computational techniques to better understand the impacts of environmental change on populations, species, and ecosystems.

