## Supplementary Materials for "Revealing centurial changes in butterfly wing shape via the automated analysis of museum specimens"

### 1.1: Data

***Supp. Table 1: The number of specimens of each species included in the morphospace.*** *65 species were included in the morphospace.*

| Species | Number of specimens |
|---|---|
| *Aglais urticae* | 3728 |
| *Anthocharis cardamines* | 2940 |
| *Apatura iris* | 569 |
| *Aphantopus hyperantus* | 3735 |
| *Aporia crataegi* | 464 |
| *Argynnis adippe* | 1538 |
| *Argynnis aglaja* | 1899 |
| *Argynnis paphia* | 2217 |
| *Aricia agestis* | 4192 |
| *Aricia artaxerxes* | 1846 |
| *Callophrys rubi* | 1441 |
| *Carterocephalus palaemon* | 950 |
| *Celastrina argiolus* | 2424 |
| *Coenonympha pamphilus* | 4055 |
| *Coenonympha tullia* | 5341 |
| *Colias alfacariensis* | 67 |
| *Colias croceus* | 2818 |
| *Colias hyale* | 678 |
| *Cupido minimus* | 1951 |
| *Cyaniris semiargus* | 236 |
| *Erebia aethiops* | 1757 |
| *Erebia epiphron* | 1470 |
| *Erynnis tages* | 1233 |
| *Euphydryas aurinia* | 6459 |
| *Gonepteryx rhamni* | 892 |
| *Hamearis lucina* | 1725 |
| *Hesperia comma* | 883 |
| *Hipparchia semele* | 2925 |
| *Inachis io* | 1326 |
| *Issoria lathonia* | 126 |
| *Lasiommata megera* | 1928 |
| *Leptidea sinapis* | 1780 |
| *Limenitis camilla* | 1198 |
| *Lycaena dispar* | 1110 |
| *Lycaena phlaeas* | 6076 |
| *Lysandra bellargus* | 8710 |
| *Lysandra coridon* | 21923 |
| *Maculinea arion* | 1842 |
| *Maniola jurtina* | 7289 |
| *Melanargia galathea* | 2758 |
| *Melitaea athalia* | 2272 |
| *Melitaea cinxia* | 1878 |
| *Neozephyrus quercus* | 1212 |
| *Nymphalis antiopa* | 136 |
| *Nymphalis polychloros* | 823 |

| | |
|---|---|
| *Ochlodes venata* | 1289 |
| *Papilio machaon* | 833 |
| *Pararge aegeria* | 3026 |
| *Pieris brassicae* | 2455 |
| *Pieris napi* | 8256 |
| *Pieris rapae* | 2465 |
| *Plebejus argus* | 9316 |
| *Polygonia c-album* | 2088 |
| *Polyommatus icarus* | 12573 |
| *Pontia daplidice* | 162 |
| *Pyrgus malvae* | 1701 |
| *Pyronia tithonus* | 3042 |
| *Satyrium pruni* | 910 |
| *Satyrium w-album* | 1012 |
| *Thecla betulae* | 1220 |
| *Thymelicus acteon* | 732 |
| *Thymelicus lineola* | 879 |
| *Thymelicus sylvestris* | 1270 |
| *Vanessa atalanta* | 1308 |
| *Vanessa cardui* | 1002 |

***Supp. Table 2: The number of species of each family included in the morphospace.*** *65 species from six families were included in the morphospace.*

| Family | Number of species |
|---|---|
| Nymphalidae | 27 |
| Pieridae | 11 |
| Lycaenidae | 17 |
| Hesperiidae | 8 |
| Riodinidae | 1 |
| Papilionidae | 1 |

***Supp. Table 3: The number of specimens of each species included in the models.*** *Species with fewer than 1000 observations were excluded from this dataset. 34 species were included in the models.*

| **Species** | **Number of specimens** |
|---|---|
| *Aglais urticae* | 1624 |
| *Anthocharis cardamines* | 1853 |
| *Aphantopus hyperantus* | 2589 |
| *Argynnis adippe* | 1023 |
| *Argynnis aglaja* | 1393 |
| *Argynnis paphia* | 1497 |
| *Aricia agestis* | 2176 |
| *Aricia artaxerxes* | 1345 |
| *Callophrys rubi* | 1043 |
| *Celastrina argiolus* | 1396 |
| *Coenonympha pamphilus* | 3356 |
| *Coenonympha tullia* | 4465 |
| *Colias croceus* | 1285 |
| *Cupido minimus* | 1459 |
| *Erebia aethiops* | 1283 |
| *Euphydryas aurinia* | 3003 |
| *Hipparchia semele* | 2323 |
| *Lasiommata megera* | 1334 |
| *Leptidea sinapis* | 1280 |
| *Lycaena phlaeas* | 4370 |
| *Lysandra bellargus* | 7533 |
| *Lysandra coridon* | 20031 |
| *Maculinea arion* | 1311 |
| *Maniola jurtina* | 5849 |
| *Melanargia galathea* | 2152 |
| *Melitaea athalia* | 1641 |
| *Pararge aegeria* | 1693 |
| *Pieris napi* | 3418 |
| *Pieris rapae* | 1340 |
| *Plebejus argus* | 8041 |
| *Polygonia c-album* | 1100 |
| *Polyommatus icarus* | 10598 |
| *Pyrgus malvae* | 1264 |
| *Pyronia tithonus* | 2443 |

### 1.2: Source of NHM metadata

***Supp. Table 4: A table listing the species studied and the DOI representing the source of the metadata for this species.***

| Species | DOI |
|---|---|
| *Aglais urticae* | https://doi.org/10.5519/qd.blnju1yz |
| *Anthocharis cardamines* | https://doi.org/10.5519/qd.rnrz8nlz |
| *Apatura iris* | https://doi.org/10.5519/qd.w7l9hi0y |
| *Aphantopus hyperantus* | https://doi.org/10.5519/qd.8p2yipk2 |
| *Aporia crataegi* | https://doi.org/10.5519/qd.083ag8mz |
| *Argynnis adippe* | https://doi.org/10.5519/qd.3axc3psf |
| *Argynnis aglaja* | https://doi.org/10.5519/qd.nnjibwbk |
| *Argynnis paphia* | https://doi.org/10.5519/qd.ydzkkawb |
| *Aricia agestis* | https://doi.org/10.5519/qd.3mozbc4p |
| *Aricia artaxerxes* | https://doi.org/10.5519/qd.222j8qem |
| *Callophrys rubi* | https://doi.org/10.5519/qd.1hztjas9 |
| *Carterocephalus palaemon* | https://doi.org/10.5519/qd.twyx9xv5 |
| *Celastrina argiolus* | https://doi.org/10.5519/qd.vpbo6ten |
| *Coenonympha pamphilus* | https://doi.org/10.5519/qd.3jk9oxe6 |
| *Coenonympha tullia* | https://doi.org/10.5519/qd.qo4ozy0j |
| *Colias alfacariensis* | https://doi.org/10.5519/qd.s03vp2z5 |
| *Colias croceus* | https://doi.org/10.5519/qd.dur7elje |
| *Colias hyale* | https://doi.org/10.5519/qd.1y58e6hm |
| *Cupido minimus* | https://doi.org/10.5519/qd.tfyvhk57 |
| *Cyaniris semiargus* | https://doi.org/10.5519/qd.mex24ppa |
| *Erebia aethiops* | https://doi.org/10.5519/qd.az6eqt3a |
| *Erebia epiphron* | https://doi.org/10.5519/qd.5f5mbg65 |
| *Erynnis tages* | https://doi.org/10.5519/qd.g0lo6jx4 |
| *Euphydryas aurinia* | https://doi.org/10.5519/qd.8ygpagik |
| *Gonepteryx rhamni* | https://doi.org/10.5519/qd.a6jsjrre |
| *Hamearis lucina* | https://doi.org/10.5519/qd.32dh1l33 |
| *Hesperia comma* | https://doi.org/10.5519/qd.em35wpfg |
| *Hipparchia semele* | https://doi.org/10.5519/qd.9jq1gd70 |
| *Aglais io* | https://doi.org/10.5519/qd.yftba23p |
| *Inachis io* | https://doi.org/10.5519/qd.d39rhnj4 |
| *Issoria lathonia* | https://doi.org/10.5519/qd.43g4qo41 |
| *Lasiommata megera* | https://doi.org/10.5519/qd.d1709xuc |
| *Leptidea sinapis* | https://doi.org/10.5519/qd.oksm7y5p |
| *Limenitis camilla* | https://doi.org/10.5519/qd.kzxu9yi5 |
| *Lycaena dispar* | https://doi.org/10.5519/qd.4xow91yc |
| *Lycaena phlaeas* | https://doi.org/10.5519/qd.cjh57ggx |
| *Lysandra bellargus* | https://doi.org/10.5519/qd.wcdtn2e5 |
| *Lysandra coridon* | https://doi.org/10.5519/qd.dozkvt91 |
| *Maculinea arion* | https://doi.org/10.5519/qd.2fpuao2l |
| *Maniola jurtina* | https://doi.org/10.5519/qd.ntpcptzg |
| *Melanargia galathea* | https://doi.org/10.5519/qd.c8th0eco |
| *Melitaea athalia* | https://doi.org/10.5519/qd.imcxscx5 |
| *Melitaea cinxia* | https://doi.org/10.5519/qd.izem6ic3 |
| *Neozephyrus quercus* | https://doi.org/10.5519/qd.ylesdx0v |
| *Nymphalis antiopa* | https://doi.org/10.5519/qd.s1lmlh52 |
| *Nymphalis polychloros* | https://doi.org/10.5519/qd.m3owr97s |
| *Ochlodes venata* | https://doi.org/10.5519/qd.b0ifrkuk |

| | |
|---|---|
| *Papilio machaon* | https://doi.org/10.5519/qd.ok9n6s44 |
| *Pararge aegeria* | https://doi.org/10.5519/qd.2m9hr2g3 |
| *Pieris brassicae* | https://doi.org/10.5519/qd.icc28g7s |
| *Pieris napi* | https://doi.org/10.5519/qd.5cw3ce6n |
| *Pieris rapae* | https://doi.org/10.5519/qd.julivhnl |
| *Plebejus argus* | https://doi.org/10.5519/qd.dw210igj |
| *Polygonia c-album* | https://doi.org/10.5519/qd.rpnuldr0 |
| *Polyommatus icarus* | https://doi.org/10.5519/qd.7efutdu5 |
| *Pontia daplidice* | https://doi.org/10.5519/qd.a4gk0aes |
| *Pyrgus malvae* | https://doi.org/10.5519/qd.d05b4wyd |
| *Pyronia tithonus* | https://doi.org/10.5519/qd.oqrsvdzb |
| *Satyrium pruni* | https://doi.org/10.5519/qd.cr54srx8 |
| *Satyrium w-album* | https://doi.org/10.5519/qd.n6wrrh7j |
| *Thecla betulae* | https://doi.org/10.5519/qd.azwir6eb |
| *Thymelicus acteon* | https://doi.org/10.5519/qd.qmrgwnrk |
| *Thymelicus lineola* | https://doi.org/10.5519/qd.e9qdewm7 |
| *Thymelicus sylvestris* | https://doi.org/10.5519/qd.09y6141h |
| *Vanessa atalanta* | https://doi.org/10.5519/qd.cfqcoikm |
| *Vanessa cardui* | https://doi.org/10.5519/qd.loab32q4 |

### 1.3: Results

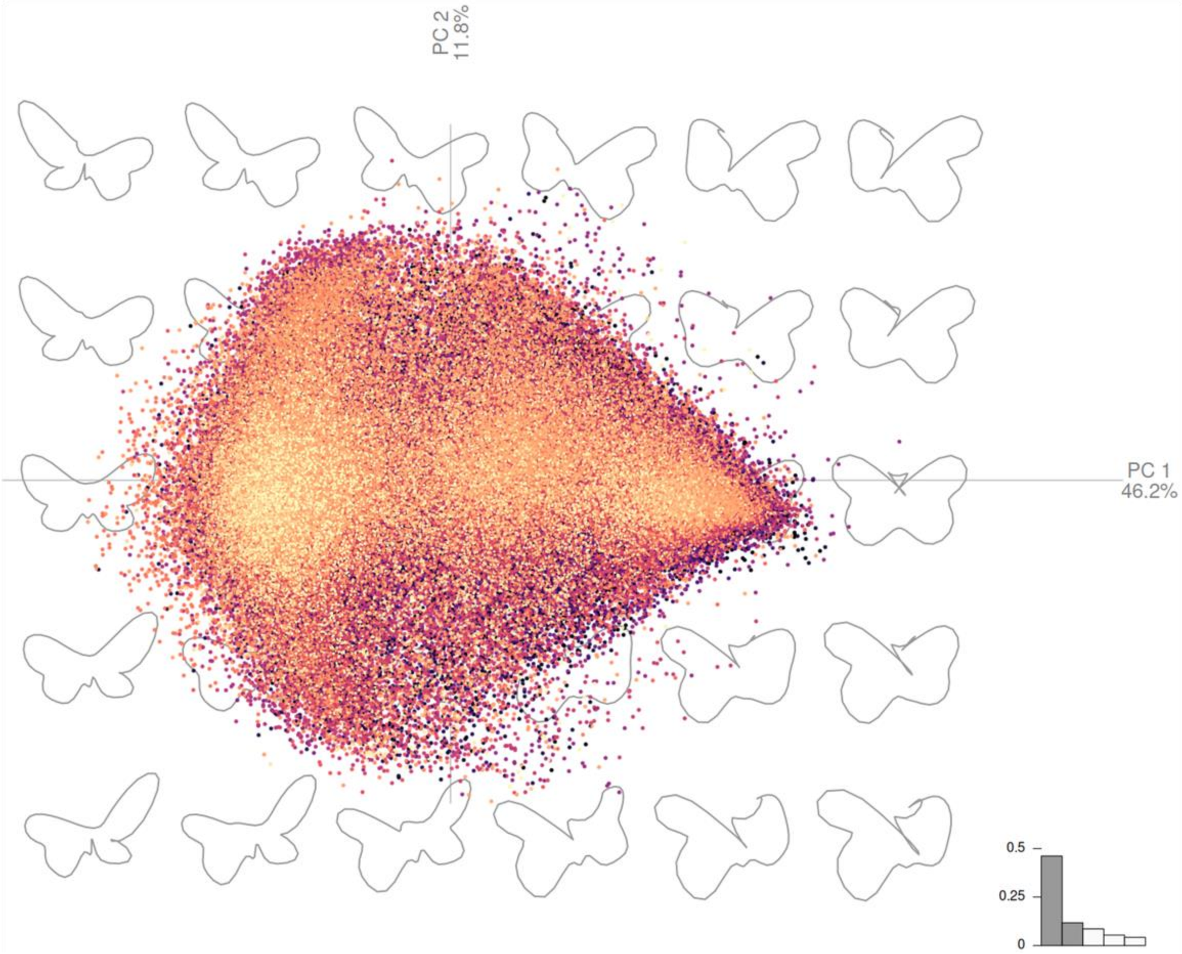


***Supp. Fig. 1: A 2-dimensional morphospace showing the variation in shape across the dataset for PC1 (x axis) and PC2 (y axis).*** *Each point represents a specimen, and points are coloured according to species. Outlines representing shape variation are shown. A scree plot is shown at the bottom right of the PCA.*

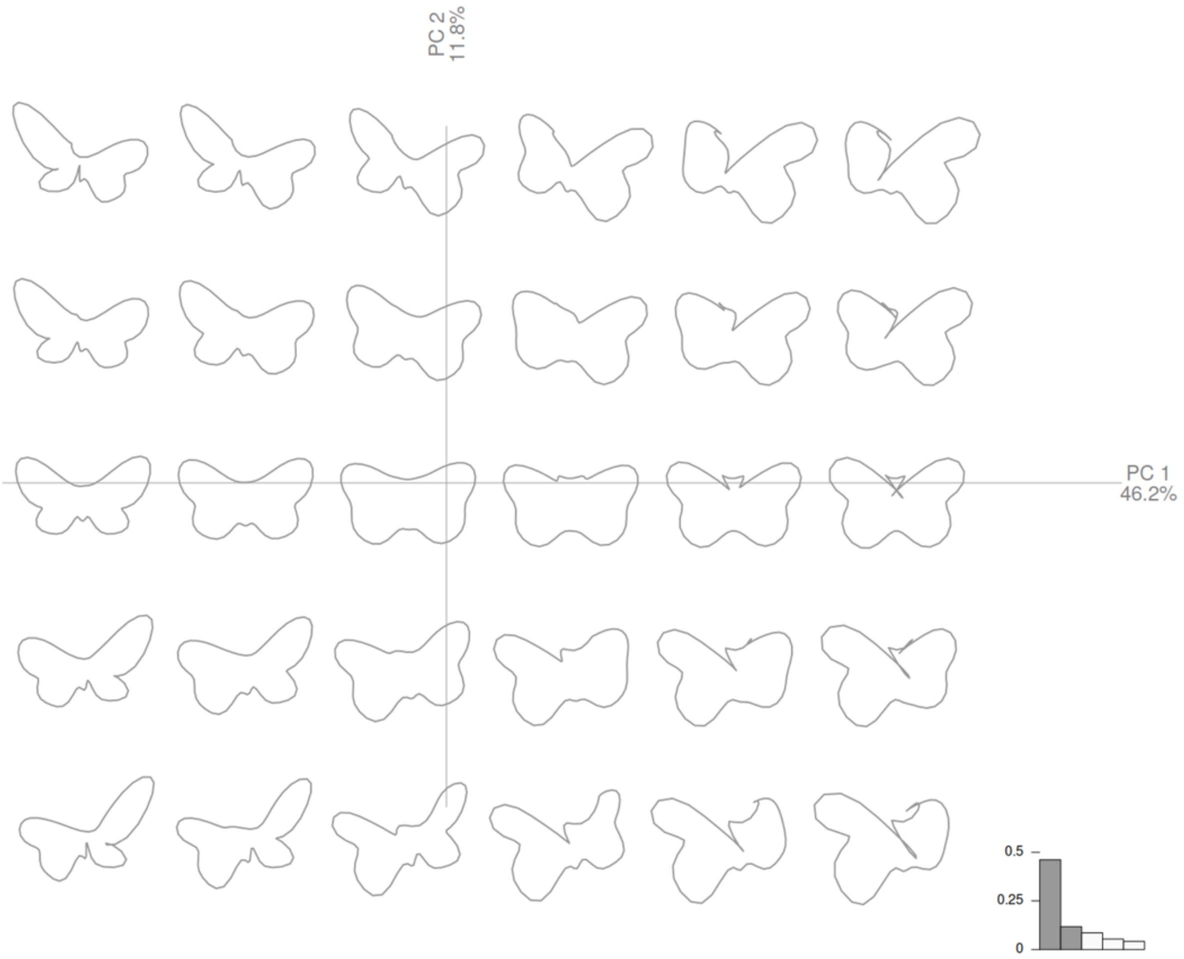


***Supp. Fig. 2: A 2-dimensional morphospace showing the variation in shape across the dataset for PC1 (x axis) and PC2 (y axis).*** *Outlines representing shape variation are shown. A scree plot is shown at the bottom right of the PCA. Despite some abnormal outlines in the morphospace, specimens did not fit these abnormal outlines.*

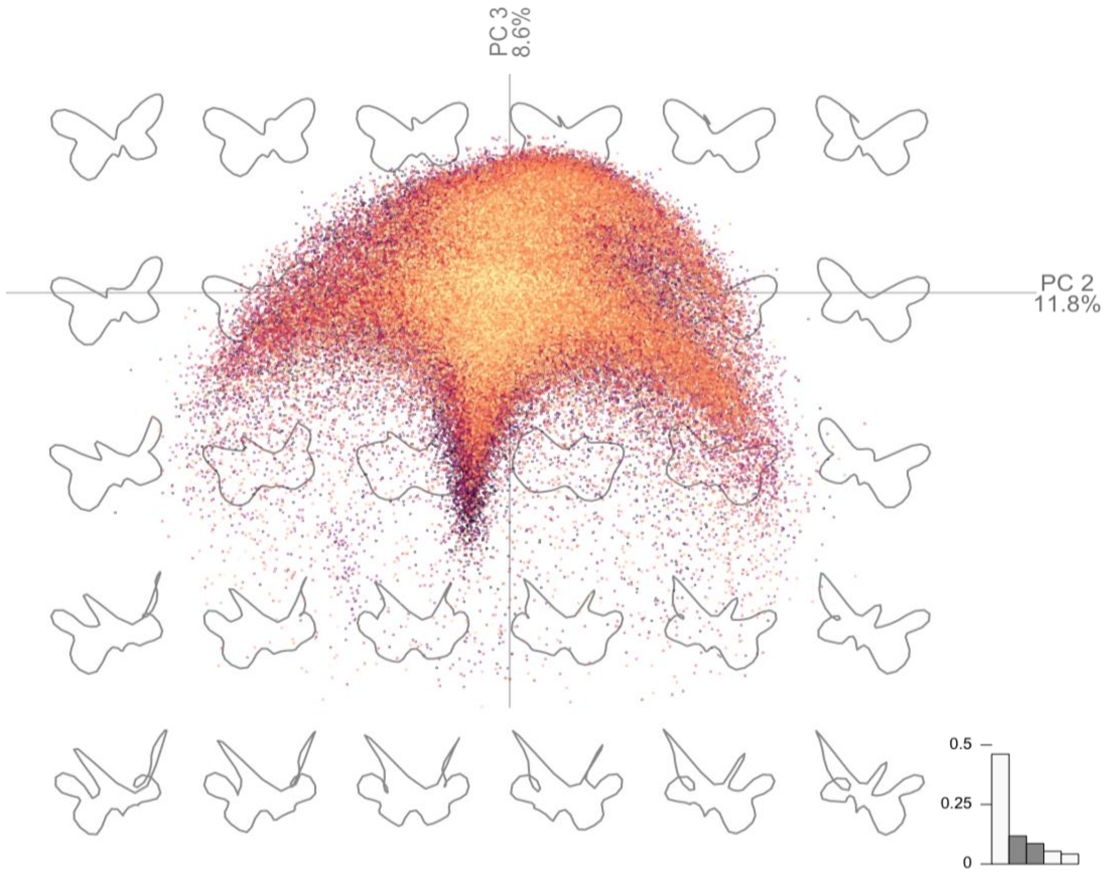


***Supp. Fig. 3: A 2-dimensional morphospace showing the variation in shape across the dataset for PC2 (x axis) and PC3 (y axis).*** *Each point represents a specimen, and points are coloured according to species. Outlines representing shape variation are shown. A scree plot is shown at the bottom right of the PCA.*

***Supp. Table 5: The PC1 linear model results.***

| Term | Estimate | Std. Error | t value | P value |
|---|---|---|---|---|
| *Aglais urticae* | 0.82594 | 0.02226 | 37.09833 | 0 |
| *Anthocharis cardamines* | 0.54163 | 0.02061 | 26.28205 | 0 |
| *Aphantopus hyperantus* | 0.50041 | 0.01778 | 28.13944 | 0 |
| *Argynnis adippe* | -0.45731 | 0.02871 | -15.92763 | 0 |
| *Argynnis aglaja* | 0.56606 | 0.02414 | 23.44565 | 0 |
| *Argynnis paphia* | 0.02777 | 0.02589 | 1.07251 | 0.28349 |
| *Aricia agestis* | 0.20139 | 0.01963 | 10.25749 | 0 |
| *Aricia artaxerxes* | 0.32181 | 0.0474 | 6.78863 | 0 |
| *Callophrys rubi* | -0.72268 | 0.02763 | -26.15316 | 0 |
| *Celastrina argiolus* | -0.84861 | 0.0255 | -33.27267 | 0 |
| *Coenonympha pamphilus* | 0.40082 | 0.0156 | 25.69048 | 0 |
| *Coenonympha tullia* | 0.54873 | 0.02007 | 27.34487 | 0 |
| *Colias croceus* | 1.28549 | 0.04249 | 30.25562 | 0 |
| *Cupido minimus* | -0.25339 | 0.02363 | -10.72275 | 0 |
| *Erebia aethiops* | 0.96073 | 0.04929 | 19.49214 | 0 |
| *Euphydryas aurinia* | -0.41195 | 0.0165 | -24.97156 | 0 |
| *Hipparchia semele* | 0.05936 | 0.01864 | 3.18421 | 0.00145 |
| *Lasiommata megera* | -0.39616 | 0.02568 | -15.42962 | 0 |
| *Leptidea sinapis* | -0.93821 | 0.02529 | -37.09402 | 0 |
| *Lycaena phlaeas* | 0.78726 | 0.0146 | 53.91599 | 0 |
| *Lysandra bellargus* | -0.52521 | 0.01484 | -35.40234 | 0 |
| *Lysandra coridon* | -0.12945 | 0.00639 | -20.27083 | 0 |
| *Maculinea arion* | 0.28259 | 0.056 | 5.04654 | 0 |
| *Maniola jurtina* | 0.41068 | 0.01191 | 34.48229 | 0 |
| *Melanargia galathea* | -0.59162 | 0.0229 | -25.83833 | 0 |
| *Melitaea athalia* | -0.03351 | 0.04447 | -0.75352 | 0.45114 |
| *Pararge aegeria* | 0.63738 | 0.02184 | 29.1815 | 0 |
| *Pieris napi* | -0.78372 | 0.01679 | -46.68534 | 0 |
| *Pieris rapae* | 0.16094 | 0.02443 | 6.58776 | 0 |
| *Plebejus argus* | -0.1886 | 0.00991 | -19.02761 | 0 |
| *Polygonia c-album* | -0.20618 | 0.02832 | -7.27923 | 0 |
| *Polyommatus icarus* | -0.24354 | 0.00868 | -28.04855 | 0 |
| *Pyrgus malvae* | -0.16615 | 0.02697 | -6.16031 | 0 |
| *Pyronia tithonus* | 0.50303 | 0.02007 | 25.06914 | 0 |
| temperature : *Aglais urticae* | 0.03219 | 0.02731 | 1.17858 | 0.23857 |
| temperature : *Anthocharis cardamines* | 0.07874 | 0.02663 | 2.95744 | 0.0031 |
| temperature : *Aphantopus hyperantus* | -0.05481 | 0.01602 | -3.42148 | 0.00062 |
| temperature : *Argynnis adippe* | -0.01065 | 0.04197 | -0.25365 | 0.79977 |
| temperature : *Argynnis aglaja* | 0.01486 | 0.01733 | 0.85713 | 0.39137 |
| temperature : *Argynnis paphia* | -0.01278 | 0.04336 | -0.29482 | 0.76813 |
| temperature : *Aricia agestis* | 0.13726 | 0.02792 | 4.91697 | 0 |
| temperature : *Aricia artaxerxes* | 0.0081 | 0.01926 | 0.42051 | 0.67412 |
| temperature : *Callophrys rubi* | -0.03159 | 0.02425 | -1.30264 | 0.1927 |
| temperature : *Celastrina argiolus* | 0.02405 | 0.04062 | 0.59213 | 0.55376 |
| temperature : *Coenonympha pamphilus* | 0.01468 | 0.01501 | 0.97807 | 0.32804 |
| temperature : *Coenonympha tullia* | 0.02483 | 0.01063 | 2.33527 | 0.01953 |
| temperature : *Colias croceus* | -0.0214 | 0.04403 | -0.48605 | 0.62693 |
| temperature : *Cupido minimus* | 0.09726 | 0.02768 | 3.51409 | 0.00044 |
| temperature : *Erebia aethiops* | -0.00098 | 0.01753 | -0.05577 | 0.95552 |
| temperature : *Euphydryas aurinia* | 0.01929 | 0.01838 | 1.04955 | 0.29393 |
| temperature : *Hipparchia semele* | 0.02903 | 0.01946 | 1.49206 | 0.13569 |
| temperature : *Lasiommata megera* | -0.08219 | 0.03493 | -2.35323 | 0.01861 |
| temperature : *Leptidea sinapis* | -0.09234 | 0.04281 | -2.15677 | 0.03103 |
| temperature : *Lycaena phlaeas* | 0.02772 | 0.01771 | 1.56547 | 0.11747 |
| temperature : *Lysandra bellargus* | 0.01951 | 0.01776 | 1.0985 | 0.27199 |

| | | | | |
|---|---|---|---|---|
| temperature : *Lysandra coridon* | 0.05271 | 0.01014 | 5.20087 | 0 |
| temperature : *Maculinea arion* | -0.07284 | 0.07383 | -0.98649 | 0.32389 |
| temperature : *Maniola jurtina* | 0.06184 | 0.01394 | 4.4372 | 1e-05 |
| temperature : *Melanargia galathea* | 0.03761 | 0.0336 | 1.11926 | 0.26303 |
| temperature : *Melitaea athalia* | -0.13535 | 0.03957 | -3.42053 | 0.00063 |
| temperature : *Pararge aegeria* | 0.06376 | 0.0269 | 2.37001 | 0.01779 |
| temperature : *Pieris napi* | -0.02681 | 0.01286 | -2.0857 | 0.03701 |
| temperature : *Pieris rapae* | 0.02101 | 0.02846 | 0.7382 | 0.46039 |
| temperature : *Plebejus argus* | 0.02535 | 0.01334 | 1.90018 | 0.05741 |
| temperature : *Polygonia c-album* | 0.1521 | 0.04269 | 3.56283 | 0.00037 |
| temperature : *Polyommatus icarus* | -0.03658 | 0.00837 | -4.36777 | 1e-05 |
| temperature : *Pyrgus malvae* | -0.02236 | 0.0426 | -0.52489 | 0.59966 |
| temperature : *Pyronia tithonus* | 0.02939 | 0.03105 | 0.94653 | 0.34388 |
| *Aglais urticae* : year | -0.14931 | 0.01859 | -8.03377 | 0 |
| *Anthocharis cardamines* : year | -0.02895 | 0.01729 | -1.67489 | 0.09396 |
| *Aphantopus hyperantus* : year | -0.04629 | 0.0176 | -2.63077 | 0.00852 |
| *Argynnis adippe* : year | -0.07233 | 0.0293 | -2.46887 | 0.01356 |
| *Argynnis aglaja* : year | -0.08311 | 0.0204 | -4.07462 | 5e-05 |
| *Argynnis paphia* : year | 0.05468 | 0.02187 | 2.50071 | 0.0124 |
| *Aricia agestis* : year | -0.10422 | 0.01907 | -5.46543 | 0 |
| *Aricia artaxerxes* : year | -0.07175 | 0.02394 | -2.99766 | 0.00272 |
| *Callophrys rubi* : year | -0.03415 | 0.02734 | -1.24895 | 0.21168 |
| *Celastrina argiolus* : year | -0.01888 | 0.02293 | -0.82324 | 0.41037 |
| *Coenonympha pamphilus* : year | -0.02157 | 0.01419 | -1.51952 | 0.12863 |
| *Coenonympha tullia* : year | -0.0396 | 0.01408 | -2.81334 | 0.0049 |
| *Colias croceus* : year | -0.00768 | 0.03335 | -0.23026 | 0.81789 |
| *Cupido minimus* : year | -0.08442 | 0.02403 | -3.51349 | 0.00044 |
| *Erebia aethiops* : year | 0.05141 | 0.01893 | 2.71588 | 0.00661 |
| *Euphydryas aurinia* : year | -0.0443 | 0.01508 | -2.93709 | 0.00331 |
| *Hipparchia semele* : year | -0.05384 | 0.01554 | -3.46338 | 0.00053 |
| *Lasiommata megera* : year | -0.09594 | 0.02382 | -4.02817 | 6e-05 |
| *Leptidea sinapis* : year | -0.05927 | 0.02554 | -2.32089 | 0.02029 |
| *Lycaena phlaeas* : year | -0.09043 | 0.01426 | -6.34134 | 0 |
| *Lysandra bellargus* : year | -0.00305 | 0.0122 | -0.24976 | 0.80278 |
| *Lysandra coridon* : year | -0.05057 | 0.00691 | -7.31372 | 0 |
| *Maculinea arion* : year | 0.09398 | 0.02989 | 3.14372 | 0.00167 |
| *Maniola jurtina* : year | 0.01859 | 0.01159 | 1.60432 | 0.10865 |
| *Melanargia galathea* : year | -0.01658 | 0.01663 | -0.99712 | 0.31871 |
| *Melitaea athalia* : year | 0.09857 | 0.02874 | 3.4296 | 6e-04 |
| *Pararge aegeria* : year | 0.00029 | 0.02017 | 0.01448 | 0.98845 |
| *Pieris napi* : year | 0.04724 | 0.01436 | 3.28996 | 0.001 |
| *Pieris rapae* : year | 0.01307 | 0.02307 | 0.56633 | 0.57117 |
| *Plebejus argus* : year | -0.11224 | 0.01142 | -9.83025 | 0 |
| *Polygonia c-album* : year | -0.05982 | 0.03149 | -1.89984 | 0.05746 |
| *Polyommatus icarus* : year | -0.08587 | 0.00952 | -9.0218 | 0 |
| *Pyrgus malvae* : year | -0.05127 | 0.02482 | -2.06541 | 0.03889 |
| *Pyronia tithonus* : year | -0.05359 | 0.01629 | -3.28993 | 0.001 |

***Supp. Table 6: The PC1 quantile regression model results for a τ value of 0.1.***

| **Term** | **Value** | **Std. Error** | **t value** | **P value** |
|---|---|---|---|---|
| *Aglais urticae* | -0.89269 | 0.03327 | -26.83105 | 0 |
| *Anthocharis cardamines* | -0.8335 | 0.04216 | -19.77041 | 0 |
| *Aphantopus hyperantus* | -0.76953 | 0.0353 | -21.79702 | 0 |
| *Argynnis adippe* | -1.05026 | 0.01684 | -62.36654 | 0 |
| *Argynnis aglaja* | -0.92879 | 0.03548 | -26.18068 | 0 |
| *Argynnis paphia* | -1.0331 | 0.01266 | -81.58397 | 0 |
| *Aricia agestis* | -1.09635 | 0.021 | -52.21904 | 0 |
| *Aricia artaxerxes* | -1.13973 | 0.03239 | -35.18624 | 0 |
| *Callophrys rubi* | -1.17256 | 0.01489 | -78.73733 | 0 |
| *Celastrina argiolus* | -1.27521 | 0.01438 | -88.6939 | 0 |
| *Coenonympha pamphilus* | -0.85306 | 0.01217 | -70.11008 | 0 |
| *Coenonympha tullia* | -0.67266 | 0.05072 | -13.26215 | 0 |
| *Colias croceus* | 0.47415 | 0.05142 | 9.22156 | 0 |
| *Cupido minimus* | -1.22336 | 0.01521 | -80.44164 | 0 |
| *Erebia aethiops* | -0.54848 | 0.22082 | -2.48378 | 0.013 |
| *Euphydryas aurinia* | -1.23406 | 0.00781 | -158.04892 | 0 |
| *Hipparchia semele* | -1.13345 | 0.01662 | -68.19711 | 0 |
| *Lasiommata megera* | -1.17098 | 0.00817 | -143.36783 | 0 |
| *Leptidea sinapis* | -1.66563 | 0.01404 | -118.64675 | 0 |
| *Lycaena phlaeas* | -0.55341 | 0.0499 | -11.08986 | 0 |
| *Lysandra bellargus* | -1.24048 | 0.00867 | -143.10123 | 0 |
| *Lysandra coridon* | -1.27989 | 0.00398 | -321.56808 | 0 |
| *Maculinea arion* | -1.06656 | 0.06938 | -15.37258 | 0 |
| *Maniola jurtina* | -0.93454 | 0.01423 | -65.68675 | 0 |
| *Melanargia galathea* | -1.24988 | 0.01325 | -94.30456 | 0 |
| *Melitaea athalia* | -1.23408 | 0.03394 | -36.36498 | 0 |
| *Pararge aegeria* | -0.83664 | 0.02707 | -30.90921 | 0 |
| *Pieris napi* | -1.34863 | 0.00778 | -173.36311 | 0 |
| *Pieris rapae* | -1.15055 | 0.01874 | -61.38317 | 0 |
| *Plebejus argus* | -1.25134 | 0.00762 | -164.26632 | 0 |
| *Polygonia c-album* | -1.25746 | 0.01403 | -89.64847 | 0 |
| *Polyommatus icarus* | -1.22386 | 0.00658 | -185.94204 | 0 |
| *Pyrgus malvae* | -1.3908 | 0.0143 | -97.24138 | 0 |
| *Pyronia tithonus* | -0.83206 | 0.02482 | -33.51826 | 0 |
| temperature : *Aglais urticae* | 0.03394 | 0.03834 | 0.88508 | 0.37612 |
| temperature : *Anthocharis cardamines* | 0.04601 | 0.03263 | 1.40987 | 0.15858 |
| temperature : *Aphantopus hyperantus* | -0.06246 | 0.03492 | -1.78853 | 0.07369 |
| temperature : *Argynnis adippe* | -0.01921 | 0.01968 | -0.9764 | 0.32887 |
| temperature : *Argynnis aglaja* | -0.00441 | 0.01749 | -0.2522 | 0.80089 |
| temperature : *Argynnis paphia* | -0.01151 | 0.02271 | -0.50697 | 0.61218 |
| temperature : *Aricia agestis* | 0.06444 | 0.02754 | 2.33929 | 0.01932 |
| temperature : *Aricia artaxerxes* | -0.01866 | 0.01363 | -1.36889 | 0.17104 |
| temperature : *Callophrys rubi* | -0.02 | 0.01843 | -1.08479 | 0.27802 |
| temperature : *Celastrina argiolus* | 0.00146 | 0.02054 | 0.07128 | 0.94318 |
| temperature : *Coenonympha pamphilus* | 0.00286 | 0.01287 | 0.22266 | 0.8238 |
| temperature : *Coenonympha tullia* | 0.02506 | 0.02153 | 1.16403 | 0.24442 |
| temperature : *Colias croceus* | -0.03102 | 0.05334 | -0.58165 | 0.56081 |
| temperature : *Cupido minimus* | 0.02525 | 0.01744 | 1.4475 | 0.14776 |
| temperature : *Erebia aethiops* | -0.02326 | 0.04196 | -0.5544 | 0.57931 |
| temperature : *Euphydryas aurinia* | 0.03035 | 0.00718 | 4.22877 | 2e-05 |
| temperature : *Hipparchia semele* | 0.03079 | 0.00872 | 3.52947 | 0.00042 |
| temperature : *Lasiommata megera* | -0.00667 | 0.00782 | -0.85249 | 0.39394 |
| temperature : *Leptidea sinapis* | -0.02425 | 0.02389 | -1.0151 | 0.31006 |
| temperature : *Lycaena phlaeas* | 0.03598 | 0.05227 | 0.68831 | 0.49126 |
| temperature : *Lysandra bellargus* | 0.01798 | 0.00949 | 1.89474 | 0.05813 |

| | | | | |
|---|---|---|---|---|
| temperature : *Lysandra coridon* | 0.0241 | 0.00673 | 3.58226 | 0.00034 |
| temperature : *Maculinea arion* | -0.16304 | 0.06857 | -2.37764 | 0.01743 |
| temperature : *Maniola jurtina* | 0.02681 | 0.0185 | 1.44919 | 0.14729 |
| temperature : *Melanargia galathea* | -0.0011 | 0.01807 | -0.06084 | 0.95148 |
| temperature : *Melitaea athalia* | -0.02765 | 0.03064 | -0.90255 | 0.36677 |
| temperature : *Pararge aegeria* | 0.02366 | 0.02676 | 0.88408 | 0.37665 |
| temperature : *Pieris napi* | -0.01063 | 0.00478 | -2.22132 | 0.02633 |
| temperature : *Pieris rapae* | 0.02755 | 0.01932 | 1.42564 | 0.15397 |
| temperature : *Plebejus argus* | -0.03806 | 0.00868 | -4.38405 | 1e-05 |
| temperature : *Polygonia c-album* | -0.01144 | 0.01866 | -0.61306 | 0.53984 |
| temperature : *Polyommatus icarus* | -0.01373 | 0.00591 | -2.32387 | 0.02013 |
| temperature : *Pyrgus malvae* | 0.05433 | 0.02529 | 2.14822 | 0.0317 |
| temperature : *Pyronia tithonus* | 0.03847 | 0.03337 | 1.15284 | 0.24898 |
| *Aglais urticae* : year | -0.08928 | 0.01895 | -4.71179 | 0 |
| *Anthocharis cardamines* : year | -0.06613 | 0.03127 | -2.1147 | 0.03446 |
| *Aphantopus hyperantus* : year | -0.02459 | 0.03203 | -0.76752 | 0.44278 |
| *Argynnis adippe* : year | -0.02251 | 0.01239 | -1.81621 | 0.06934 |
| *Argynnis aglaja* : year | -0.05978 | 0.0151 | -3.95771 | 8e-05 |
| *Argynnis paphia* : year | 0.00656 | 0.01193 | 0.54967 | 0.58254 |
| *Aricia agestis* : year | -0.01707 | 0.01848 | -0.92367 | 0.35566 |
| *Aricia artaxerxes* : year | -0.00974 | 0.01548 | -0.62904 | 0.52932 |
| *Callophrys rubi* : year | -0.00207 | 0.00837 | -0.24761 | 0.80444 |
| *Celastrina argiolus* : year | -0.00603 | 0.01297 | -0.46485 | 0.64204 |
| *Coenonympha pamphilus* : year | -0.00718 | 0.01174 | -0.61133 | 0.54098 |
| *Coenonympha tullia* : year | -0.0109 | 0.03668 | -0.29705 | 0.76643 |
| *Colias croceus* : year | -0.07493 | 0.02134 | -3.5113 | 0.00045 |
| *Cupido minimus* : year | -0.03821 | 0.0149 | -2.56469 | 0.01033 |
| *Erebia aethiops* : year | 0.20012 | 0.05211 | 3.84024 | 0.00012 |
| *Euphydryas aurinia* : year | -0.02307 | 0.00694 | -3.32289 | 0.00089 |
| *Hipparchia semele* : year | -0.01122 | 0.01143 | -0.98138 | 0.32641 |
| *Lasiommata megera* : year | -0.00326 | 0.00456 | -0.71357 | 0.47549 |
| *Leptidea sinapis* : year | 0.01808 | 0.01479 | 1.22219 | 0.22164 |
| *Lycaena phlaeas* : year | -0.18579 | 0.03765 | -4.93408 | 0 |
| *Lysandra bellargus* : year | 0.00282 | 0.0065 | 0.43408 | 0.66423 |
| *Lysandra coridon* : year | 0.01171 | 0.00455 | 2.57531 | 0.01002 |
| *Maculinea arion* : year | 0.1356 | 0.03154 | 4.2992 | 2e-05 |
| *Maniola jurtina* : year | 0.02228 | 0.0156 | 1.42785 | 0.15334 |
| *Melanargia galathea* : year | 0.00257 | 0.0092 | 0.27947 | 0.77989 |
| *Melitaea athalia* : year | 0.03261 | 0.02407 | 1.35498 | 0.17543 |
| *Pararge aegeria* : year | 0.00478 | 0.02021 | 0.2366 | 0.81297 |
| *Pieris napi* : year | 0.01969 | 0.00679 | 2.90008 | 0.00373 |
| *Pieris rapae* : year | 0.00604 | 0.01781 | 0.3392 | 0.73446 |
| *Plebejus argus* : year | -0.03925 | 0.0089 | -4.40864 | 1e-05 |
| *Polygonia c-album* : year | 0.01394 | 0.00989 | 1.40871 | 0.15892 |
| *Polyommatus icarus* : year | -0.0367 | 0.00744 | -4.93134 | 0 |
| *Pyrgus malvae* : year | -0.03161 | 0.0124 | -2.54849 | 0.01082 |
| *Pyronia tithonus* : year | -0.08312 | 0.01632 | -5.09198 | 0 |

*Supp. Table 7: The PC1 quantile regression model results for a τ value of 0.25.*

| Term | Value | Std. Error | t value | P value |
|---|---|---|---|---|
| *Aglais urticae* | 0.16256 | 0.06474 | 2.51107 | 0.01204 |
| *Anthocharis cardamines* | 0.00942 | 0.03328 | 0.28306 | 0.77713 |
| *Aphantopus hyperantus* | -0.02223 | 0.02185 | -1.01762 | 0.30886 |
| *Argynnis adippe* | -0.89965 | 0.01619 | -55.55558 | 0 |
| *Argynnis aglaja* | -0.20894 | 0.06303 | -3.31463 | 0.00092 |
| *Argynnis paphia* | -0.83002 | 0.01252 | -66.30356 | 0 |
| *Aricia agestis* | -0.73841 | 0.02192 | -33.6901 | 0 |
| *Aricia artaxerxes* | -0.72351 | 0.06386 | -11.32973 | 0 |
| *Callophrys rubi* | -1.02987 | 0.0085 | -121.2222 | 0 |
| *Celastrina argiolus* | -1.12098 | 0.01325 | -84.62476 | 0 |
| *Coenonympha pamphilus* | -0.49025 | 0.03223 | -15.21277 | 0 |
| *Coenonympha tullia* | 0.06602 | 0.01978 | 3.33783 | 0.00084 |
| *Colias croceus* | 1.04665 | 0.05253 | 19.92481 | 0 |
| *Cupido minimus* | -1.00576 | 0.01797 | -55.97259 | 0 |
| *Erebia aethiops* | 0.48951 | 0.10476 | 4.67282 | 0 |
| *Euphydryas aurinia* | -1.05554 | 0.00672 | -157.07623 | 0 |
| *Hipparchia semele* | -0.85237 | 0.02637 | -32.31931 | 0 |
| *Lasiommata megera* | -0.99854 | 0.01078 | -92.6712 | 0 |
| *Leptidea sinapis* | -1.45544 | 0.01973 | -73.77012 | 0 |
| *Lycaena phlaeas* | 0.21832 | 0.02904 | 7.51828 | 0 |
| *Lysandra bellargus* | -1.03971 | 0.00898 | -115.78845 | 0 |
| *Lysandra coridon* | -1.02264 | 0.00521 | -196.4395 | 0 |
| *Maculinea arion* | -0.3457 | 0.15924 | -2.17088 | 0.02994 |
| *Maniola jurtina* | -0.21206 | 0.02398 | -8.84241 | 0 |
| *Melanargia galathea* | -1.09219 | 0.00833 | -131.09214 | 0 |
| *Melitaea athalia* | -0.95474 | 0.04101 | -23.27804 | 0 |
| *Pararge aegeria* | 0.11315 | 0.03509 | 3.22475 | 0.00126 |
| *Pieris napi* | -1.20532 | 0.00765 | -157.5573 | 0 |
| *Pieris rapae* | -0.80184 | 0.0269 | -29.80846 | 0 |
| *Plebejus argus* | -0.98107 | 0.00747 | -131.40737 | 0 |
| *Polygonia c-album* | -1.12619 | 0.01013 | -111.17875 | 0 |
| *Polyommatus icarus* | -0.98102 | 0.00633 | -154.91237 | 0 |
| *Pyrgus malvae* | -1.03544 | 0.02445 | -42.34819 | 0 |
| *Pyronia tithonus* | -0.05549 | 0.03638 | -1.52546 | 0.12715 |
| temperature : *Aglais urticae* | 0.06019 | 0.08737 | 0.68886 | 0.49091 |
| temperature : *Anthocharis cardamines* | 0.11955 | 0.03022 | 3.95631 | 8e-05 |
| temperature : *Aphantopus hyperantus* | -0.0459 | 0.01775 | -2.58641 | 0.0097 |
| temperature : *Argynnis adippe* | -0.02152 | 0.01658 | -1.29791 | 0.19432 |
| temperature : *Argynnis aglaja* | 0.03105 | 0.03004 | 1.03368 | 0.30129 |
| temperature : *Argynnis paphia* | -0.00845 | 0.02167 | -0.39001 | 0.69653 |
| temperature : *Aricia agestis* | 0.10581 | 0.01548 | 6.83598 | 0 |
| temperature : *Aricia artaxerxes* | -0.0086 | 0.02897 | -0.29677 | 0.76664 |
| temperature : *Callophrys rubi* | -0.01981 | 0.01039 | -1.9068 | 0.05655 |
| temperature : *Celastrina argiolus* | 0.02492 | 0.01913 | 1.30277 | 0.19266 |
| temperature : *Coenonympha pamphilus* | 0.00244 | 0.01807 | 0.13525 | 0.89242 |
| temperature : *Coenonympha tullia* | 0.02771 | 0.01203 | 2.30425 | 0.02121 |
| temperature : *Colias croceus* | -0.0297 | 0.05561 | -0.53405 | 0.59331 |
| temperature : *Cupido minimus* | 0.01878 | 0.00918 | 2.04597 | 0.04076 |
| temperature : *Erebia aethiops* | -0.00377 | 0.03526 | -0.10705 | 0.91475 |
| temperature : *Euphydryas aurinia* | 0.02187 | 0.00729 | 3.00171 | 0.00269 |
| temperature : *Hipparchia semele* | 0.02992 | 0.02265 | 1.32107 | 0.18648 |
| temperature : *Lasiommata megera* | -0.00927 | 0.01477 | -0.62759 | 0.53028 |
| temperature : *Leptidea sinapis* | -0.03952 | 0.03234 | -1.22211 | 0.22167 |
| temperature : *Lycaena phlaeas* | 0.05825 | 0.03253 | 1.79105 | 0.07329 |
| temperature : *Lysandra bellargus* | 0.0157 | 0.01008 | 1.55702 | 0.11947 |

| | | | | |
|---|---|---|---|---|
| temperature : *Lysandra coridon* | 0.03285 | 0.00825 | 3.98 | 7e-05 |
| temperature : *Maculinea arion* | -0.28442 | 0.17858 | -1.5927 | 0.11123 |
| temperature : *Maniola jurtina* | 0.05707 | 0.02777 | 2.05495 | 0.03989 |
| temperature : *Melanargia galathea* | 0.02193 | 0.01193 | 1.83806 | 0.06606 |
| temperature : *Melitaea athalia* | -0.05247 | 0.03129 | -1.6772 | 0.09351 |
| temperature : *Pararge aegeria* | 0.07005 | 0.0303 | 2.31229 | 0.02076 |
| temperature : *Pieris napi* | -0.02179 | 0.00451 | -4.82666 | 0 |
| temperature : *Pieris rapae* | -0.01265 | 0.03176 | -0.39829 | 0.69042 |
| temperature : *Plebejus argus* | -0.02071 | 0.01043 | -1.98552 | 0.04709 |
| temperature : *Polygonia c-album* | 0.02323 | 0.01737 | 1.33711 | 0.18119 |
| temperature : *Polyommatus icarus* | -0.0164 | 0.00662 | -2.47686 | 0.01326 |
| temperature : *Pyrgus malvae* | -0.02592 | 0.04234 | -0.61235 | 0.54031 |
| temperature : *Pyronia tithonus* | 0.01103 | 0.05282 | 0.20886 | 0.83456 |
| *Aglais urticae* : year | -0.34616 | 0.05575 | -6.20881 | 0 |
| *Anthocharis cardamines* : year | -0.05413 | 0.02881 | -1.87847 | 0.06032 |
| *Aphantopus hyperantus* : year | -0.05473 | 0.02127 | -2.57317 | 0.01008 |
| *Argynnis adippe* : year | -0.02771 | 0.00932 | -2.97179 | 0.00296 |
| *Argynnis aglaja* : year | -0.23632 | 0.05336 | -4.42841 | 1e-05 |
| *Argynnis paphia* : year | 0.01896 | 0.01189 | 1.59364 | 0.11102 |
| *Aricia agestis* : year | -0.04126 | 0.01643 | -2.51195 | 0.01201 |
| *Aricia artaxerxes* : year | -0.04053 | 0.02886 | -1.40397 | 0.16033 |
| *Callophrys rubi* : year | -0.02465 | 0.00646 | -3.81525 | 0.00014 |
| *Celastrina argiolus* : year | 0.00937 | 0.01076 | 0.87118 | 0.38366 |
| *Coenonympha pamphilus* : year | 0.0237 | 0.02552 | 0.92836 | 0.35323 |
| *Coenonympha tullia* : year | -0.04847 | 0.01282 | -3.78156 | 0.00016 |
| *Colias croceus* : year | 0.01218 | 0.04008 | 0.30397 | 0.76115 |
| *Cupido minimus* : year | -0.03746 | 0.01723 | -2.17389 | 0.02972 |
| *Erebia aethiops* : year | 0.10645 | 0.04767 | 2.23306 | 0.02555 |
| *Euphydryas aurinia* : year | -0.03304 | 0.00551 | -5.99252 | 0 |
| *Hipparchia semele* : year | -0.02537 | 0.0207 | -1.2255 | 0.22039 |
| *Lasiommata megera* : year | -0.02297 | 0.01033 | -2.22291 | 0.02622 |
| *Leptidea sinapis* : year | 0.00351 | 0.0181 | 0.19377 | 0.84636 |
| *Lycaena phlaeas* : year | -0.14446 | 0.02575 | -5.61049 | 0 |
| *Lysandra bellargus* : year | -0.0016 | 0.00729 | -0.22008 | 0.82581 |
| *Lysandra coridon* : year | -0.01276 | 0.0056 | -2.27657 | 0.02281 |
| *Maculinea arion* : year | 0.21735 | 0.08096 | 2.68473 | 0.00726 |
| *Maniola jurtina* : year | 0.04934 | 0.0227 | 2.17387 | 0.02972 |
| *Melanargia galathea* : year | -0.01592 | 0.00597 | -2.6669 | 0.00766 |
| *Melitaea athalia* : year | 0.04339 | 0.02437 | 1.78018 | 0.07505 |
| *Pararge aegeria* : year | -0.00025 | 0.0279 | -0.00912 | 0.99272 |
| *Pieris napi* : year | 0.01952 | 0.00554 | 3.52282 | 0.00043 |
| *Pieris rapae* : year | 0.05008 | 0.02582 | 1.93979 | 0.05241 |
| *Plebejus argus* : year | -0.05027 | 0.00793 | -6.34068 | 0 |
| *Polygonia c-album* : year | -0.00575 | 0.01026 | -0.56022 | 0.57533 |
| *Polyommatus icarus* : year | -0.03958 | 0.00683 | -5.7932 | 0 |
| *Pyrgus malvae* : year | -0.06175 | 0.02354 | -2.62327 | 0.00871 |
| *Pyronia tithonus* : year | -0.05838 | 0.03001 | -1.94524 | 0.05175 |

***Supp. Table 8: The PC1 quantile regression model results for a τ value of 0.5.***

| **Term** | **Value** | **Std. Error** | **t value** | **P value** |
|---|---|---|---|---|
| *Aglais urticae* | 1.02157 | 0.04048 | 25.2343 | 0 |
| *Anthocharis cardamines* | 0.73399 | 0.02462 | 29.81399 | 0 |
| *Aphantopus hyperantus* | 0.6347 | 0.03301 | 19.22654 | 0 |
| *Argynnis adippe* | -0.69328 | 0.01177 | -58.88351 | 0 |
| *Argynnis aglaja* | 0.62777 | 0.03821 | 16.42761 | 0 |
| *Argynnis paphia* | -0.25191 | 0.07094 | -3.55096 | 0.00038 |
| *Aricia agestis* | 0.25406 | 0.0442 | 5.74847 | 0 |
| *Aricia artaxerxes* | 0.51646 | 0.09006 | 5.73464 | 0 |
| *Callophrys rubi* | -0.86469 | 0.01573 | -54.95967 | 0 |
| *Celastrina argiolus* | -0.94564 | 0.01172 | -80.6662 | 0 |
| *Coenonympha pamphilus* | 0.42575 | 0.02146 | 19.83459 | 0 |
| *Coenonympha tullia* | 0.69346 | 0.02988 | 23.21034 | 0 |
| *Colias croceus* | 1.52008 | 0.02387 | 63.68935 | 0 |
| *Cupido minimus* | -0.50339 | 0.04071 | -12.36542 | 0 |
| *Erebia aethiops* | 1.34152 | 0.06252 | 21.45709 | 0 |
| *Euphydryas aurinia* | -0.78067 | 0.01259 | -61.99663 | 0 |
| *Hipparchia semele* | 0.11313 | 0.03009 | 3.75944 | 0.00017 |
| *Lasiommata megera* | -0.73617 | 0.01567 | -46.98831 | 0 |
| *Leptidea sinapis* | -1.03317 | 0.02645 | -39.05392 | 0 |
| *Lycaena phlaeas* | 1.00206 | 0.0221 | 45.35216 | 0 |
| *Lysandra bellargus* | -0.74998 | 0.0114 | -65.79873 | 0 |
| *Lysandra coridon* | -0.21091 | 0.02137 | -9.87074 | 0 |
| *Maculinea arion* | 0.31599 | 0.07342 | 4.30355 | 2e-05 |
| *Maniola jurtina* | 0.46238 | 0.02057 | 22.47881 | 0 |
| *Melanargia galathea* | -0.87737 | 0.01291 | -67.9652 | 0 |
| *Melitaea athalia* | -0.20893 | 0.10733 | -1.94674 | 0.05157 |
| *Pararge aegeria* | 0.66738 | 0.04029 | 16.56497 | 0 |
| *Pieris napi* | -0.99755 | 0.00814 | -122.57825 | 0 |
| *Pieris rapae* | 0.22054 | 0.04032 | 5.46945 | 0 |
| *Plebejus argus* | -0.43175 | 0.02231 | -19.3513 | 0 |
| *Polygonia c-album* | -0.46961 | 0.13039 | -3.6015 | 0.00032 |
| *Polyommatus icarus* | -0.47449 | 0.01794 | -26.44807 | 0 |
| *Pyrgus malvae* | -0.11249 | 0.06231 | -1.80543 | 0.07101 |
| *Pyronia tithonus* | 0.55151 | 0.02959 | 18.63789 | 0 |
| temperature : *Aglais urticae* | 0.03886 | 0.03578 | 1.08585 | 0.27755 |
| temperature : *Anthocharis cardamines* | 0.0951 | 0.03252 | 2.9245 | 0.00345 |
| temperature : *Aphantopus hyperantus* | -0.0887 | 0.02715 | -3.26714 | 0.00109 |
| temperature : *Argynnis adippe* | -0.02593 | 0.01324 | -1.95843 | 0.05018 |
| temperature : *Argynnis aglaja* | 0.01279 | 0.01117 | 1.14446 | 0.25244 |
| temperature : *Argynnis paphia* | -0.09998 | 0.08902 | -1.12309 | 0.2614 |
| temperature : *Aricia agestis* | 0.20769 | 0.06014 | 3.45372 | 0.00055 |
| temperature : *Aricia artaxerxes* | 0.04707 | 0.05388 | 0.87356 | 0.38236 |
| temperature : *Callophrys rubi* | -0.01634 | 0.01199 | -1.36273 | 0.17297 |
| temperature : *Celastrina argiolus* | 0.03266 | 0.01862 | 1.75388 | 0.07945 |
| temperature : *Coenonympha pamphilus* | 0.00673 | 0.01141 | 0.59045 | 0.55489 |
| temperature : *Coenonympha tullia* | 0.03004 | 0.01338 | 2.24464 | 0.02479 |
| temperature : *Colias croceus* | -0.05539 | 0.02359 | -2.34778 | 0.01889 |
| temperature : *Cupido minimus* | 0.07512 | 0.0434 | 1.73084 | 0.08348 |
| temperature : *Erebia aethiops* | 0.01266 | 0.02212 | 0.57236 | 0.56708 |
| temperature : *Euphydryas aurinia* | 0.00411 | 0.0141 | 0.29141 | 0.77074 |
| temperature : *Hipparchia semele* | 0.06355 | 0.03563 | 1.78346 | 0.07451 |
| temperature : *Lasiommata megera* | -0.03552 | 0.01519 | -2.33835 | 0.01937 |
| temperature : *Leptidea sinapis* | -0.11846 | 0.04508 | -2.62774 | 0.0086 |
| temperature : *Lycaena phlaeas* | 0.03035 | 0.02686 | 1.13001 | 0.25847 |
| temperature : *Lysandra bellargus* | -8.00E-05 | 0.0138 | -0.00615 | 0.99509 |

| | | | | |
|---|---|---|---|---|
| temperature : *Lysandra coridon* | 0.09 | 0.03406 | 2.64231 | 0.00824 |
| temperature : *Maculinea arion* | -0.04465 | 0.1059 | -0.42164 | 0.67329 |
| temperature : *Maniola jurtina* | 0.09522 | 0.01882 | 5.06004 | 0 |
| temperature : *Melanargia galathea* | 0.02233 | 0.01863 | 1.19872 | 0.23064 |
| temperature : *Melitaea athalia* | -0.22443 | 0.06232 | -3.60096 | 0.00032 |
| temperature : *Pararge aegeria* | 0.10285 | 0.04723 | 2.17774 | 0.02943 |
| temperature : *Pieris napi* | -0.02032 | 0.00633 | -3.21086 | 0.00132 |
| temperature : *Pieris rapae* | 0.03428 | 0.03206 | 1.06915 | 0.285 |
| temperature : *Plebejus argus* | 0.01784 | 0.02755 | 0.64744 | 0.51735 |
| temperature : *Polygonia c-album* | 0.39824 | 0.14104 | 2.82354 | 0.00475 |
| temperature : *Polyommatus icarus* | -0.04913 | 0.01879 | -2.61442 | 0.00894 |
| temperature : *Pyrgus malvae* | -0.05537 | 0.09596 | -0.57701 | 0.56394 |
| temperature : *Pyronia tithonus* | 0.06981 | 0.04534 | 1.53963 | 0.12365 |
| *Aglais urticae* : year | -0.20172 | 0.01892 | -10.66196 | 0 |
| *Anthocharis cardamines* : year | -0.02822 | 0.01869 | -1.51036 | 0.13096 |
| *Aphantopus hyperantus* : year | -0.09183 | 0.03262 | -2.8154 | 0.00487 |
| *Argynnis adippe* : year | -0.06575 | 0.00874 | -7.51884 | 0 |
| *Argynnis aglaja* : year | -0.08292 | 0.026 | -3.18867 | 0.00143 |
| *Argynnis paphia* : year | 0.20383 | 0.06247 | 3.26283 | 0.0011 |
| *Aricia agestis* : year | -0.17292 | 0.04151 | -4.16604 | 3e-05 |
| *Aricia artaxerxes* : year | -0.10639 | 0.05544 | -1.91906 | 0.05498 |
| *Callophrys rubi* : year | -0.04304 | 0.01595 | -2.69825 | 0.00697 |
| *Celastrina argiolus* : year | -0.00949 | 0.01015 | -0.93512 | 0.34973 |
| *Coenonympha pamphilus* : year | -0.02464 | 0.01715 | -1.43659 | 0.15084 |
| *Coenonympha tullia* : year | -0.04826 | 0.01948 | -2.47725 | 0.01324 |
| *Colias croceus* : year | 0.00299 | 0.01725 | 0.17364 | 0.86215 |
| *Cupido minimus* : year | -0.13583 | 0.03508 | -3.87137 | 0.00011 |
| *Erebia aethiops* : year | 0.01457 | 0.02253 | 0.64645 | 0.51799 |
| *Euphydryas aurinia* : year | -0.03518 | 0.01156 | -3.04405 | 0.00233 |
| *Hipparchia semele* : year | -0.08857 | 0.0251 | -3.52797 | 0.00042 |
| *Lasiommata megera* : year | -0.05435 | 0.0095 | -5.72353 | 0 |
| *Leptidea sinapis* : year | -0.02247 | 0.02792 | -0.80485 | 0.42091 |
| *Lycaena phlaeas* : year | -0.11876 | 0.022 | -5.39899 | 0 |
| *Lysandra bellargus* : year | 0.00215 | 0.00938 | 0.22943 | 0.81854 |
| *Lysandra coridon* : year | -0.12006 | 0.0232 | -5.17484 | 0 |
| *Maculinea arion* : year | 0.1059 | 0.04168 | 2.54107 | 0.01105 |
| *Maniola jurtina* : year | 0.0505 | 0.01976 | 2.55575 | 0.0106 |
| *Melanargia galathea* : year | -0.02759 | 0.00941 | -2.93189 | 0.00337 |
| *Melitaea athalia* : year | 0.17463 | 0.06777 | 2.57673 | 0.00998 |
| *Pararge aegeria* : year | -0.00752 | 0.03305 | -0.22746 | 0.82007 |
| *Pieris napi* : year | 0.02051 | 0.00715 | 2.86894 | 0.00412 |
| *Pieris rapae* : year | 0.0692 | 0.02043 | 3.38731 | 0.00071 |
| *Plebejus argus* : year | -0.15216 | 0.01891 | -8.04513 | 0 |
| *Polygonia c-album* : year | -0.14659 | 0.06911 | -2.12095 | 0.03393 |
| *Polyommatus icarus* : year | -0.11227 | 0.01647 | -6.81747 | 0 |
| *Pyrgus malvae* : year | -0.05964 | 0.05926 | -1.00642 | 0.31422 |
| *Pyronia tithonus* : year | -0.07179 | 0.02288 | -3.13795 | 0.0017 |

***Supp. Table 9: The PC1 quantile regression model results for a τ value of 0.75.***

| **Term** | **Value** | **Std. Error** | **t value** | **P value** |
|---|---|---|---|---|
| *Aglais urticae* | 1.74799 | 0.01158 | 150.91297 | 0 |
| *Anthocharis cardamines* | 1.1929 | 0.01976 | 60.35644 | 0 |
| *Aphantopus hyperantus* | 1.16412 | 0.01454 | 80.05247 | 0 |
| *Argynnis adippe* | -0.30039 | 0.04314 | -6.96226 | 0 |
| *Argynnis aglaja* | 1.462 | 0.03476 | 42.06489 | 0 |
| *Argynnis paphia* | 0.76153 | 0.02905 | 26.21896 | 0 |
| *Aricia agestis* | 1.02562 | 0.03538 | 28.98875 | 0 |
| *Aricia artaxerxes* | 1.19154 | 0.08154 | 14.6136 | 0 |
| *Callophrys rubi* | -0.64828 | 0.02186 | -29.66005 | 0 |
| *Celastrina argiolus* | -0.71349 | 0.01703 | -41.885 | 0 |
| *Coenonympha pamphilus* | 1.26574 | 0.02408 | 52.56621 | 0 |
| *Coenonympha tullia* | 1.19674 | 0.01367 | 87.56241 | 0 |
| *Colias croceus* | 1.67533 | 0.00401 | 418.1392 | 0 |
| *Cupido minimus* | 0.41007 | 0.02732 | 15.00882 | 0 |
| *Erebia aethiops* | 1.6778 | 0.01767 | 94.94794 | 0 |
| *Euphydryas aurinia* | 0.20679 | 0.06082 | 3.39999 | 0.00067 |
| *Hipparchia semele* | 0.72648 | 0.02922 | 24.86462 | 0 |
| *Lasiommata megera* | 0.14279 | 0.07129 | 2.0028 | 0.0452 |
| *Leptidea sinapis* | -0.56173 | 0.03414 | -16.455 | 0 |
| *Lycaena phlaeas* | 1.5162 | 0.01253 | 120.9983 | 0 |
| *Lysandra bellargus* | -0.10486 | 0.03645 | -2.87655 | 0.00402 |
| *Lysandra coridon* | 0.63473 | 0.01038 | 61.16117 | 0 |
| *Maculinea arion* | 1.13144 | 0.06461 | 17.51139 | 0 |
| *Maniola jurtina* | 1.23405 | 0.01048 | 117.71135 | 0 |
| *Melanargia galathea* | -0.33787 | 0.04465 | -7.5673 | 0 |
| *Melitaea athalia* | 0.83004 | 0.07989 | 10.38917 | 0 |
| *Pararge aegeria* | 1.4578 | 0.01655 | 88.06124 | 0 |
| *Pieris napi* | -0.69682 | 0.01681 | -41.44863 | 0 |
| *Pieris rapae* | 1.02147 | 0.04315 | 23.67209 | 0 |
| *Plebejus argus* | 0.5164 | 0.01519 | 34.00378 | 0 |
| *Polygonia c-album* | 0.59918 | 0.06008 | 9.97371 | 0 |
| *Polyommatus icarus* | 0.44994 | 0.01242 | 36.2411 | 0 |
| *Pyrgus malvae* | 0.59684 | 0.05529 | 10.79523 | 0 |
| *Pyronia tithonus* | 1.27283 | 0.01496 | 85.09271 | 0 |
| temperature : *Aglais urticae* | 0.00534 | 0.01172 | 0.45604 | 0.64837 |
| temperature : *Anthocharis cardamines* | 0.07019 | 0.02528 | 2.77663 | 0.00549 |
| temperature : *Aphantopus hyperantus* | -0.03882 | 0.01178 | -3.29469 | 0.00099 |
| temperature : *Argynnis adippe* | 0.00262 | 0.06214 | 0.04222 | 0.96632 |
| temperature : *Argynnis aglaja* | 0.03774 | 0.03194 | 1.18148 | 0.23742 |
| temperature : *Argynnis paphia* | 0.01698 | 0.05457 | 0.31112 | 0.75571 |
| temperature : *Aricia agestis* | 0.1287 | 0.04968 | 2.59072 | 0.00958 |
| temperature : *Aricia artaxerxes* | -0.00708 | 0.0373 | -0.18978 | 0.84948 |
| temperature : *Callophrys rubi* | -0.01794 | 0.00829 | -2.16539 | 0.03036 |
| temperature : *Celastrina argiolus* | 0.00217 | 0.02783 | 0.07808 | 0.93776 |
| temperature : *Coenonympha pamphilus* | 0.02601 | 0.02876 | 0.90427 | 0.36585 |
| temperature : *Coenonympha tullia* | 0.02786 | 0.00854 | 3.26225 | 0.00111 |
| temperature : *Colias croceus* | -0.02011 | 0.008 | -2.5151 | 0.0119 |
| temperature : *Cupido minimus* | 0.13733 | 0.02087 | 6.58095 | 0 |
| temperature : *Erebia aethiops* | 0.01605 | 0.00864 | 1.85737 | 0.06326 |
| temperature : *Euphydryas aurinia* | -0.02426 | 0.0639 | -0.37958 | 0.70426 |
| temperature : *Hipparchia semele* | -0.01498 | 0.03221 | -0.46522 | 0.64178 |
| temperature : *Lasiommata megera* | -0.19925 | 0.06684 | -2.98104 | 0.00287 |
| temperature : *Leptidea sinapis* | -0.12213 | 0.02728 | -4.47659 | 1e-05 |
| temperature : *Lycaena phlaeas* | 0.02085 | 0.01427 | 1.46096 | 0.14403 |
| temperature : *Lysandra bellargus* | 0.05509 | 0.04687 | 1.17523 | 0.23991 |

| | | | | |
|---|---|---|---|---|
| temperature : *Lysandra coridon* | 0.06106 | 0.01556 | 3.92498 | 9e-05 |
| temperature : *Maculinea arion* | 0.00873 | 0.06948 | 0.12564 | 0.90002 |
| temperature : *Maniola jurtina* | 0.04488 | 0.01328 | 3.37992 | 0.00073 |
| temperature : *Melanargia galathea* | 0.09716 | 0.078 | 1.24561 | 0.21291 |
| temperature : *Melitaea athalia* | -0.21026 | 0.0752 | -2.79589 | 0.00518 |
| temperature : *Pararge aegeria* | 0.05464 | 0.01831 | 2.98463 | 0.00284 |
| temperature : *Pieris napi* | -0.03109 | 0.01317 | -2.36029 | 0.01826 |
| temperature : *Pieris rapae* | 0.028 | 0.04959 | 0.56459 | 0.57235 |
| temperature : *Plebejus argus* | 0.05509 | 0.02076 | 2.65316 | 0.00798 |
| temperature : *Polygonia c-album* | 0.18973 | 0.07079 | 2.68012 | 0.00736 |
| temperature : *Polyommatus icarus* | -0.04086 | 0.01185 | -3.44695 | 0.00057 |
| temperature : *Pyrgus malvae* | -0.07072 | 0.066 | -1.0716 | 0.2839 |
| temperature : *Pyronia tithonus* | 0.03499 | 0.0215 | 1.62744 | 0.10365 |
| *Aglais urticae* : year | -0.00646 | 0.01052 | -0.61414 | 0.53912 |
| *Anthocharis cardamines* : year | -0.00213 | 0.0156 | -0.13672 | 0.89125 |
| *Aphantopus hyperantus* : year | -0.04484 | 0.01544 | -2.90298 | 0.0037 |
| *Argynnis adippe* : year | -0.11702 | 0.0437 | -2.67796 | 0.00741 |
| *Argynnis aglaja* : year | -0.02964 | 0.0317 | -0.93517 | 0.3497 |
| *Argynnis paphia* : year | -0.01489 | 0.0256 | -0.5816 | 0.56083 |
| *Aricia agestis* : year | -0.17537 | 0.03473 | -5.04993 | 0 |
| *Aricia artaxerxes* : year | -0.17072 | 0.0492 | -3.47009 | 0.00052 |
| *Callophrys rubi* : year | -0.06298 | 0.02071 | -3.04067 | 0.00236 |
| *Celastrina argiolus* : year | -0.01678 | 0.01528 | -1.09807 | 0.27218 |
| *Coenonympha pamphilus* : year | -0.04526 | 0.02161 | -2.09379 | 0.03628 |
| *Coenonympha tullia* : year | -0.04798 | 0.01083 | -4.43206 | 1e-05 |
| *Colias croceus* : year | -0.01307 | 0.00635 | -2.059 | 0.0395 |
| *Cupido minimus* : year | -0.09223 | 0.02006 | -4.59897 | 0 |
| *Erebia aethiops* : year | -0.00284 | 0.00742 | -0.38281 | 0.70186 |
| *Euphydryas aurinia* : year | -0.06532 | 0.05609 | -1.16456 | 0.2442 |
| *Hipparchia semele* : year | -0.0855 | 0.0215 | -3.97776 | 7e-05 |
| *Lasiommata megera* : year | -0.18374 | 0.05046 | -3.64132 | 0.00027 |
| *Leptidea sinapis* : year | -0.10445 | 0.02005 | -5.21008 | 0 |
| *Lycaena phlaeas* : year | -0.03327 | 0.01177 | -2.82747 | 0.00469 |
| *Lysandra bellargus* : year | 0.03221 | 0.03249 | 0.9915 | 0.32144 |
| *Lysandra coridon* : year | -0.08393 | 0.01 | -8.39176 | 0 |
| *Maculinea arion* : year | 0.04085 | 0.03251 | 1.25649 | 0.20894 |
| *Maniola jurtina* : year | -0.01102 | 0.00961 | -1.14661 | 0.25154 |
| *Melanargia galathea* : year | -0.00928 | 0.04171 | -0.22241 | 0.824 |
| *Melitaea athalia* : year | 0.1286 | 0.05558 | 2.31401 | 0.02067 |
| *Pararge aegeria* : year | 0.00223 | 0.01405 | 0.15847 | 0.87408 |
| *Pieris napi* : year | 0.04928 | 0.01698 | 2.90215 | 0.00371 |
| *Pieris rapae* : year | -0.06193 | 0.04274 | -1.44905 | 0.14733 |
| *Plebejus argus* : year | -0.1406 | 0.01749 | -8.04131 | 0 |
| *Polygonia c-album* : year | -0.11865 | 0.0523 | -2.26843 | 0.0233 |
| *Polyommatus icarus* : year | -0.12066 | 0.01421 | -8.49138 | 0 |
| *Pyrgus malvae* : year | -0.05263 | 0.04406 | -1.1946 | 0.23225 |
| *Pyronia tithonus* : year | -0.03235 | 0.01147 | -2.81955 | 0.00481 |

*Supp. Table 10: The PC1 quantile regression model results for a τ value of 0.9.*

| Term | Value | Std. Error | t value | P value |
|---|---|---|---|---|
| *Aglais urticae* | 1.87205 | 0.00867 | 215.93496 | 0 |
| *Anthocharis cardamines* | 1.47593 | 0.01411 | 104.5926 | 0 |
| *Aphantopus hyperantus* | 1.41253 | 0.00872 | 161.99407 | 0 |
| *Argynnis adippe* | 0.69682 | 0.11816 | 5.89725 | 0 |
| *Argynnis aglaja* | 1.83816 | 0.01804 | 101.87678 | 0 |
| *Argynnis paphia* | 1.69684 | 0.05615 | 30.21819 | 0 |
| *Aricia agestis* | 1.62203 | 0.02135 | 75.97151 | 0 |
| *Aricia artaxerxes* | 1.7517 | 0.02663 | 65.76733 | 0 |
| *Callophrys rubi* | 0.09 | 0.06912 | 1.30221 | 0.19285 |
| *Celastrina argiolus* | -0.28712 | 0.05739 | -5.00281 | 0 |
| *Coenonympha pamphilus* | 1.59494 | 0.01055 | 151.18038 | 0 |
| *Coenonympha tullia* | 1.4015 | 0.01138 | 123.10193 | 0 |
| *Colias croceus* | 1.78227 | 0.01302 | 136.91203 | 0 |
| *Cupido minimus* | 1.01878 | 0.07097 | 14.35536 | 0 |
| *Erebia aethiops* | 1.82138 | 0.021 | 86.72739 | 0 |
| *Euphydryas aurinia* | 0.98832 | 0.032 | 30.88913 | 0 |
| *Hipparchia semele* | 1.43745 | 0.03084 | 46.60421 | 0 |
| *Lasiommata megera* | 0.79823 | 0.05389 | 14.81156 | 0 |
| *Leptidea sinapis* | -0.07147 | 0.06344 | -1.12664 | 0.2599 |
| *Lycaena phlaeas* | 1.73339 | 0.00617 | 280.90447 | 0 |
| *Lysandra bellargus* | 0.5273 | 0.02807 | 18.78473 | 0 |
| *Lysandra coridon* | 1.30999 | 0.01704 | 76.8897 | 0 |
| *Maculinea arion* | 1.42542 | 0.05189 | 27.47096 | 0 |
| *Maniola jurtina* | 1.44871 | 0.00818 | 177.17678 | 0 |
| *Melanargia galathea* | 0.64643 | 0.03398 | 19.02231 | 0 |
| *Melitaea athalia* | 1.38287 | 0.13548 | 10.20725 | 0 |
| *Pararge aegeria* | 1.63187 | 0.01513 | 107.84353 | 0 |
| *Pieris napi* | 0.32584 | 0.04976 | 6.54771 | 0 |
| *Pieris rapae* | 1.54522 | 0.02225 | 69.45623 | 0 |
| *Plebejus argus* | 1.27773 | 0.02386 | 53.54506 | 0 |
| *Polygonia c-album* | 1.43282 | 0.04809 | 29.79311 | 0 |
| *Polyommatus icarus* | 1.00024 | 0.02012 | 49.72067 | 0 |
| *Pyrgus malvae* | 1.17335 | 0.02387 | 49.16052 | 0 |
| *Pyronia tithonus* | 1.48948 | 0.00993 | 150.04335 | 0 |
| temperature : *Aglais urticae* | 0.00438 | 0.01147 | 0.38149 | 0.70284 |
| temperature : *Anthocharis cardamines* | 0.04525 | 0.01426 | 3.17425 | 0.0015 |
| temperature : *Aphantopus hyperantus* | -0.02904 | 0.00633 | -4.58391 | 0 |
| temperature : *Argynnis adippe* | 0.06152 | 0.11181 | 0.55021 | 0.58218 |
| temperature : *Argynnis aglaja* | 0.01416 | 0.00444 | 3.18723 | 0.00144 |
| temperature : *Argynnis paphia* | -0.01512 | 0.06469 | -0.2337 | 0.81522 |
| temperature : *Aricia agestis* | 0.08019 | 0.0285 | 2.81352 | 0.0049 |
| temperature : *Aricia artaxerxes* | 0.01647 | 0.01083 | 1.52057 | 0.12837 |
| temperature : *Callophrys rubi* | -0.04964 | 0.07201 | -0.68936 | 0.4906 |
| temperature : *Celastrina argiolus* | 0.03074 | 0.12527 | 0.24539 | 0.80615 |
| temperature : *Coenonympha pamphilus* | 0.01166 | 0.0082 | 1.42288 | 0.15477 |
| temperature : *Coenonympha tullia* | 0.02019 | 0.00527 | 3.82754 | 0.00013 |
| temperature : *Colias croceus* | -0.00574 | 0.01413 | -0.40629 | 0.68453 |
| temperature : *Cupido minimus* | 0.1162 | 0.0863 | 1.34655 | 0.17813 |
| temperature : *Erebia aethiops* | 0.01282 | 0.01004 | 1.27623 | 0.20188 |
| temperature : *Euphydryas aurinia* | 0.07024 | 0.03297 | 2.13027 | 0.03315 |
| temperature : *Hipparchia semele* | -0.01571 | 0.0341 | -0.46065 | 0.64505 |
| temperature : *Lasiommata megera* | -0.21035 | 0.07293 | -2.88436 | 0.00392 |
| temperature : *Leptidea sinapis* | -0.2136 | 0.05632 | -3.79278 | 0.00015 |
| temperature : *Lycaena phlaeas* | 0.00947 | 0.00697 | 1.35932 | 0.17405 |
| temperature : *Lysandra bellargus* | 0.04998 | 0.02961 | 1.68781 | 0.09145 |

| | | | | |
|---|---|---|---|---|
| temperature : *Lysandra coridon* | 0.08283 | 0.027 | 3.06803 | 0.00216 |
| temperature : *Maculinea arion* | -0.01824 | 0.0588 | -0.31024 | 0.75638 |
| temperature : *Maniola jurtina* | 0.03333 | 0.00901 | 3.70097 | 0.00021 |
| temperature : *Melanargia galathea* | 0.01292 | 0.05178 | 0.24953 | 0.80295 |
| temperature : *Melitaea athalia* | -0.08067 | 0.11003 | -0.73323 | 0.46342 |
| temperature : *Pararge aegeria* | 0.04337 | 0.01724 | 2.51563 | 0.01188 |
| temperature : *Pieris napi* | -0.04446 | 0.03399 | -1.30815 | 0.19083 |
| temperature : *Pieris rapae* | 0.04701 | 0.02252 | 2.08763 | 0.03683 |
| temperature : *Plebejus argus* | 0.1363 | 0.02543 | 5.35916 | 0 |
| temperature : *Polygonia c-album* | 0.08545 | 0.04744 | 1.80129 | 0.07166 |
| temperature : *Polyommatus icarus* | -0.074 | 0.02279 | -3.24651 | 0.00117 |
| temperature : *Pyrgus malvae* | -0.06189 | 0.03776 | -1.6392 | 0.10117 |
| temperature : *Pyronia tithonus* | 0.00844 | 0.00783 | 1.07735 | 0.28133 |
| *Aglais urticae* : year | 0.00373 | 0.0065 | 0.57399 | 0.56597 |
| *Anthocharis cardamines* : year | -0.02082 | 0.01111 | -1.87351 | 0.061 |
| *Aphantopus hyperantus* : year | -0.03326 | 0.00821 | -4.0505 | 5e-05 |
| *Argynnis adippe* : year | -0.13421 | 0.08477 | -1.58322 | 0.11337 |
| *Argynnis aglaja* : year | -0.01526 | 0.01297 | -1.17618 | 0.23953 |
| *Argynnis paphia* : year | -0.00895 | 0.02961 | -0.30216 | 0.76253 |
| *Aricia agestis* : year | -0.05703 | 0.02045 | -2.78908 | 0.00529 |
| *Aricia artaxerxes* : year | 0.00025 | 0.01654 | 0.01518 | 0.98789 |
| *Callophrys rubi* : year | -0.12364 | 0.06684 | -1.84981 | 0.06434 |
| *Celastrina argiolus* : year | -0.09302 | 0.05142 | -1.80914 | 0.07043 |
| *Coenonympha pamphilus* : year | -0.02596 | 0.00999 | -2.59955 | 0.00934 |
| *Coenonympha tullia* : year | -0.02862 | 0.00874 | -3.27261 | 0.00107 |
| *Colias croceus* : year | -0.0183 | 0.01066 | -1.71649 | 0.08608 |
| *Cupido minimus* : year | -0.10518 | 0.04925 | -2.13575 | 0.0327 |
| *Erebia aethiops* : year | 0.00427 | 0.00952 | 0.4482 | 0.65401 |
| *Euphydryas aurinia* : year | -0.10281 | 0.02793 | -3.68106 | 0.00023 |
| *Hipparchia semele* : year | -0.06759 | 0.02769 | -2.44121 | 0.01464 |
| *Lasiommata megera* : year | -0.14433 | 0.04598 | -3.13916 | 0.00169 |
| *Leptidea sinapis* : year | -0.18911 | 0.0432 | -4.37752 | 1e-05 |
| *Lycaena phlaeas* : year | 0.00223 | 0.00586 | 0.38118 | 0.70307 |
| *Lysandra bellargus* : year | -0.0331 | 0.01968 | -1.68159 | 0.09265 |
| *Lysandra coridon* : year | -0.09567 | 0.01961 | -4.87838 | 0 |
| *Maculinea arion* : year | 0.03186 | 0.02643 | 1.2054 | 0.22805 |
| *Maniola jurtina* : year | -0.01927 | 0.00766 | -2.51572 | 0.01188 |
| *Melanargia galathea* : year | -0.02883 | 0.02522 | -1.14276 | 0.25314 |
| *Melitaea athalia* : year | 0.04933 | 0.05445 | 0.90604 | 0.36492 |
| *Pararge aegeria* : year | -0.00495 | 0.00997 | -0.496 | 0.6199 |
| *Pieris napi* : year | 0.08543 | 0.03702 | 2.3076 | 0.02102 |
| *Pieris rapae* : year | -0.04122 | 0.02112 | -1.95226 | 0.05091 |
| *Plebejus argus* : year | -0.17333 | 0.02473 | -7.00756 | 0 |
| *Polygonia c-album* : year | -0.02773 | 0.04621 | -0.60004 | 0.54848 |
| *Polyommatus icarus* : year | -0.1371 | 0.01953 | -7.02081 | 0 |
| *Pyrgus malvae* : year | -0.01767 | 0.02014 | -0.87743 | 0.38025 |
| *Pyronia tithonus* : year | -0.04047 | 0.00721 | -5.61566 | 0 |

***Supp. Table 11: The PC2 linear model results.***

| Term | Value | Std. Error | t value | P value |
|---|---|---|---|---|
| *Aglais urticae* | -0.36812 | 0.02416 | -15.23366 | 0 |
| *Anthocharis cardamines* | 0.09125 | 0.02237 | 4.07961 | 5e-05 |
| *Aphantopus hyperantus* | -0.21863 | 0.0193 | -11.32693 | 0 |
| *Argynnis adippe* | 0.46184 | 0.03116 | 14.81972 | 0 |
| *Argynnis aglaja* | 0.07378 | 0.02621 | 2.81534 | 0.00487 |
| *Argynnis paphia* | -0.0918 | 0.0281 | -3.26681 | 0.00109 |
| *Aricia agestis* | 0.16295 | 0.02131 | 7.64673 | 0 |
| *Aricia artaxerxes* | -0.02617 | 0.05145 | -0.50866 | 0.61099 |
| *Callophrys rubi* | -0.13986 | 0.02999 | -4.66304 | 0 |
| *Celastrina argiolus* | 0.18994 | 0.02768 | 6.86114 | 0 |
| *Coenonympha pamphilus* | -0.37355 | 0.01693 | -22.0586 | 0 |
| *Coenonympha tullia* | -0.34921 | 0.02178 | -16.03289 | 0 |
| *Colias croceus* | -0.31839 | 0.04612 | -6.90411 | 0 |
| *Cupido minimus* | 0.00763 | 0.02565 | 0.29756 | 0.76604 |
| *Erebia aethiops* | -0.28984 | 0.0535 | -5.41782 | 0 |
| *Euphydryas aurinia* | -0.04569 | 0.01791 | -2.55164 | 0.01072 |
| *Hipparchia semele* | 0.02335 | 0.02023 | 1.15381 | 0.24858 |
| *Lasiommata megera* | -0.10407 | 0.02787 | -3.73446 | 0.00019 |
| *Leptidea sinapis* | 1.37907 | 0.02745 | 50.23413 | 0 |
| *Lycaena phlaeas* | -0.10999 | 0.01585 | -6.93976 | 0 |
| *Lysandra bellargus* | 0.4249 | 0.0161 | 26.38701 | 0 |
| *Lysandra coridon* | -0.04005 | 0.00693 | -5.77799 | 0 |
| *Maculinea arion* | 0.04084 | 0.06078 | 0.67195 | 0.50162 |
| *Maniola jurtina* | -0.22838 | 0.01293 | -17.66708 | 0 |
| *Melanargia galathea* | -0.0498 | 0.02485 | -2.00375 | 0.0451 |
| *Melitaea athalia* | 0.27568 | 0.04827 | 5.71171 | 0 |
| *Pararge aegeria* | -0.43479 | 0.02371 | -18.34012 | 0 |
| *Pieris napi* | 0.29757 | 0.01822 | 16.33124 | 0 |
| *Pieris rapae* | 0.34153 | 0.02652 | 12.88003 | 0 |
| *Plebejus argus* | 0.11254 | 0.01076 | 10.46039 | 0 |
| *Polygonia c-album* | -0.35975 | 0.03074 | -11.70158 | 0 |
| *Polyommatus icarus* | 0.05596 | 0.00942 | 5.9379 | 0 |
| *Pyrgus malvae* | 0.01713 | 0.02927 | 0.58522 | 0.5584 |
| *Pyronia tithonus* | -0.35953 | 0.02178 | -16.50766 | 0 |
| temperature : *Aglais urticae* | 0.01931 | 0.02964 | 0.65122 | 0.51491 |
| temperature : *Anthocharis cardamines* | -0.09702 | 0.0289 | -3.35694 | 0.00079 |
| temperature : *Aphantopus hyperantus* | 0.03205 | 0.01739 | 1.84327 | 0.06529 |
| temperature : *Argynnis adippe* | 0.04907 | 0.04556 | 1.07704 | 0.28146 |
| temperature : *Argynnis aglaja* | 0.04578 | 0.01881 | 2.43354 | 0.01495 |
| temperature : *Argynnis paphia* | 0.0215 | 0.04707 | 0.45678 | 0.64783 |
| temperature : *Aricia agestis* | -0.03973 | 0.0303 | -1.31107 | 0.18984 |
| temperature : *Aricia artaxerxes* | 0.00344 | 0.02091 | 0.16471 | 0.86918 |
| temperature : *Callophrys rubi* | -0.00141 | 0.02632 | -0.05348 | 0.95735 |
| temperature : *Celastrina argiolus* | -0.0921 | 0.04409 | -2.08873 | 0.03673 |
| temperature : *Coenonympha pamphilus* | -0.05118 | 0.01629 | -3.14179 | 0.00168 |
| temperature : *Coenonympha tullia* | -0.00558 | 0.01154 | -0.48388 | 0.62847 |
| temperature : *Colias croceus* | -0.04256 | 0.04779 | -0.89059 | 0.37315 |
| temperature : *Cupido minimus* | -0.02597 | 0.03004 | -0.86463 | 0.38724 |
| temperature : *Erebia aethiops* | -0.00047 | 0.01902 | -0.02468 | 0.98031 |
| temperature : *Euphydryas aurinia* | -0.05729 | 0.01995 | -2.8718 | 0.00408 |
| temperature : *Hipparchia semele* | 0.09321 | 0.02112 | 4.41374 | 1e-05 |
| temperature : *Lasiommata megera* | 0.02872 | 0.03791 | 0.75772 | 0.44862 |
| temperature : *Leptidea sinapis* | 0.08577 | 0.04647 | 1.84571 | 0.06494 |
| temperature : *Lycaena phlaeas* | 0.0296 | 0.01922 | 1.53983 | 0.1236 |
| temperature : *Lysandra bellargus* | 0.04829 | 0.01927 | 2.50559 | 0.01223 |

| | | | | |
|---|---|---|---|---|
| temperature : *Lysandra coridon* | 0.01269 | 0.011 | 1.15381 | 0.24858 |
| temperature : *Maculinea arion* | -0.07546 | 0.08014 | -0.94167 | 0.34636 |
| temperature : *Maniola jurtina* | 0.00131 | 0.01513 | 0.08651 | 0.93106 |
| temperature : *Melanargia galathea* | 0.01657 | 0.03647 | 0.45447 | 0.64949 |
| temperature : *Melitaea athalia* | 0.03299 | 0.04295 | 0.76805 | 0.44246 |
| temperature : *Pararge aegeria* | -0.02023 | 0.0292 | -0.69301 | 0.48831 |
| temperature : *Pieris napi* | -0.05762 | 0.01395 | -4.12942 | 4e-05 |
| temperature : *Pieris rapae* | -0.01033 | 0.03089 | -0.33444 | 0.73805 |
| temperature : *Plebejus argus* | 0.02098 | 0.01448 | 1.44863 | 0.14744 |
| temperature : *Polygonia c-album* | 0.02694 | 0.04634 | 0.58135 | 0.56101 |
| temperature : *Polyommatus icarus* | 0.0271 | 0.00909 | 2.98183 | 0.00287 |
| temperature : *Pyrgus malvae* | 0.00157 | 0.04624 | 0.03385 | 0.973 |
| temperature : *Pyronia tithonus* | -0.00681 | 0.0337 | -0.20198 | 0.83993 |
| *Aglais urticae* : year | 0.02055 | 0.02017 | 1.0185 | 0.30844 |
| *Anthocharis cardamines* : year | 0.00305 | 0.01876 | 0.16245 | 0.87095 |
| *Aphantopus hyperantus* : year | 0.05531 | 0.0191 | 2.89615 | 0.00378 |
| *Argynnis adippe* : year | 0.0207 | 0.0318 | 0.65095 | 0.51508 |
| *Argynnis aglaja* : year | 0.03162 | 0.02214 | 1.42846 | 0.15316 |
| *Argynnis paphia* : year | 0.02539 | 0.02373 | 1.06989 | 0.28467 |
| *Aricia agestis* : year | 0.03461 | 0.0207 | 1.67212 | 0.0945 |
| *Aricia artaxerxes* : year | -0.00327 | 0.02598 | -0.126 | 0.89973 |
| *Callophrys rubi* : year | 0.04122 | 0.02968 | 1.38894 | 0.16485 |
| *Celastrina argiolus* : year | 0.01697 | 0.02489 | 0.68192 | 0.49529 |
| *Coenonympha pamphilus* : year | -0.01436 | 0.0154 | -0.93237 | 0.35115 |
| *Coenonympha tullia* : year | 0.0791 | 0.01528 | 5.17757 | 0 |
| *Colias croceus* : year | 0.01987 | 0.03619 | 0.54893 | 0.58305 |
| *Cupido minimus* : year | -0.00193 | 0.02608 | -0.07411 | 0.94092 |
| *Erebia aethiops* : year | 0.04382 | 0.02055 | 2.13273 | 0.03295 |
| *Euphydryas aurinia* : year | 0.0956 | 0.01637 | 5.83963 | 0 |
| *Hipparchia semele* : year | 0.05067 | 0.01687 | 3.00327 | 0.00267 |
| *Lasiommata megera* : year | -0.02109 | 0.02585 | -0.8159 | 0.41456 |
| *Leptidea sinapis* : year | 0.03973 | 0.02772 | 1.43312 | 0.15183 |
| *Lycaena phlaeas* : year | 0.09064 | 0.01548 | 5.85569 | 0 |
| *Lysandra bellargus* : year | 0.02426 | 0.01324 | 1.83309 | 0.06679 |
| *Lysandra coridon* : year | 0.00252 | 0.0075 | 0.33594 | 0.73692 |
| *Maculinea arion* : year | 0.10312 | 0.03245 | 3.17802 | 0.00148 |
| *Maniola jurtina* : year | 0.04265 | 0.01258 | 3.39145 | 7e-04 |
| *Melanargia galathea* : year | -0.09292 | 0.01805 | -5.14872 | 0 |
| *Melitaea athalia* : year | 0.08977 | 0.0312 | 2.8777 | 0.00401 |
| *Pararge aegeria* : year | 0.00267 | 0.02189 | 0.12211 | 0.90281 |
| *Pieris napi* : year | -0.01087 | 0.01558 | -0.6975 | 0.48549 |
| *Pieris rapae* : year | 0.02376 | 0.02505 | 0.94854 | 0.34286 |
| *Plebejus argus* : year | 0.06051 | 0.01239 | 4.8824 | 0 |
| *Polygonia c-album* : year | 0.01963 | 0.03418 | 0.57435 | 0.56573 |
| *Polyommatus icarus* : year | 0.03749 | 0.01033 | 3.62861 | 0.00029 |
| *Pyrgus malvae* : year | -0.00649 | 0.02694 | -0.24106 | 0.80951 |
| *Pyronia tithonus* : year | 0.02218 | 0.01768 | 1.25437 | 0.20971 |

*Supp. Table 12: The PC2 quantile regression model results for a τ value of 0.1.*

| Term | Value | Std. Error | t value | P value |
|---|---|---|---|---|
| *Aglais urticae* | -0.95656 | 0.01171 | -81.68611 | 0 |
| *Anthocharis cardamines* | -0.95219 | 0.011 | -86.58719 | 0 |
| *Aphantopus hyperantus* | -0.98787 | 0.00718 | -137.63612 | 0 |
| *Argynnis adippe* | -0.87087 | 0.01615 | -53.91501 | 0 |
| *Argynnis aglaja* | -0.92454 | 0.01258 | -73.48092 | 0 |
| *Argynnis paphia* | -0.95005 | 0.01091 | -87.08759 | 0 |
| *Aricia agestis* | -0.89673 | 0.01621 | -55.32556 | 0 |
| *Aricia artaxerxes* | -0.89343 | 0.02424 | -36.86372 | 0 |
| *Callophrys rubi* | -0.9821 | 0.01703 | -57.67246 | 0 |
| *Celastrina argiolus* | -0.90777 | 0.02006 | -45.24301 | 0 |
| *Coenonympha pamphilus* | -1.01352 | 0.00668 | -151.63898 | 0 |
| *Coenonympha tullia* | -1.00339 | 0.00692 | -144.91019 | 0 |
| *Colias croceus* | -0.96505 | 0.02923 | -33.01051 | 0 |
| *Cupido minimus* | -0.92363 | 0.01042 | -88.6033 | 0 |
| *Erebia aethiops* | -0.94669 | 0.02616 | -36.1818 | 0 |
| *Euphydryas aurinia* | -0.94902 | 0.00869 | -109.14742 | 0 |
| *Hipparchia semele* | -0.96845 | 0.00813 | -119.11561 | 0 |
| *Lasiommata megera* | -0.94411 | 0.01052 | -89.72381 | 0 |
| *Leptidea sinapis* | -0.75234 | 0.0188 | -40.01134 | 0 |
| *Lycaena phlaeas* | -0.9374 | 0.00702 | -133.58155 | 0 |
| *Lysandra bellargus* | -0.87528 | 0.0092 | -95.13872 | 0 |
| *Lysandra coridon* | -0.96058 | 0.00298 | -322.03237 | 0 |
| *Maculinea arion* | -0.97587 | 0.01424 | -68.52719 | 0 |
| *Maniola jurtina* | -0.9664 | 0.00565 | -170.91347 | 0 |
| *Melanargia galathea* | -0.96457 | 0.00594 | -162.40038 | 0 |
| *Melitaea athalia* | -0.92446 | 0.02175 | -42.50534 | 0 |
| *Pararge aegeria* | -0.98933 | 0.00765 | -129.34851 | 0 |
| *Pieris napi* | -0.92851 | 0.00988 | -93.93559 | 0 |
| *Pieris rapae* | -0.9198 | 0.00877 | -104.83179 | 0 |
| *Plebejus argus* | -0.94305 | 0.00537 | -175.65025 | 0 |
| *Polygonia c-album* | -0.98289 | 0.01067 | -92.14255 | 0 |
| *Polyommatus icarus* | -0.9393 | 0.00452 | -207.69144 | 0 |
| *Pyrgus malvae* | -0.93298 | 0.00947 | -98.5367 | 0 |
| *Pyronia tithonus* | -1.01495 | 0.00589 | -172.3817 | 0 |
| temperature : *Aglais urticae* | 0.00562 | 0.01532 | 0.36664 | 0.71389 |
| temperature : *Anthocharis cardamines* | -0.0182 | 0.01382 | -1.31687 | 0.18789 |
| temperature : *Aphantopus hyperantus* | -0.002 | 0.00725 | -0.27609 | 0.78248 |
| temperature : *Argynnis adippe* | 0.00079 | 0.02347 | 0.03346 | 0.97331 |
| temperature : *Argynnis aglaja* | 0.01505 | 0.00318 | 4.73203 | 0 |
| temperature : *Argynnis paphia* | 0.02244 | 0.01826 | 1.22928 | 0.21897 |
| temperature : *Aricia agestis* | 0.02472 | 0.01553 | 1.59226 | 0.11133 |
| temperature : *Aricia artaxerxes* | 0.00719 | 0.0089 | 0.80845 | 0.41883 |
| temperature : *Callophrys rubi* | 0.00851 | 0.02793 | 0.30486 | 0.76048 |
| temperature : *Celastrina argiolus* | -0.0277 | 0.02935 | -0.94376 | 0.34529 |
| temperature : *Coenonympha pamphilus* | -0.01319 | 0.00518 | -2.54736 | 0.01086 |
| temperature : *Coenonympha tullia* | 0.00267 | 0.00426 | 0.62553 | 0.53162 |
| temperature : *Colias croceus* | -0.02326 | 0.0262 | -0.888 | 0.37454 |
| temperature : *Cupido minimus* | -0.00954 | 0.01117 | -0.85351 | 0.39338 |
| temperature : *Erebia aethiops* | 0.00414 | 0.00961 | 0.43111 | 0.66639 |
| temperature : *Euphydryas aurinia* | 0.00146 | 0.01052 | 0.13835 | 0.88997 |
| temperature : *Hipparchia semele* | 0.0066 | 0.00974 | 0.67824 | 0.49762 |
| temperature : *Lasiommata megera* | -0.02207 | 0.01249 | -1.76724 | 0.07719 |
| temperature : *Leptidea sinapis* | 0.11465 | 0.03056 | 3.75165 | 0.00018 |
| temperature : *Lycaena phlaeas* | 0.00969 | 0.00776 | 1.24773 | 0.21213 |
| temperature : *Lysandra bellargus* | -0.01472 | 0.01155 | -1.27511 | 0.20227 |

| | | | | |
|---|---|---|---|---|
| temperature : *Lysandra coridon* | 0.00581 | 0.00471 | 1.23332 | 0.21746 |
| temperature : *Maculinea arion* | 0.01028 | 0.01991 | 0.51615 | 0.60575 |
| temperature : *Maniola jurtina* | -0.00156 | 0.0073 | -0.21296 | 0.83136 |
| temperature : *Melanargia galathea* | -0.00577 | 0.01034 | -0.55837 | 0.57659 |
| temperature : *Melitaea athalia* | 0.02306 | 0.02355 | 0.9792 | 0.32748 |
| temperature : *Pararge aegeria* | -0.00511 | 0.00701 | -0.72944 | 0.46574 |
| temperature : *Pieris napi* | 0.00078 | 0.00737 | 0.10653 | 0.91517 |
| temperature : *Pieris rapae* | -0.00293 | 0.00586 | -0.50023 | 0.61691 |
| temperature : *Plebejus argus* | 0.00467 | 0.00686 | 0.6804 | 0.49625 |
| temperature : *Polygonia c-album* | 0.0225 | 0.01132 | 1.98768 | 0.04685 |
| temperature : *Polyommatus icarus* | 0.00344 | 0.00341 | 1.00826 | 0.31333 |
| temperature : *Pyrgus malvae* | -0.00414 | 0.01653 | -0.25034 | 0.80232 |
| temperature : *Pyronia tithonus* | -0.00725 | 0.00905 | -0.80108 | 0.42309 |
| *Aglais urticae* : year | -0.00288 | 0.00884 | -0.32587 | 0.74452 |
| *Anthocharis cardamines* : year | -0.00186 | 0.00905 | -0.20599 | 0.8368 |
| *Aphantopus hyperantus* : year | 0.01546 | 0.00608 | 2.54321 | 0.01099 |
| *Argynnis adippe* : year | 0.00056 | 0.01678 | 0.0335 | 0.97328 |
| *Argynnis aglaja* : year | -0.02995 | 0.00625 | -4.79134 | 0 |
| *Argynnis paphia* : year | -0.01532 | 0.00669 | -2.28985 | 0.02203 |
| *Aricia agestis* : year | -0.02462 | 0.00904 | -2.72238 | 0.00648 |
| *Aricia artaxerxes* : year | -0.03175 | 0.01062 | -2.99044 | 0.00279 |
| *Callophrys rubi* : year | -0.00245 | 0.01543 | -0.1588 | 0.87382 |
| *Celastrina argiolus* : year | 0.01995 | 0.01817 | 1.09796 | 0.27223 |
| *Coenonympha pamphilus* : year | 0.00166 | 0.00559 | 0.2966 | 0.76678 |
| *Coenonympha tullia* : year | 0.00897 | 0.00546 | 1.64407 | 0.10016 |
| *Colias croceus* : year | 0.00908 | 0.0199 | 0.45634 | 0.64814 |
| *Cupido minimus* : year | -0.0078 | 0.00969 | -0.80484 | 0.42092 |
| *Erebia aethiops* : year | -0.00557 | 0.00984 | -0.56601 | 0.57139 |
| *Euphydryas aurinia* : year | 0.00604 | 0.0082 | 0.73706 | 0.46109 |
| *Hipparchia semele* : year | -0.01114 | 0.00663 | -1.67969 | 0.09302 |
| *Lasiommata megera* : year | -0.01236 | 0.00612 | -2.01866 | 0.04353 |
| *Leptidea sinapis* : year | 0.01106 | 0.01897 | 0.58307 | 0.55985 |
| *Lycaena phlaeas* : year | 0.00307 | 0.00698 | 0.44034 | 0.65969 |
| *Lysandra bellargus* : year | 0.00366 | 0.0078 | 0.46963 | 0.63862 |
| *Lysandra coridon* : year | -0.00396 | 0.00326 | -1.21453 | 0.22455 |
| *Maculinea arion* : year | -0.00071 | 0.00804 | -0.0888 | 0.92924 |
| *Maniola jurtina* : year | -0.01106 | 0.00554 | -1.99502 | 0.04604 |
| *Melanargia galathea* : year | -0.01287 | 0.00494 | -2.60687 | 0.00914 |
| *Melitaea athalia* : year | -0.00132 | 0.01059 | -0.12482 | 0.90066 |
| *Pararge aegeria* : year | -0.01 | 0.00288 | -3.47192 | 0.00052 |
| *Pieris napi* : year | -0.00348 | 0.0051 | -0.68254 | 0.4949 |
| *Pieris rapae* : year | -0.01902 | 0.00507 | -3.75442 | 0.00017 |
| *Plebejus argus* : year | 0.00786 | 0.00633 | 1.24263 | 0.21401 |
| *Polygonia c-album* : year | -0.02685 | 0.0103 | -2.60785 | 0.00911 |
| *Polyommatus icarus* : year | 0.00055 | 0.00449 | 0.12224 | 0.90271 |
| *Pyrgus malvae* : year | 0.00806 | 0.00926 | 0.87099 | 0.38376 |
| *Pyronia tithonus* : year | 3e-05 | 0.0044 | 0.00713 | 0.99431 |

***Supp. Table 13: The PC2 quantile regression model results for a τ value of 0.25.***

| Term | Value | Std. Error | t value | P value |
|---|---|---|---|---|
| *Aglais urticae* | -0.73141 | 0.0152 | -48.12262 | 0 |
| *Anthocharis cardamines* | -0.71947 | 0.01381 | -52.09771 | 0 |
| *Aphantopus hyperantus* | -0.80724 | 0.01268 | -63.67176 | 0 |
| *Argynnis adippe* | -0.5334 | 0.02851 | -18.70821 | 0 |
| *Argynnis aglaja* | -0.6513 | 0.02008 | -32.4408 | 0 |
| *Argynnis paphia* | -0.73749 | 0.02051 | -35.96542 | 0 |
| *Aricia agestis* | -0.61227 | 0.01709 | -35.82762 | 0 |
| *Aricia artaxerxes* | -0.66418 | 0.04449 | -14.93002 | 0 |
| *Callophrys rubi* | -0.74567 | 0.01666 | -44.76376 | 0 |
| *Celastrina argiolus* | -0.64631 | 0.03946 | -16.37847 | 0 |
| *Coenonympha pamphilus* | -0.84876 | 0.00727 | -116.78883 | 0 |
| *Coenonympha tullia* | -0.86109 | 0.01028 | -83.76646 | 0 |
| *Colias croceus* | -0.77342 | 0.02406 | -32.14756 | 0 |
| *Cupido minimus* | -0.6948 | 0.01589 | -43.73846 | 0 |
| *Erebia aethiops* | -0.78227 | 0.03144 | -24.87851 | 0 |
| *Euphydryas aurinia* | -0.74404 | 0.01101 | -67.57271 | 0 |
| *Hipparchia semele* | -0.768 | 0.01309 | -58.67131 | 0 |
| *Lasiommata megera* | -0.72244 | 0.01447 | -49.94229 | 0 |
| *Leptidea sinapis* | -0.17604 | 0.05034 | -3.49674 | 0.00047 |
| *Lycaena phlaeas* | -0.70229 | 0.01045 | -67.19229 | 0 |
| *Lysandra bellargus* | -0.56401 | 0.01812 | -31.1259 | 0 |
| *Lysandra coridon* | -0.73505 | 0.00494 | -148.65711 | 0 |
| *Maculinea arion* | -0.76781 | 0.02109 | -36.39783 | 0 |
| *Maniola jurtina* | -0.78053 | 0.0085 | -91.79086 | 0 |
| *Melanargia galathea* | -0.74713 | 0.01504 | -49.66653 | 0 |
| *Melitaea athalia* | -0.68054 | 0.04794 | -14.1955 | 0 |
| *Pararge aegeria* | -0.85301 | 0.00927 | -92.03204 | 0 |
| *Pieris napi* | -0.65743 | 0.01625 | -40.46667 | 0 |
| *Pieris rapae* | -0.6504 | 0.0248 | -26.22197 | 0 |
| *Plebejus argus* | -0.69301 | 0.00764 | -90.72224 | 0 |
| *Polygonia c-album* | -0.81892 | 0.02008 | -40.79142 | 0 |
| *Polyommatus icarus* | -0.6901 | 0.00722 | -95.62867 | 0 |
| *Pyrgus malvae* | -0.68791 | 0.01957 | -35.15809 | 0 |
| *Pyronia tithonus* | -0.87304 | 0.01129 | -77.30094 | 0 |
| temperature : *Aglais urticae* | 0.03825 | 0.01775 | 2.15471 | 0.03119 |
| temperature : *Anthocharis cardamines* | -0.06506 | 0.01582 | -4.11329 | 4e-05 |
| temperature : *Aphantopus hyperantus* | 0.0118 | 0.01084 | 1.08794 | 0.27663 |
| temperature : *Argynnis adippe* | -0.01727 | 0.03771 | -0.45792 | 0.64701 |
| temperature : *Argynnis aglaja* | 0.02507 | 0.01684 | 1.48864 | 0.13659 |
| temperature : *Argynnis paphia* | 0.03624 | 0.033 | 1.09846 | 0.272 |
| temperature : *Aricia agestis* | 0.00195 | 0.02384 | 0.08191 | 0.93472 |
| temperature : *Aricia artaxerxes* | -0.00152 | 0.02434 | -0.06257 | 0.95011 |
| temperature : *Callophrys rubi* | 0.00833 | 0.01294 | 0.6434 | 0.51997 |
| temperature : *Celastrina argiolus* | -0.05071 | 0.03612 | -1.40399 | 0.16032 |
| temperature : *Coenonympha pamphilus* | -0.01864 | 0.00703 | -2.65326 | 0.00797 |
| temperature : *Coenonympha tullia* | 0.00313 | 0.00456 | 0.687 | 0.49208 |
| temperature : *Colias croceus* | -0.03415 | 0.0251 | -1.3607 | 0.17361 |
| temperature : *Cupido minimus* | -0.01099 | 0.01283 | -0.85654 | 0.3917 |
| temperature : *Erebia aethiops* | -0.00527 | 0.0103 | -0.51159 | 0.60894 |
| temperature : *Euphydryas aurinia* | -0.02028 | 0.01364 | -1.48625 | 0.13722 |
| temperature : *Hipparchia semele* | 0.02078 | 0.00647 | 3.21331 | 0.00131 |
| temperature : *Lasiommata megera* | -0.02478 | 0.02162 | -1.14611 | 0.25175 |
| temperature : *Leptidea sinapis* | 0.14432 | 0.08501 | 1.69769 | 0.08957 |
| temperature : *Lycaena phlaeas* | 0.0268 | 0.01157 | 2.3159 | 0.02057 |
| temperature : *Lysandra bellargus* | 0.03148 | 0.02051 | 1.53517 | 0.12475 |

| | | | | |
|---|---|---|---|---|
| temperature : *Lysandra coridon* | 0.00692 | 0.0076 | 0.91114 | 0.36222 |
| temperature : *Maculinea arion* | -0.00526 | 0.03391 | -0.15509 | 0.87675 |
| temperature : *Maniola jurtina* | 0.00753 | 0.0097 | 0.7764 | 0.43752 |
| temperature : *Melanargia galathea* | -0.03996 | 0.02289 | -1.74566 | 0.08087 |
| temperature : *Melitaea athalia* | 0.06492 | 0.03072 | 2.11339 | 0.03457 |
| temperature : *Pararge aegeria* | 0.00429 | 0.01095 | 0.39206 | 0.69502 |
| temperature : *Pieris napi* | -0.02576 | 0.01468 | -1.75436 | 0.07937 |
| temperature : *Pieris rapae* | -0.01768 | 0.03085 | -0.57289 | 0.56672 |
| temperature : *Plebejus argus* | 0.01348 | 0.01023 | 1.31759 | 0.18764 |
| temperature : *Polygonia c-album* | 0.02957 | 0.02544 | 1.16253 | 0.24502 |
| temperature : *Polyommatus icarus* | 0.00482 | 0.00597 | 0.80788 | 0.41916 |
| temperature : *Pyrgus malvae* | 2e-04 | 0.02735 | 0.00717 | 0.99428 |
| temperature : *Pyronia tithonus* | 0.01463 | 0.01948 | 0.75085 | 0.45275 |
| *Aglais urticae* : year | -0.0072 | 0.01219 | -0.5905 | 0.55485 |
| *Anthocharis cardamines* : year | -0.0054 | 0.01139 | -0.47424 | 0.63533 |
| *Aphantopus hyperantus* : year | 0.02127 | 0.01163 | 1.82974 | 0.06729 |
| *Argynnis adippe* : year | -0.01642 | 0.02598 | -0.63193 | 0.52743 |
| *Argynnis aglaja* : year | -0.0028 | 0.01824 | -0.15375 | 0.87781 |
| *Argynnis paphia* : year | -0.0179 | 0.01908 | -0.93832 | 0.34808 |
| *Aricia agestis* : year | -0.02559 | 0.01602 | -1.59812 | 0.11002 |
| *Aricia artaxerxes* : year | -0.05757 | 0.02453 | -2.34681 | 0.01894 |
| *Callophrys rubi* : year | 0.0214 | 0.01668 | 1.2828 | 0.19956 |
| *Celastrina argiolus* : year | -0.00172 | 0.02703 | -0.06363 | 0.94926 |
| *Coenonympha pamphilus* : year | -0.00521 | 0.00621 | -0.83815 | 0.40195 |
| *Coenonympha tullia* : year | 0.01498 | 0.00726 | 2.06292 | 0.03912 |
| *Colias croceus* : year | 0.01901 | 0.01933 | 0.98377 | 0.32523 |
| *Cupido minimus* : year | -0.0188 | 0.01595 | -1.17869 | 0.23852 |
| *Erebia aethiops* : year | -0.0123 | 0.01172 | -1.04971 | 0.29385 |
| *Euphydryas aurinia* : year | 0.03666 | 0.01069 | 3.43079 | 6e-04 |
| *Hipparchia semele* : year | -0.00263 | 0.01019 | -0.25799 | 0.79642 |
| *Lasiommata megera* : year | -0.03982 | 0.01431 | -2.78188 | 0.00541 |
| *Leptidea sinapis* : year | 0.08362 | 0.0528 | 1.58374 | 0.11326 |
| *Lycaena phlaeas* : year | 6e-05 | 0.00911 | 0.00659 | 0.99474 |
| *Lysandra bellargus* : year | -0.00355 | 0.01502 | -0.23655 | 0.81301 |
| *Lysandra coridon* : year | 0.00097 | 0.00517 | 0.18717 | 0.85153 |
| *Maculinea arion* : year | 0.02096 | 0.01654 | 1.26756 | 0.20496 |
| *Maniola jurtina* : year | -0.01057 | 0.00815 | -1.29661 | 0.19477 |
| *Melanargia galathea* : year | -0.03237 | 0.01118 | -2.89651 | 0.00377 |
| *Melitaea athalia* : year | 0.04331 | 0.02488 | 1.74075 | 0.08173 |
| *Pararge aegeria* : year | -0.00921 | 0.00747 | -1.23345 | 0.21741 |
| *Pieris napi* : year | -0.01875 | 0.01498 | -1.25132 | 0.21082 |
| *Pieris rapae* : year | -0.01264 | 0.02371 | -0.53314 | 0.59394 |
| *Plebejus argus* : year | 0.01022 | 0.00889 | 1.15001 | 0.25014 |
| *Polygonia c-album* : year | -0.00372 | 0.02157 | -0.17248 | 0.86306 |
| *Polyommatus icarus* : year | 0.01134 | 0.00782 | 1.45005 | 0.14705 |
| *Pyrgus malvae* : year | 0.01606 | 0.01611 | 0.99701 | 0.31876 |
| *Pyronia tithonus* : year | 0.00242 | 0.01001 | 0.24179 | 0.80894 |

***Supp. Table 14: The PC2 quantile regression model results for a τ value of 0.5.***

| Term | Value | Std. Error | t value | P value |
|---|---|---|---|---|
| *Aglais urticae* | -0.47333 | 0.01081 | -43.77645 | 0 |
| *Anthocharis cardamines* | -0.27332 | 0.02456 | -11.12678 | 0 |
| *Aphantopus hyperantus* | -0.45155 | 0.01767 | -25.54825 | 0 |
| *Argynnis adippe* | 0.12803 | 0.04814 | 2.65945 | 0.00783 |
| *Argynnis aglaja* | -0.21047 | 0.02961 | -7.10695 | 0 |
| *Argynnis paphia* | -0.34841 | 0.0223 | -15.6266 | 0 |
| *Aricia agestis* | -0.1983 | 0.02069 | -9.58476 | 0 |
| *Aricia artaxerxes* | -0.28257 | 0.03223 | -8.76765 | 0 |
| *Callophrys rubi* | -0.2981 | 0.02041 | -14.60502 | 0 |
| *Celastrina argiolus* | -0.08153 | 0.02738 | -2.97725 | 0.00291 |
| *Coenonympha pamphilus* | -0.55848 | 0.01206 | -46.29742 | 0 |
| *Coenonympha tullia* | -0.55806 | 0.01514 | -36.86772 | 0 |
| *Colias croceus* | -0.45582 | 0.0325 | -14.02569 | 0 |
| *Cupido minimus* | -0.28272 | 0.02572 | -10.99305 | 0 |
| *Erebia aethiops* | -0.46275 | 0.03621 | -12.77885 | 0 |
| *Euphydryas aurinia* | -0.32644 | 0.02252 | -14.49355 | 0 |
| *Hipparchia semele* | -0.3828 | 0.01784 | -21.46256 | 0 |
| *Lasiommata megera* | -0.35809 | 0.02959 | -12.10062 | 0 |
| *Leptidea sinapis* | 1.5355 | 0.10046 | 15.28446 | 0 |
| *Lycaena phlaeas* | -0.37596 | 0.01235 | -30.44106 | 0 |
| *Lysandra bellargus* | 0.07894 | 0.02523 | 3.12928 | 0.00175 |
| *Lysandra coridon* | -0.31075 | 0.00714 | -43.50862 | 0 |
| *Maculinea arion* | -0.378 | 0.08737 | -4.32668 | 2e-05 |
| *Maniola jurtina* | -0.43837 | 0.01127 | -38.88935 | 0 |
| *Melanargia galathea* | -0.35145 | 0.02627 | -13.37756 | 0 |
| *Melitaea athalia* | -0.02575 | 0.05573 | -0.46211 | 0.644 |
| *Pararge aegeria* | -0.59912 | 0.01453 | -41.2322 | 0 |
| *Pieris napi* | -0.07723 | 0.02741 | -2.81697 | 0.00485 |
| *Pieris rapae* | -0.09363 | 0.05885 | -1.59094 | 0.11163 |
| *Plebejus argus* | -0.20908 | 0.01291 | -16.1897 | 0 |
| *Polygonia c-album* | -0.5406 | 0.01428 | -37.85693 | 0 |
| *Polyommatus icarus* | -0.19928 | 0.01127 | -17.68168 | 0 |
| *Pyrgus malvae* | -0.25248 | 0.04046 | -6.24036 | 0 |
| *Pyronia tithonus* | -0.5613 | 0.01464 | -38.34169 | 0 |
| temperature : *Aglais urticae* | 0.03143 | 0.01423 | 2.20927 | 0.02716 |
| temperature : *Anthocharis cardamines* | -0.11615 | 0.03264 | -3.55808 | 0.00037 |
| temperature : *Aphantopus hyperantus* | 0.02953 | 0.0129 | 2.28963 | 0.02204 |
| temperature : *Argynnis adippe* | 0.13628 | 0.05845 | 2.33167 | 0.01972 |
| temperature : *Argynnis aglaja* | 0.03234 | 0.01476 | 2.19118 | 0.02844 |
| temperature : *Argynnis paphia* | 0.05068 | 0.04633 | 1.09394 | 0.27398 |
| temperature : *Aricia agestis* | -0.06674 | 0.02708 | -2.46482 | 0.01371 |
| temperature : *Aricia artaxerxes* | 0.0096 | 0.01156 | 0.83028 | 0.40638 |
| temperature : *Callophrys rubi* | -0.01238 | 0.02726 | -0.45426 | 0.64964 |
| temperature : *Celastrina argiolus* | -0.114 | 0.04406 | -2.58718 | 0.00968 |
| temperature : *Coenonympha pamphilus* | -0.03358 | 0.01088 | -3.08551 | 0.00203 |
| temperature : *Coenonympha tullia* | 0.00654 | 0.00745 | 0.87782 | 0.38004 |
| temperature : *Colias croceus* | -0.08307 | 0.03021 | -2.74987 | 0.00596 |
| temperature : *Cupido minimus* | -0.01979 | 0.01829 | -1.08202 | 0.27925 |
| temperature : *Erebia aethiops* | 0.00918 | 0.01434 | 0.63977 | 0.52233 |
| temperature : *Euphydryas aurinia* | -0.03395 | 0.02507 | -1.35412 | 0.1757 |
| temperature : *Hipparchia semele* | 0.04505 | 0.01753 | 2.56999 | 0.01017 |
| temperature : *Lasiommata megera* | 0.01636 | 0.03604 | 0.45387 | 0.64993 |
| temperature : *Leptidea sinapis* | 0.14179 | 0.159 | 0.89176 | 0.37252 |
| temperature : *Lycaena phlaeas* | 0.01455 | 0.01491 | 0.97608 | 0.32903 |
| temperature : *Lysandra bellargus* | 0.06603 | 0.03025 | 2.18289 | 0.02905 |

| | | | | |
|---|---|---|---|---|
| temperature : *Lysandra coridon* | 0.01575 | 0.01127 | 1.3977 | 0.16221 |
| temperature : *Maculinea arion* | 0.01355 | 0.09437 | 0.14362 | 0.8858 |
| temperature : *Maniola jurtina* | 0.0105 | 0.01328 | 0.79102 | 0.42894 |
| temperature : *Melanargia galathea* | -0.03135 | 0.0372 | -0.84262 | 0.39944 |
| temperature : *Melitaea athalia* | -0.04262 | 0.05214 | -0.81739 | 0.4137 |
| temperature : *Pararge aegeria* | -0.02599 | 0.01743 | -1.49099 | 0.13597 |
| temperature : *Pieris napi* | -0.08809 | 0.02301 | -3.82913 | 0.00013 |
| temperature : *Pieris rapae* | -0.01862 | 0.0677 | -0.27501 | 0.78331 |
| temperature : *Plebejus argus* | 0.0018 | 0.01757 | 0.10242 | 0.91842 |
| temperature : *Polygonia c-album* | 0.02602 | 0.01304 | 1.995 | 0.04605 |
| temperature : *Polyommatus icarus* | 0.01668 | 0.01016 | 1.64252 | 0.10049 |
| temperature : *Pyrgus malvae* | 0.03216 | 0.05377 | 0.59814 | 0.54975 |
| temperature : *Pyronia tithonus* | -0.00325 | 0.02162 | -0.15025 | 0.88056 |
| *Aglais urticae* : year | -0.00186 | 0.00844 | -0.21997 | 0.8259 |
| *Anthocharis cardamines* : year | -0.01697 | 0.02066 | -0.82138 | 0.41143 |
| *Aphantopus hyperantus* : year | 0.04535 | 0.01709 | 2.65358 | 0.00797 |
| *Argynnis adippe* : year | -0.02223 | 0.04226 | -0.52607 | 0.59884 |
| *Argynnis aglaja* : year | 0.03188 | 0.02509 | 1.27054 | 0.20389 |
| *Argynnis paphia* : year | 0.00489 | 0.0189 | 0.25889 | 0.79572 |
| *Aricia agestis* : year | 0.01565 | 0.01889 | 0.82847 | 0.40741 |
| *Aricia artaxerxes* : year | -0.01838 | 0.01709 | -1.07551 | 0.28215 |
| *Callophrys rubi* : year | 0.01262 | 0.0135 | 0.93443 | 0.35008 |
| *Celastrina argiolus* : year | 0.03613 | 0.02593 | 1.39342 | 0.1635 |
| *Coenonympha pamphilus* : year | -0.01503 | 0.01046 | -1.43705 | 0.15071 |
| *Coenonympha tullia* : year | 0.0547 | 0.01087 | 5.03202 | 0 |
| *Colias croceus* : year | 0.0223 | 0.02335 | 0.95481 | 0.33968 |
| *Cupido minimus* : year | 0.00877 | 0.0258 | 0.34006 | 0.73381 |
| *Erebia aethiops* : year | -0.00293 | 0.01454 | -0.20164 | 0.8402 |
| *Euphydryas aurinia* : year | 0.07145 | 0.02057 | 3.47435 | 0.00051 |
| *Hipparchia semele* : year | 0.0339 | 0.01575 | 2.15213 | 0.03139 |
| *Lasiommata megera* : year | -0.02244 | 0.02172 | -1.03309 | 0.30157 |
| *Leptidea sinapis* : year | 0.13319 | 0.1015 | 1.31229 | 0.18942 |
| *Lycaena phlaeas* : year | 0.04303 | 0.01232 | 3.49142 | 0.00048 |
| *Lysandra bellargus* : year | 0.00372 | 0.02076 | 0.17924 | 0.85775 |
| *Lysandra coridon* : year | -0.0046 | 0.00738 | -0.62293 | 0.53333 |
| *Maculinea arion* : year | 0.05258 | 0.04448 | 1.18219 | 0.23713 |
| *Maniola jurtina* : year | 0.00553 | 0.01096 | 0.50447 | 0.61393 |
| *Melanargia galathea* : year | -0.08657 | 0.01843 | -4.69747 | 0 |
| *Melitaea athalia* : year | 0.09819 | 0.03663 | 2.68064 | 0.00735 |
| *Pararge aegeria* : year | -0.01257 | 0.01321 | -0.95215 | 0.34102 |
| *Pieris napi* : year | 0.00231 | 0.02565 | 0.09022 | 0.92811 |
| *Pieris rapae* : year | 0.05199 | 0.05785 | 0.89884 | 0.36874 |
| *Plebejus argus* : year | 0.05062 | 0.0149 | 3.39638 | 0.00068 |
| *Polygonia c-album* : year | -0.00598 | 0.01483 | -0.40344 | 0.68662 |
| *Polyommatus icarus* : year | 0.03179 | 0.01185 | 2.68144 | 0.00733 |
| *Pyrgus malvae* : year | 0.00504 | 0.0335 | 0.15029 | 0.88054 |
| *Pyronia tithonus* : year | 0.00766 | 0.01159 | 0.66134 | 0.50839 |

***Supp. Table 15: The PC2 quantile regression model results for a τ value of 0.75.***

| **Term** | **Value** | **Std. Error** | **t value** | **P value** |
|---|---|---|---|---|
| *Aglais urticae* | -0.18535 | 0.01682 | -11.02066 | 0 |
| *Anthocharis cardamines* | 0.73293 | 0.05333 | 13.74257 | 0 |
| *Aphantopus hyperantus* | 0.13236 | 0.02825 | 4.68461 | 0 |
| *Argynnis adippe* | 1.25045 | 0.09725 | 12.8587 | 0 |
| *Argynnis aglaja* | 0.62599 | 0.06795 | 9.2129 | 0 |
| *Argynnis paphia* | 0.29166 | 0.03987 | 7.31568 | 0 |
| *Aricia agestis* | 0.62024 | 0.05369 | 11.55123 | 0 |
| *Aricia artaxerxes* | 0.36113 | 0.08319 | 4.34119 | 1e-05 |
| *Callophrys rubi* | 0.2477 | 0.03834 | 6.46022 | 0 |
| *Celastrina argiolus* | 0.81286 | 0.04856 | 16.73861 | 0 |
| *Coenonympha pamphilus* | -0.12588 | 0.01805 | -6.97517 | 0 |
| *Coenonympha tullia* | -0.05885 | 0.02736 | -2.15073 | 0.0315 |
| *Colias croceus* | -0.06324 | 0.05251 | -1.20421 | 0.22851 |
| *Cupido minimus* | 0.39621 | 0.05226 | 7.58217 | 0 |
| *Erebia aethiops* | -0.10765 | 0.05271 | -2.04225 | 0.04113 |
| *Euphydryas aurinia* | 0.36634 | 0.03348 | 10.94207 | 0 |
| *Hipparchia semele* | 0.40804 | 0.04666 | 8.74416 | 0 |
| *Lasiommata megera* | 0.27716 | 0.04668 | 5.93748 | 0 |
| *Leptidea sinapis* | 2.87118 | 0.04371 | 65.6888 | 0 |
| *Lycaena phlaeas* | 0.21344 | 0.02501 | 8.53484 | 0 |
| *Lysandra bellargus* | 1.19962 | 0.04976 | 24.11045 | 0 |
| *Lysandra coridon* | 0.39396 | 0.01373 | 28.68919 | 0 |
| *Maculinea arion* | 0.61973 | 0.10498 | 5.90357 | 0 |
| *Maniola jurtina* | 0.08491 | 0.01801 | 4.7155 | 0 |
| *Melanargia galathea* | 0.29715 | 0.04233 | 7.0201 | 0 |
| *Melitaea athalia* | 0.85899 | 0.11303 | 7.59982 | 0 |
| *Pararge aegeria* | -0.23712 | 0.02051 | -11.56351 | 0 |
| *Pieris napi* | 1.14562 | 0.06485 | 17.66557 | 0 |
| *Pieris rapae* | 1.10624 | 0.10164 | 10.88376 | 0 |
| *Plebejus argus* | 0.63581 | 0.02328 | 27.3099 | 0 |
| *Polygonia c-album* | -0.12842 | 0.04515 | -2.84441 | 0.00445 |
| *Polyommatus icarus* | 0.53962 | 0.01949 | 27.69186 | 0 |
| *Pyrgus malvae* | 0.52661 | 0.04862 | 10.83215 | 0 |
| *Pyronia tithonus* | -0.074 | 0.03194 | -2.31716 | 0.0205 |
| temperature : *Aglais urticae* | 0.03222 | 0.00381 | 8.45241 | 0 |
| temperature : *Anthocharis cardamines* | -0.15374 | 0.06522 | -2.35726 | 0.01841 |
| temperature : *Aphantopus hyperantus* | 0.03576 | 0.02106 | 1.69794 | 0.08952 |
| temperature : *Argynnis adippe* | 0.08618 | 0.1103 | 0.78131 | 0.43462 |
| temperature : *Argynnis aglaja* | 0.08293 | 0.04051 | 2.04742 | 0.04062 |
| temperature : *Argynnis paphia* | 0.02061 | 0.06548 | 0.31474 | 0.75296 |
| temperature : *Aricia agestis* | -0.17419 | 0.07069 | -2.46415 | 0.01374 |
| temperature : *Aricia artaxerxes* | 0.02649 | 0.04115 | 0.64363 | 0.51982 |
| temperature : *Callophrys rubi* | -0.0423 | 0.01049 | -4.03218 | 6e-05 |
| temperature : *Celastrina argiolus* | -0.1658 | 0.07495 | -2.21203 | 0.02697 |
| temperature : *Coenonympha pamphilus* | -0.05818 | 0.02136 | -2.72374 | 0.00646 |
| temperature : *Coenonympha tullia* | -0.00404 | 0.01558 | -0.2593 | 0.7954 |
| temperature : *Colias croceus* | -0.06515 | 0.05268 | -1.23666 | 0.21621 |
| temperature : *Cupido minimus* | 0.01513 | 0.05068 | 0.29859 | 0.76526 |
| temperature : *Erebia aethiops* | -0.01915 | 0.01969 | -0.97219 | 0.33096 |
| temperature : *Euphydryas aurinia* | -0.12339 | 0.04346 | -2.8391 | 0.00452 |
| temperature : *Hipparchia semele* | 0.17386 | 0.02009 | 8.65554 | 0 |
| temperature : *Lasiommata megera* | 0.02732 | 0.05619 | 0.48618 | 0.62684 |
| temperature : *Leptidea sinapis* | 0.00741 | 0.06485 | 0.11431 | 0.90899 |
| temperature : *Lycaena phlaeas* | 0.02732 | 0.02986 | 0.91483 | 0.36028 |
| temperature : *Lysandra bellargus* | 0.08316 | 0.0564 | 1.47442 | 0.14037 |

| | | | | |
|---|---|---|---|---|
| temperature : *Lysandra coridon* | 0.01674 | 0.02101 | 0.79679 | 0.42558 |
| temperature : *Maculinea arion* | -0.26533 | 0.14286 | -1.85728 | 0.06327 |
| temperature : *Maniola jurtina* | 0.00358 | 0.02089 | 0.1716 | 0.86375 |
| temperature : *Melanargia galathea* | -0.03017 | 0.05748 | -0.5249 | 0.59965 |
| temperature : *Melitaea athalia* | 0.07 | 0.11186 | 0.62571 | 0.5315 |
| temperature : *Pararge aegeria* | -0.02681 | 0.02408 | -1.11325 | 0.2656 |
| temperature : *Pieris napi* | -0.12772 | 0.03586 | -3.56119 | 0.00037 |
| temperature : *Pieris rapae* | 0.00472 | 0.12526 | 0.03769 | 0.96994 |
| temperature : *Plebejus argus* | 0.04348 | 0.03122 | 1.3927 | 0.16371 |
| temperature : *Polygonia c-album* | -0.00011 | 0.04827 | -0.00225 | 0.9982 |
| temperature : *Polyommatus icarus* | 0.03016 | 0.01595 | 1.89155 | 0.05855 |
| temperature : *Pyrgus malvae* | -0.01501 | 0.07807 | -0.19232 | 0.84749 |
| temperature : *Pyronia tithonus* | 0.00767 | 0.04915 | 0.15604 | 0.876 |
| *Aglais urticae* : year | 0.02072 | 0.01446 | 1.43251 | 0.152 |
| *Anthocharis cardamines* : year | 0.06198 | 0.03968 | 1.562 | 0.11829 |
| *Aphantopus hyperantus* : year | 0.10495 | 0.02837 | 3.69888 | 0.00022 |
| *Argynnis adippe* : year | 0.02551 | 0.08736 | 0.29195 | 0.77033 |
| *Argynnis aglaja* : year | 0.10404 | 0.05319 | 1.9562 | 0.05044 |
| *Argynnis paphia* : year | -0.00964 | 0.03308 | -0.29136 | 0.77078 |
| *Aricia agestis* : year | 0.10848 | 0.05053 | 2.14706 | 0.03179 |
| *Aricia artaxerxes* : year | 0.02603 | 0.04768 | 0.54606 | 0.58503 |
| *Callophrys rubi* : year | 0.0837 | 0.03718 | 2.2511 | 0.02438 |
| *Celastrina argiolus* : year | 0.04839 | 0.03937 | 1.22911 | 0.21903 |
| *Coenonympha pamphilus* : year | -0.02613 | 0.01609 | -1.6244 | 0.10429 |
| *Coenonympha tullia* : year | 0.10832 | 0.02 | 5.41625 | 0 |
| *Colias croceus* : year | 0.03844 | 0.03998 | 0.96151 | 0.3363 |
| *Cupido minimus* : year | -0.05908 | 0.05332 | -1.108 | 0.26786 |
| *Erebia aethiops* : year | 0.0801 | 0.02529 | 3.16772 | 0.00154 |
| *Euphydryas aurinia* : year | 0.1445 | 0.03235 | 4.46638 | 1e-05 |
| *Hipparchia semele* : year | 0.1256 | 0.03801 | 3.30482 | 0.00095 |
| *Lasiommata megera* : year | -0.01156 | 0.03387 | -0.34125 | 0.73292 |
| *Leptidea sinapis* : year | -0.00021 | 0.03305 | -0.00644 | 0.99486 |
| *Lycaena phlaeas* : year | 0.16085 | 0.02498 | 6.4393 | 0 |
| *Lysandra bellargus* : year | 0.07215 | 0.04156 | 1.73605 | 0.08256 |
| *Lysandra coridon* : year | -0.00648 | 0.01413 | -0.45863 | 0.6465 |
| *Maculinea arion* : year | 0.23862 | 0.05737 | 4.15912 | 3e-05 |
| *Maniola jurtina* : year | 0.056 | 0.01888 | 2.96638 | 0.00301 |
| *Melanargia galathea* : year | -0.1592 | 0.02884 | -5.52098 | 0 |
| *Melitaea athalia* : year | 0.18266 | 0.0808 | 2.26075 | 0.02378 |
| *Pararge aegeria* : year | -0.00093 | 0.01873 | -0.04983 | 0.96026 |
| *Pieris napi* : year | -0.01599 | 0.05292 | -0.3022 | 0.7625 |
| *Pieris rapae* : year | 0.11774 | 0.0957 | 1.23025 | 0.21861 |
| *Plebejus argus* : year | 0.10313 | 0.02654 | 3.88517 | 1e-04 |
| *Polygonia c-album* : year | 0.0025 | 0.04219 | 0.05936 | 0.95267 |
| *Polyommatus icarus* : year | 0.06545 | 0.02061 | 3.1764 | 0.00149 |
| *Pyrgus malvae* : year | -0.0529 | 0.04299 | -1.23051 | 0.21851 |
| *Pyronia tithonus* : year | 0.05764 | 0.0259 | 2.22523 | 0.02607 |

***Supp. Table 16: The PC2 quantile regression model results for a τ value of 0.9.***

| Term | Value | Std. Error | t value | P value |
|---|---|---|---|---|
| *Aglais urticae* | 0.27614 | 0.04325 | 6.38521 | 0 |
| *Anthocharis cardamines* | 1.77399 | 0.0569 | 31.17745 | 0 |
| *Aphantopus hyperantus* | 0.9272 | 0.04288 | 21.62364 | 0 |
| *Argynnis adippe* | 2.456 | 0.13245 | 18.54258 | 0 |
| *Argynnis aglaja* | 1.55264 | 0.07593 | 20.44924 | 0 |
| *Argynnis paphia* | 1.19048 | 0.08526 | 13.96238 | 0 |
| *Aricia agestis* | 1.868 | 0.09641 | 19.3753 | 0 |
| *Aricia artaxerxes* | 1.30977 | 0.09517 | 13.7626 | 0 |
| *Callophrys rubi* | 0.96481 | 0.07242 | 13.32181 | 0 |
| *Celastrina argiolus* | 1.73266 | 0.08448 | 20.51002 | 0 |
| *Coenonympha pamphilus* | 0.50314 | 0.0369 | 13.63523 | 0 |
| *Coenonympha tullia* | 0.61512 | 0.06125 | 10.04292 | 0 |
| *Colias croceus* | 0.58383 | 0.11498 | 5.07768 | 0 |
| *Cupido minimus* | 1.48836 | 0.06312 | 23.5797 | 0 |
| *Erebia aethiops* | 0.74491 | 0.13363 | 5.57453 | 0 |
| *Euphydryas aurinia* | 1.27173 | 0.06797 | 18.71019 | 0 |
| *Hipparchia semele* | 1.65743 | 0.06674 | 24.835 | 0 |
| *Lasiommata megera* | 1.16137 | 0.06642 | 17.48417 | 0 |
| *Leptidea sinapis* | 3.28892 | 0.03467 | 94.86194 | 0 |
| *Lycaena phlaeas* | 1.1191 | 0.04462 | 25.08141 | 0 |
| *Lysandra bellargus* | 2.33325 | 0.04611 | 50.5984 | 0 |
| *Lysandra coridon* | 1.31799 | 0.02099 | 62.78526 | 0 |
| *Maculinea arion* | 1.82845 | 0.06703 | 27.2775 | 0 |
| *Maniola jurtina* | 0.84713 | 0.03648 | 23.22279 | 0 |
| *Melanargia galathea* | 1.4652 | 0.09076 | 16.14356 | 0 |
| *Melitaea athalia* | 2.1447 | 0.12234 | 17.53006 | 0 |
| *Pararge aegeria* | 0.3625 | 0.06823 | 5.31287 | 0 |
| *Pieris napi* | 2.14971 | 0.04318 | 49.78929 | 0 |
| *Pieris rapae* | 2.32182 | 0.06882 | 33.73678 | 0 |
| *Plebejus argus* | 1.75799 | 0.03561 | 49.36807 | 0 |
| *Polygonia c-album* | 0.41418 | 0.02114 | 19.59262 | 0 |
| *Polyommatus icarus* | 1.51956 | 0.02585 | 58.77317 | 0 |
| *Pyrgus malvae* | 1.3883 | 0.05995 | 23.15874 | 0 |
| *Pyronia tithonus* | 0.681 | 0.0421 | 16.17718 | 0 |
| temperature : *Aglais urticae* | 0.01372 | 0.04525 | 0.30322 | 0.76173 |
| temperature : *Anthocharis cardamines* | -0.19731 | 0.07048 | -2.79933 | 0.00512 |
| temperature : *Aphantopus hyperantus* | 0.10938 | 0.04362 | 2.50774 | 0.01215 |
| temperature : *Argynnis adippe* | 0.05957 | 0.18964 | 0.31412 | 0.75343 |
| temperature : *Argynnis aglaja* | 0.11018 | 0.04936 | 2.23234 | 0.02559 |
| temperature : *Argynnis paphia* | -0.0825 | 0.1506 | -0.5478 | 0.58383 |
| temperature : *Aricia agestis* | 0.03175 | 0.13479 | 0.23551 | 0.81381 |
| temperature : *Aricia artaxerxes* | 0.00727 | 0.03472 | 0.20941 | 0.83413 |
| temperature : *Callophrys rubi* | -0.00644 | 0.08566 | -0.07516 | 0.94009 |
| temperature : *Celastrina argiolus* | -0.13724 | 0.1396 | -0.98309 | 0.32556 |
| temperature : *Coenonympha pamphilus* | -0.14358 | 0.04571 | -3.14087 | 0.00168 |
| temperature : *Coenonympha tullia* | -0.0589 | 0.03021 | -1.94979 | 0.0512 |
| temperature : *Colias croceus* | -0.02757 | 0.10198 | -0.27035 | 0.78689 |
| temperature : *Cupido minimus* | -0.24342 | 0.0753 | -3.23249 | 0.00123 |
| temperature : *Erebia aethiops* | -0.01314 | 0.03645 | -0.36056 | 0.71843 |
| temperature : *Euphydryas aurinia* | -0.13811 | 0.07382 | -1.87095 | 0.06135 |
| temperature : *Hipparchia semele* | 0.18717 | 0.07579 | 2.4696 | 0.01353 |
| temperature : *Lasiommata megera* | 0.09734 | 0.08854 | 1.09939 | 0.2716 |
| temperature : *Leptidea sinapis* | -0.02377 | 0.05525 | -0.43024 | 0.66702 |
| temperature : *Lycaena phlaeas* | 0.06302 | 0.02775 | 2.2705 | 0.02318 |
| temperature : *Lysandra bellargus* | 0.08507 | 0.05414 | 1.57127 | 0.11612 |

| | | | | |
|---|---|---|---|---|
| temperature : *Lysandra coridon* | 0.02214 | 0.03255 | 0.68037 | 0.49627 |
| temperature : *Maculinea arion* | -0.30684 | 0.27026 | -1.13533 | 0.25624 |
| temperature : *Maniola jurtina* | -0.03228 | 0.04724 | -0.68331 | 0.49441 |
| temperature : *Melanargia galathea* | 0.25341 | 0.1372 | 1.84702 | 0.06475 |
| temperature : *Melitaea athalia* | 0.11226 | 0.15257 | 0.73575 | 0.46189 |
| temperature : *Pararge aegeria* | -0.05203 | 0.08029 | -0.64801 | 0.51698 |
| temperature : *Pieris napi* | -0.04261 | 0.02468 | -1.72637 | 0.08428 |
| temperature : *Pieris rapae* | 0.06109 | 0.04683 | 1.30465 | 0.19202 |
| temperature : *Plebejus argus* | 0.00839 | 0.04673 | 0.17954 | 0.85751 |
| temperature : *Polygonia c-album* | 0.05196 | 0.02411 | 2.15492 | 0.03117 |
| temperature : *Polyommatus icarus* | 0.09826 | 0.01469 | 6.68828 | 0 |
| temperature : *Pyrgus malvae* | 0.02425 | 0.10348 | 0.23437 | 0.8147 |
| temperature : *Pyronia tithonus* | -0.11342 | 0.05072 | -2.23632 | 0.02533 |
| *Aglais urticae* : year | 0.10409 | 0.03944 | 2.63906 | 0.00831 |
| *Anthocharis cardamines* : year | -0.00601 | 0.03827 | -0.15713 | 0.87514 |
| *Aphantopus hyperantus* : year | 0.12643 | 0.04386 | 2.88272 | 0.00394 |
| *Argynnis adippe* : year | 0.08377 | 0.13814 | 0.60642 | 0.54424 |
| *Argynnis aglaja* : year | 0.07198 | 0.05553 | 1.29621 | 0.1949 |
| *Argynnis paphia* : year | 0.15909 | 0.07666 | 2.07516 | 0.03797 |
| *Aricia agestis* : year | 0.11428 | 0.08922 | 1.28089 | 0.20024 |
| *Aricia artaxerxes* : year | 0.15381 | 0.06002 | 2.56279 | 0.01038 |
| *Callophrys rubi* : year | 0.1147 | 0.0329 | 3.48604 | 0.00049 |
| *Celastrina argiolus* : year | 0.00451 | 0.0759 | 0.05937 | 0.95266 |
| *Coenonympha pamphilus* : year | -0.02591 | 0.0331 | -0.78282 | 0.43373 |
| *Coenonympha tullia* : year | 0.22947 | 0.04618 | 4.96934 | 0 |
| *Colias croceus* : year | -0.04481 | 0.0791 | -0.56655 | 0.57102 |
| *Cupido minimus* : year | 0.05533 | 0.06396 | 0.86513 | 0.38697 |
| *Erebia aethiops* : year | 0.26769 | 0.0433 | 6.18286 | 0 |
| *Euphydryas aurinia* : year | 0.24206 | 0.06338 | 3.81909 | 0.00013 |
| *Hipparchia semele* : year | 0.10625 | 0.05336 | 1.99125 | 0.04646 |
| *Lasiommata megera* : year | 0.02086 | 0.06241 | 0.33423 | 0.73821 |
| *Leptidea sinapis* : year | 0.00238 | 0.01749 | 0.13609 | 0.89175 |
| *Lycaena phlaeas* : year | 0.23054 | 0.04494 | 5.13027 | 0 |
| *Lysandra bellargus* : year | 0.08369 | 0.03789 | 2.20903 | 0.02717 |
| *Lysandra coridon* : year | 0.00654 | 0.02282 | 0.28645 | 0.77453 |
| *Maculinea arion* : year | 0.29478 | 0.05064 | 5.82126 | 0 |
| *Maniola jurtina* : year | 0.16223 | 0.03849 | 4.21514 | 2e-05 |
| *Melanargia galathea* : year | -0.16759 | 0.06711 | -2.4974 | 0.01251 |
| *Melitaea athalia* : year | 0.22304 | 0.09802 | 2.2755 | 0.02288 |
| *Pararge aegeria* : year | 0.05893 | 0.05215 | 1.12988 | 0.25853 |
| *Pieris napi* : year | -0.0276 | 0.03394 | -0.81316 | 0.41613 |
| *Pieris rapae* : year | -0.01623 | 0.06408 | -0.25321 | 0.80011 |
| *Plebejus argus* : year | 0.15293 | 0.04049 | 3.77673 | 0.00016 |
| *Polygonia c-album* : year | 0.07018 | 0.02075 | 3.38144 | 0.00072 |
| *Polyommatus icarus* : year | 0.07876 | 0.02458 | 3.20402 | 0.00136 |
| *Pyrgus malvae* : year | -0.07312 | 0.05703 | -1.28203 | 0.19983 |
| *Pyronia tithonus* : year | 0.02448 | 0.02825 | 0.86666 | 0.38613 |
